# Diverging Parkinson’s Disease Pathology between patient-derived *GBA^N370S^, LRRK2^G2019S^* and engineered *SNCA^A53T^* iPSC-derived Dopaminergic Neurons

**DOI:** 10.1101/2023.01.06.521264

**Authors:** Ali Fathi, Kiranmayee Bakshy, Lida Zieghami, Rebecca Fiene, Robert Bradley, Sarah Dickerson, Coby Carlson, Scott Schachtele, Jing Liu

## Abstract

Multiple neurodegenerative disorders, including Parkinson’s disease (PD) and Alzheimer’s disease-associated dementia (ADAD), are linked with dopaminergic (DA) neuron death and a resulting reduction in dopamine levels in the brain. DA neuron degeneration and the risk of developing PD is connected to genetic mutations affiliated with lysosomal function and protein degradation. Accessible human cellular models for PD-relevant genetic mutations are needed to investigate mechanisms of DA cell death and define points of therapeutic intervention. Human induced pluripotent stem cell (iPSC)-derived midbrain DA neurons offer a developmentally and physiologically relevant *in vitro* model for investigating PD pathogenic mechanisms across genetic backgrounds. In this study, we generated DA neurons using iPSCs from two clinically diagnosed PD patients, one harboring an inherited *GBA*^N370S^ mutation and the other a mutation in *LRRK2*^G2019S^ and compared pathophysiology against DA neurons from genetically engineered *SNCA*^A53T^ iPSCs and its isogenic apparently healthy normal (AHN) iPSCs. Our results present a novel phenotype for *GBA*^N370S^ and *LRRK2*^G2019S^ derived DA neurons, showing that they produced and released significantly more dopamine compared to the AHN and *SNCA*^A53T^ mutant DA neurons. All mutant DA neurons developed deficient glucocerebrosidase (GCase) activity, increased mitochondrial stress, aberrant neuronal activity patterns, and increased α-synuclein accumulation. Together these data suggest potentially divergent origins of PD pathogenesis in *GBA*^N370S^ and *LRRK2*^G2019S^ DA neurons. In addition, compound screening confirmed that GCase modulators can rescue enzyme activity and impact neural activity across all DA mutant neurons, to varying degrees. These data demonstrate unique *in vitro* phenotypes associated with PD and suggest a diversity of underlying mechanisms across different genetic backgrounds. Together, the cell lines used in this study present a valuable tool for new therapeutic discovery.

## Introduction

Cell models are simplified systems that recapitulate characteristics of cells in tissues of the body and are commonly used as an accessible, *in vitro* experimental surrogate for human studies and animal models. Human induced pluripotent stem cell (iPSC)-derived models are accelerating the study of human diseases because iPSCs preserve the genetic predisposition of the donor cells and the terminally differentiated progeny can recapitulate disease phenotypes in a dish. Disease modeling with iPSC is a broadly adopted approach to elucidate complex mechanisms of neurodegenerative diseases.

When applied to monogenic early onset diseases such as Huntington’s disease ^1^, spinal muscular atrophy ^2,3^, Thalassemia ^4^, and cystic fibrosis ^5^, iPSC-derived cell models have successfully reproduced disease phenotypes and enabled mechanistic studies. However, late onset multifactorial diseases such as Parkinson’s disease (PD) and Alzheimer’s disease (AD) ^6–8^ continue to face challenges in model validation and recapitulation of disease, including properly encompassing variations in genetic neurodegenerative disease etiology, integration of iPSC donors from diverse backgrounds, and controlling for variations in differentiation processes.

Previously reported studies using dopaminergic (DA) neurons derived from iPSC, either innately harboring PD-associated genetic mutations or genetically engineered to contain PD-relevant mutations in leucine-rich repeat kinase 2 (LRRK2), Glucocerebrosidase (GBA), and α-synuclein (α-Syn) (SNCA) mutations, have shown inconsistent disease phenotypes compared to healthy or isogenic controls. Notably, differences in α-Syn protein aggregation ^9,10^, mitochondrial dysfunction ^11–14^ and DNA damage ^15,16^ have been observed. In addition, no α-Syn fibrils or Lewy bodies were reported in DA neurons derived from disease iPSCs without using α-Syn seeds ^17^, and the reported disease phenotypes were not consistent among different studies and mutations.

Despite varying phenotypes across patient-derived iPSC models of PD, their application in the pursuit of biomarkers or novel therapeutic compounds is still ongoing. Discovery of biomarkers that facilitate early diagnosis of PD is needed to improve therapeutic strategies. Current PD treatments are often started following a late-stage PD diagnosis, which limits their therapeutic efficacy and patient benefit because the target DA population is already significantly depleted. Elucidating early mechanisms or biomarkers for PD will open avenues for DA neuron preservation by providing a larger therapeutic window for administration of neuroprotective drugs. By nature, iPSC-derived DA neurons, which exhibit developmentally young features, provide a model for early PD disease onset before cell death. Developing a more comprehensive understanding of PD by utilizing a genetically diverse pool of clinically diagnosed patients or genetically engineered iPSCs could expedite methods for early diagnosis and help discover alternative interventions to stop disease progression, both towards a general PD population or more targeted to a specific PD-associated genetic mutation.

In this study, iPSC derived from *LRRK2*^G2019S^ and *GBA*^N370S^ clinically diagnosed PD patient donor and iPSC engineered to contain the highly penetrant *SNCA*^A53T^ alpha-synuclein risk-associated genetic mutation were differentiated into pure populations of DA neurons and evaluated for disease related phenotypes. We discovered that patient-derived DA neurons showed similar onset of disease phenotypes previously reported for the engineered *SNCA*^A53T^ DA neurons. In addition, we found novel differences in dopamine synthesis, degradation, and release among PD mutant DA neurons. Together these data suggest a divergent origin of common PD disease phenotypes in iPSC-derived models and provide a more comprehensive picture of potential underlying causes of PD in human DA neurons.

## Methods

### Cell lines and DA neurons culture

All iPSC lines including AHN iPSCs from donor line 01279, donor line 11344 with GBA^N370S^ mutation and donor lines 11299 and 11305 with LRRK2^G2019S^ mutation were developed by FUJIFILM Cellular Dynamics, Inc. (FCDI) (https://hpscreg.eu/cell-line) and all protocols for derivation of source material approved by IRB committee at MJFF/PPMI. Human iPSCs were generated via episomal reprogramming of PBMC from clinically diagnosed PD patients harboring either the GBA^N370S^ or LRRK2^G2019S^ mutation and volunteers provided written IRB approved informed consent form when samples collected. Source material for iPSC lines were obtained from the Parkinson’s Progression Markers Initiative (PPMI) and the Golub Capital iPSC PPMI Sub-study (http://www.ppmi-info.org). PPMI is a public-private partnership funded by the Michael J. Fox Foundation for Parkinson’s research and other funding partners. Pluripotency factors used for reprogramming were Oct4, Sox2, Nanog, Lin28, Klf4, L-Myc and SV40 large T-antigen. *SNCA*^A53T^ heterozygous iPSCs were genetically engineered from apparently healthy normal (AHN) iPSCs (donor 01279). Mutation insertion was accomplished using nuclease-mediated homologous recombination and a donor oligo SJD 14–132:

(GTTTTACAATTTCATAGGAATCTTGAATACTGGGCCACACTAATCACTAGATA CTTTAAATATCATCTTTGGATATAAGCACAATGGAGCTTACCTGTgGtgACACCATGCACCACT CCCTCCTTGGTTTTGGAGCCTACAAAAACAAATTCAAGACATAAGTCTCAAGCTAGCCTTAA ATTGCTGATTAGCTAGTTTTC). The resulting iPSC line contained SNP rs104893877 where amino acid 53 was changed from alanine to threonine (A÷T), resulting in the *SNCA*^A53T^ heterozygous iPSC line. Clones used for DA neuron differentiation from each iPSC line were confirmed for the presence of mutation, expression of pluripotency markers and normal karyotype.

DA neurons were differentiated using previous published method ^18^ modified by FCDI for large scale manufacturing and consistency across cell lines. Cryopreserved AHN and *SNCA*^A53T^ (donor 01279), *GBA*^N370S^ (donor 11344) and *LRRK2^G2019S^* (donors 11299 and 11305) iPSC-derived DA neurons were thawed and maintained in culture using Complete Maintenance Medium (Neural Base Medium 1 supplemented with iCell Nervous System Supplement and iCell Neural Supplement B) prepared according to the iCell DopaNeurons User’s Guide (Document ID: X1003).

### Immunofluorescent (IF) staining and microscopy

Human iPSC-derived DA neurons were seeded at 6.4×10^4^ per well in the 96 well Greiner imaging plates. Cells were fixed at day 3 and 14 post thaw for 20 minutes with 4% paraformaldehyde in Phosphate Buffered Saline (PBS). Samples were washed with FACS buffer (2% fetal bovine serum in PBS) and permeabilized using 0.1% Triton X-100 (Sigma-Aldrich, T-8787) with 1% goat or donkey serum in PBS buffer and blocked with 4% donkey serum for one hour. Primary antibodies were diluted in 4% donkey serum and 0.1% Tween 20 and applied to samples overnight at 4 °C. Samples were washed with FACS buffer three times (5 min each), incubated with fluorescein-conjugated secondary antibodies for one hour at room temperature, and counterstained with Hoechst for 20 minutes followed by washing with FACS buffer. Samples were imaged on ImageXpress (Molecular Devices).

**Table 1:** Primary antibodies used for immunostaining of DA neurons.

| Antibody | Species | Catalog No. | Company | Dilution |
| --- | --- | --- | --- | --- |
| **FOXA2** | Rabbit | 8186 | Cell Signaling | 2,000 |
| **FOXA2** | Mice | Ab60721 | Abcam | 10,000 |
| **TH** | Mice | T2928 | Sigma | 10,000 |
| **LMX1** | Rabbit | Ab10533 | Millipore | 5,000 |
| **MAP2** | Mice | M2320 | Sigma | 2,000 |

### Flow cytometry assay

Human iPSC-derived DA neurons were cultured for 3 and 14 days prior to dissociation using Accutase for 15 minutes at 37 °C. Harvested cells were stained with LIVE/DEAD™ fixable red dead cell stain kit (ThermoFisher) and then fixed for 20 minutes with 4% paraformaldehyde in PBS. Samples were washed and permeabilized using flow permeabilization buffer (0.1% Triton X100 + 2% fetal bovine serum) and stained overnight using primary antibodies. Samples were washed with FACS buffer and incubated with fluorescein-conjugated secondary antibodies for one hour at room temperature. Samples were washed, run, and analyzed using the BD Accuri™ C6 Plus Flow Cytometer and software (BD Biosciences).

### Dopamine release assay

Human iPSC-derived DA neurons were seeded at a density of 3×10^6^ cells per well in 12 well plates and maintained in culture using Complete Maintenance Medium. On day 22, Complete Maintenance Medium was aspirated, cells were washed for 10 minutes using Hank’s Balanced Salt Solution (HBSS, -Mg/-Ca) followed by incubation at 37 °C for 30 minutes in 300 μL of fresh Hank’s Balanced Salt Solution (HBSS, -Mg/-Ca). After 30 minutes of incubation, supernatants were collected as pre-stimulation samples. KCl (56 mM) in HBSS was then added to each well and incubated at 37 °C for another 30 minutes. Supernatants were collected at the end of the incubation as post-stimulation samples. EDTA (1 mM final) and sodium metabisulfite (4 mM final) were added to samples prior to cryopreservation at −80 °C. To determine dopamine concentration, samples were assayed using a competitive Dopamine ELISA Assay Kit (Eagle Biosciences) following the kit instructions. Three technical replicates were performed per biological sample. The standard curve was graphed in GraphPad PRISM and all other values were interpolated with non-linear regression using the log(agonist) vs. response - variable slope (four parameters) model.

### RNA-seq procedure and analysis

iPSC-derived DA neurons were thawed and maintained according to the iCell DopaNeurons User’s Guide. In brief, 2×10^6^ cells were plated onto poly-l-ornithine/laminin coated 6-well tissue culture plates and cultured for ≥14 days. At 14 days in culture, medium was removed, and cells were lysed directly in the well using 400 μL of Qiagen RLT Plus lysis buffer. Duplicate samples/lot of ~2×10^6^ cells each are lysed and stored at −80°C until RNA extraction is performed. The QiaSymphony is used to extract the RNA. RNA concentration should be greater than 20 ng/μL and the ratio of absorbance (OD 260/280) should be greater than 2.0 to ensure acceptable purity of the sample.

cDNA library preparation was performed according to manufacturer’s recommendation (Illumina (TS)). Briefly, samples were tailed and ligated with modified oligos, size selected, and amplified. All the libraries were then tested prior to sequencing using 1) the Agilent BioAnalyzer for sizing and quality assessment of the library and 2) qPCR for measuring the concentration of the library and presence of Illumina anchor sequences.

RNA sequencing of iPSC-derived DA neurons was performed following a modified protocol ^19^. Briefly RNA sequencing was performed on Illumina NovaSeq 6000 platform (Illumina Inc., San Diego, CA) at Novogene, targeting at least 20 million paired end reads with a read length of 150 bp. FastQC v11 was used to check the quality of reads generated. High quality reads were mapped to the Ensembl (GRCh38.p10) Homo sapiens genome using splice-aware alignment program, HISAT2, v.2.1.0. Differential gene expression analysis between the wild-type, patient-derived, and engineered DA neuron samples was carried out using default settings in the Cuffdiff, v.2.2.1 program. Custom Perl scripts were used to convert the Cuffdiff output files to compatible text files for visualization in TIBCO Spotfire software v.11.0.0. All the plots were generated using TIBCO Spotfire.

### MitoSOX staining

MitoSOX imaging assay was performed using MitoSOX™ Red Mitochondrial Superoxide Indicator (Thermo Fisher, M36008). MitoSOX red (5 μM) was added to the cells and incubated for 30 minutes at 37 °C. After the removal of MitoSOX, Calcein AM (Thermo Fisher, C1430) live ready reagent was added to the neurons and imaged using the Incucyte S3 for 3 hours every 30 minutes. Rotenone (2 μM), a mitochondrial toxin, was used as a positive control condition and applied to the cells for 30 mins at 37 °C followed by a medium wash prior to MitoSOX staining.

### Western blotting

Human iPSC-derived DA neurons were seeded at high density 2× 10^6^ per well of 12-well plates. At day 14 cells were washed with PBS and lysed on ice using RIPA Lysis Buffer (Cell Signaling) supplemented with Halt Protease and Phosphatase Inhibitor Cocktail (Thermo Fisher). Samples were centrifuged at 18,000 rpm for 20 minutes at 4 ^o^C. Total protein concentrations were measured using Pierce BCA Protein Assay (Thermo Fisher). Protein samples (1 μg) were run on 25-capillary cartridge 12-230 kDa (ProteinSimple) using the WES instrument. Signals were visualized and the area under the curve quantified and analyzed using Compass for SW (V 3.1.7). The following primary antibodies were used: Tyrosine Hydroxylase (1:250, Sigma T2928), GBA (1:100, Bio-Techne, MAB4710), DDC (1:500, Novus Biologicals, NB300-174), and MAPT (1:1000, Millipore, MAB361).

### Neuronal activity using micro electrode array (MEA)

Human iPSC-derived DA neurons were plated on a 48-well Classic MEA plate (Axion BioSystems) at a cell density of 1.2×10^5^ cells per well and maintained in culture using Complete BrainPhys™ (Stem Cell Technologies). Culture Medium prepared as directed in the iCell DopaNeurons MEA Application Protocol (https://www.fujifilmcdi.com/). MEA recordings were taken every few days to monitor the development of synchronously bursting neuronal networks in culture over time. MEA signals were recorded using a Maestro MEA System (Axion BioSystems). Data acquisition was managed with AxIS software (Axion Biosystems). All channels were sampled at the same time with a gain of 1200x and a sampling frequency of 12.5 kHz/channel with a 200-5000 Hz band-pass filter. Prior to recording, MEA plates were allowed to equilibrate for 10 min on the instrument. The raw data files were recorded, converted to .spk files, and subsequently analyzed using the Neural Metric Tool software (Axion BioSystems). Burst detection was performed using the “envelope method”.

### Alpha-Synuclein protein assays (Thioflavin T staining and MSD assay)

iPSC-derived DA neurons were plated at the density of 6.4×10^4^ cells per well in a 96-well plate for Thioflavin T (Th-T) staining. At day 21 post thaw, cells were treated with 4 μg/ml of Recombinant Mouse α-Syn Protein Aggregate (Active) (Abcam, ab246002) for periods of 24, 48, and 72 hours. Cells were then stained with 10 μM Th-T (Abcam, ab120751) in HBSS and 1 μM YOYO-3 iodide (Thermofisher, Y3606) in the culture media. Following incubation for 30 min, cells were washed with warm HBSS and imaged using the Incucyte 20X objective. Th-T signal intensity was quantified in each neuron and across conditions using the Incucyte analysis software.

The U-PLEX Human α-Synuclein Kit (Meso Scale Diagnostics, MSD-K151WKK) was used for quantification of α-Syn protein expression in iPSC-derived DA neurons. For this assay, cells were not pretreated with active α-Syn protein aggregate. At day 21 and 42 post plating, cells were lysed directly into the RIPA Lysis Buffer supplemented with Halt Protease and Phosphatase Inhibitor Cocktail (Thermo Fisher). Samples were centrifuged at 18,000 rpm for 20 min at 4 °C. Total protein concentrations were measured using Pierce BCA Protein Assay (Thermo Fisher). Samples were run at 1 μg and 0.2 μg total protein with standards and calibrators on the same plate. The calibration curve used to calculate analyte concentrations was established by fitting the signals from the calibrators to a 4-parameter logistic (or sigmoidal dose-response) model with a 1/Y2 weighting. Analyte concentrations were determined from the ECL signals by back fitting to the calibration curve. The calculations to establish calibration curves and determine concentrations were carried out using the MSD DISCOVERY WORKBENCH® analysis software.

### GCase activity

Human iPSC-derived DA neurons were plated in 12 well plates at 2×10^6^ cells per well. At 14 days post thaw, cells were washed with PBS and lysed on ice using 200 μL of assay buffer (0.25% (v/v) Triton X-100, 0.25% (w/v) Taurocholic acid (Sigma-Aldrich, T9034), 1 mM EDTA, 50 mM citric acid, 200 mM Na2HPO4, pH 5.4). Cell lysates were frozen/thawed twice and incubated on ice for 30 min, then centrifuged at 20,000 x g for 20 min at 4 °C. Total protein concentrations were measured using Pierce BCA Protein Assay. Protein samples (5 μg) were used to determine GCase activity upon addition of 10 mM 4-Methylumbelliferyl ß-glucophyranoside (4-MU, Sigma-Aldrich, # M3633) in 1% BSA with or without 2 mM conduritol-b-epoxide (CBE, GCase inhibitor) in 50 μL of total volume. Recombinant Human Glucosylceramidase/GBA (rhGBA) (R&D, 7410-GHB-020) was used as positive control.

Samples were incubated for 40 min at 37 °C and the reaction stopped by adding an equal volume of 1M glycine at pH 12.5. Samples (100 μL) were loaded into white 96-well plates (Nunc, # 136101) and fluorescence (Ex = 355 nm, Em = 460 nm, 0.1 sec) signals were measured in the CLARIOstar plate reader (BMG Labtech). Corrected signal of each condition was calculated by subtracting the signal of CBE treated lysates from the corresponding signal of non-CBE treated lysates. For high throughput compound screening, DA neurons were cultured at 2×10^5^ cells per well in 96-well plates. Cells were dosed with compounds (Table S3) at three different concentrations (1X, 10X and 100X) and incubated for 3 days or 14 days (chronic treatment). At respective days post dosing, cells were harvested using 100 μL of assay buffer and GCase activity was determined using the procedure previously described. Each sample used 5 μg of protein lysate and net GCase activity was graphed in comparison to the 0.1% DMSO treated cells as the negative control.

### Calcium imaging

For calcium imaging experiments, DA neurons were cultured as spheroids (5×10^4^ cells/well) in 384 U-bottom ULA plates (S-Bio) and fed with Complete BrainPhys medium (BrainPhys basal media with iCell Neural Supplement B and iCell Nervous System Supplement). On day 7 post plating 10 μM Ambroxol (ABX) was added to some of the wells and incubate for 7 days. On day 14, 21 and 28 post plating cells were loaded with 1X EarlyTOX from the EarlyTOX Cardiotoxicity Kit (Molecular Devices) and incubated for 2 hours at 37 °C. Calcium oscillation was captured using FDSS/μCELL Functional Drug Screening System (Hamamatsu) for 10-20 min. Waveform software was used for quantification of calcium oscillation and data analyzed using GraphPad Prism.

### Statistical Analysis

Graphpad (Prism 9) was used for statistical analysis. Unpaired Student’s t-test or One Way ANOVA test with Dunnet’s multiple comparison or Tukey’s post hoc test was performed.

## Results

### *GBA*^N370S^, *LRRK2*^G2019S^ and *SNCA*^A53T^ iPSCs generated comparable DA neurons to the healthy normal iPSCs

Five iPSC lines, including *GBA*^N370S^, *LRRK2*^G2019S^ (2 lines), *SNCA*^A53T^ and apparently healthy normal (AHN), were differentiated into DA neurons following a modified protocol ^18^ (Figure 1A). All five lines generated highly pure DA neurons as shown by the majority of cells expressing tyrosine hydroxylase (TH) and FOXA2 at the end of the differentiation (Figure 1B) and prior to cryopreservation. AHN and PD mutant DA neurons cultured for 14 days post thaw continued to express TH, which was found to be co-expressed with DA specific transcription factors, FOXA2 and LMX1, indicating bona fide midbrain DA neurons present in the culture (Figure 1C). Furthermore, analysis of DA markers across multiple batches differentiations for both AHN and PD mutant DA neurons showed a consistently high percentage of neurons expressing MAP2 (general neuronal marker), TH, and FOXA2. No significant differences were detected in the percentage of the TH-positive, FOXA2-positive, and MAP2-positive neurons across all lines (Figure 1D).

**Figure 1:**
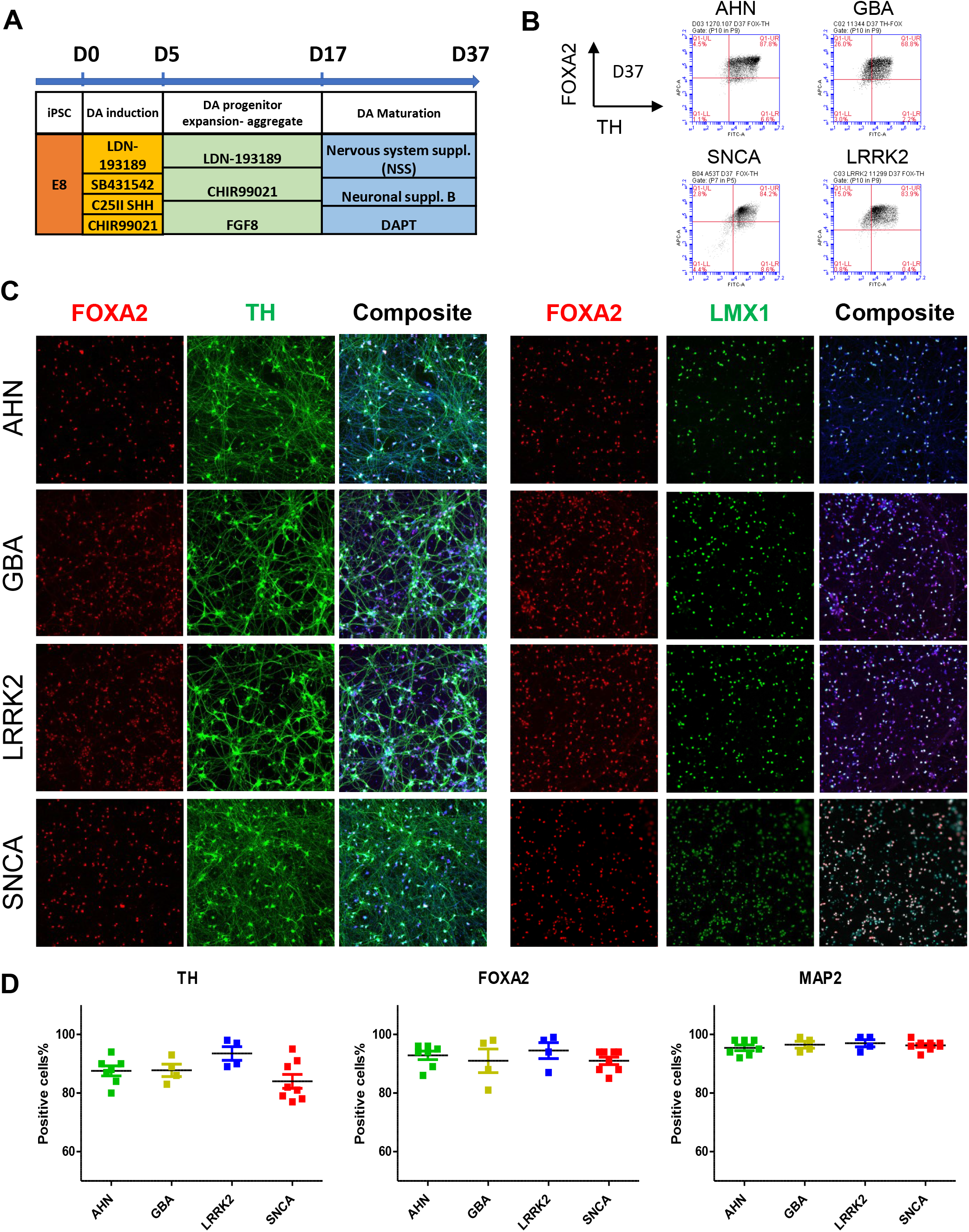
Differentiation and characterization of iPSC derived DA neurons. **(A)** Schematic of differentiation protocol for DA neurons consists of 4 stages and each stage requires different media and supplements. (B) At the end of differentiation (day 37) and prior to cell cryopreservation, all four cell lines were evaluated for expression of TH and FOXA2 expression by flow cytometry. (C) Cryopreserved DA neurons were thawed and cultured for 14 days prior to immunocytochemistry. Staining confirmed the identity of midbrain DA neurons with the majority of neurons from all four iPSC lines showing double positive for FOXA2 and TH (C; left panel) and for FOXA2 and LMX1 (C; right panel). Multiple lots of DA neurons from AHN and mutant iPSCs were analyzed for marker expression 3 days post thaw. (D) Quantified flow cytometry analysis of the TH, FOXA2 and MAP2 protein expression showed consistent marker expression across the multiple lots each line.

### RNAseq gene expression of the GBA^N370S^, *LRRK2*^G2019S^ and *SNCA*^A53T^ iPSC derived DA neurons

To further compare AHN, engineered SNCA^A53T^, and patient-derived LRRK2^G2019S^ and GBA^N370S^ iPSC-derived DA neurons, cells were harvested at day 14 post thaw, processed, and analyzed for gene expression using RNAseq. Log RPKM (reads per kilobase of exon per million reads mapped) of all transcripts showed a close correlation between the AHN and each mutant line derived DA neurons (Figure 2A). Differential expression analysis (DEA) indicated that 297 transcripts in the *GBA^N370S^,* 450 transcripts in the *LRRK2* ^G2019S^, and 441 transcripts in the *SNCA*^A53T^ were differentially regulated compared to the control AHN cells (≥2-fold change and p<0.001, Figure 2B and Table S1). Analysis of DEA transcripts annotated by top pathways that each set (mutation) was enriched, identified three pathways commonly dysregulated across mutant DA neurons compared to the AHN neurons, including: (1) transcripts involved in calcium ion binding (31 transcripts) in *LRRK2*^G2019S^ and then *SNCA*^A53T^ (28 transcripts). (2) transcripts for extracellular matrix components (including genes coding collagens and mucins), and (3) transcripts with neuropeptide hormone activity (including *PENK, VIP, GRP, CRH* and *NTS).*

**Figure 2:**
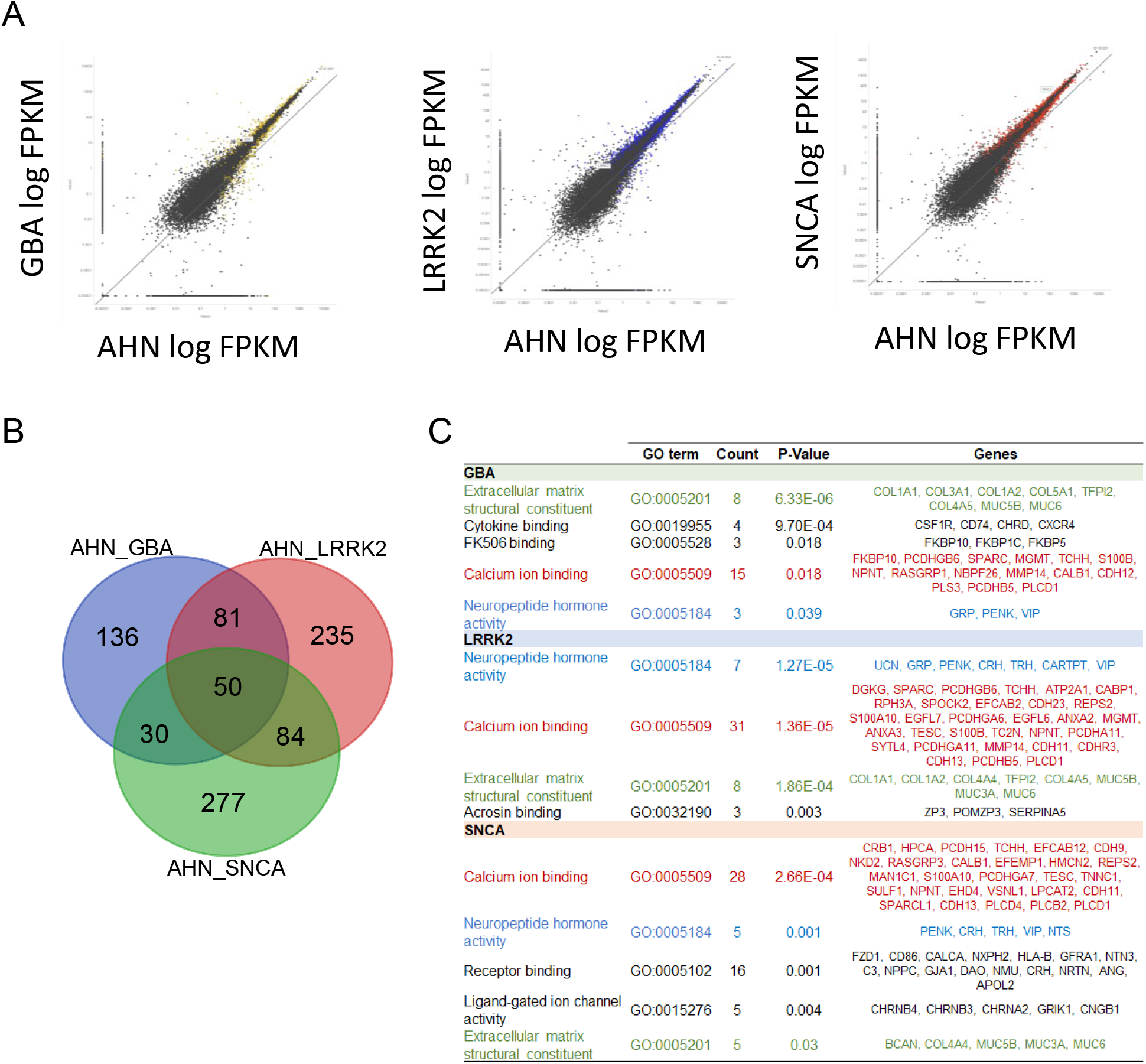
RNAseq data analysis at 14 days post thaw. **(A)** Transcripts of the mutant DA neurons were plotted against the AHN neurons and showed a high correlation in gene expression profiles. Transcripts that showed significantly different expression between lines are highlighted with color. (B) Differential gene expression analysis was carried out for each mutant DA neurons compared to the AHN DA neurons. The number of differential transcripts is summarized in the Venn diagram. (C) Gene set enrichment and pathway analysis for differentially regulated transcripts in mutant DA neurons compared to the AHN samples in DAVID database. Pathways highlighted in color were common modulated across mutant DA neurons.

Neuronal toxicity derived from increased DA levels, DA oxidation and its reactive catabolites, is recognized as one of the major causes of oxidative stress in PD ^20,21^. To further investigate the effects of these PD mutations on DA neuron gene expression, we compared mRNA expression for 132 genes involved in dopamine synthesis and release pathways. Surprisingly, few genes showed significant changes between AHN and PD mutants. However, most of the differentially expressed genes (DEGs) identified were commonly observed across GBA^N370S^, LRRK2^G2019S^ and SNCA^A53T^ DA neurons (Table S2). *DDC, CALY, CALM2, GNAS, FOS* and *GNB1* transcripts were upregulated in all mutant neurons compared to the AHN, and *CALM1* transcript (Calmodulin 1) was downregulated in the mutant neurons. Divergent DEG patterns were also observed between mutant cell lines, including *TH, MAOA, VMAT1,2 and COMT* which was upregulated in the GBA^N370S^ and *LRRK2*^G2019S^ mutants, but not in *SNCA*^A53T^ cells. On the contrary, *ATF4* and *GNAO1* were transcripts in the DA pathways that were differentially expressed in the *SNCA*^A53T^ neurons but not significant in the GBA^N370S^ and *LRRK2*^G2019S^ mutant neurons (Table S2).

### Mutant DA neurons recapitulated in vivo α-Syn phenotype

Alpha-synuclein aggregation is a major pathological hallmark of PD developed in patient brains, therefore we investigated if this is recapitulated in iPSC-derived GBA^N370S^, LRRK2^G2019S^ and SNCA^A53T^ DA neurons *in vitro.* To answer this question, we used active recombinant mouse α-Syn protein aggregate as seeds for inducing α-Syn aggregation in cultured AHN and mutant DA neurons and visualized protein aggregation using Thioflavin T (Th-T) staining ^22^.

Day 21 cultured GBA^N370S^, LRRK2^G2019S^ and SNCA^A53T^ DA neurons, seeded with active recombinant mouse α-Syn protein for 24 hours, displayed higher average TH-T signal per neuron. Analysis of Th-T signal intensity within individual neurons showed that the Th-T distribution in mutant neurons skewed toward a higher intensity suggesting increased aggregation compared to AHN DA neurons (Figure 3A-B). In addition, the average mean intensity of stained neurons following α-Syn seeding, showed a significant increase in the signal intensity in all mutant neurons compared to the AHN neurons (Figure 3C). Similar results were found when DA neurons were analyzed for Th-T staining 48 and 72 hours after treatment with α-Syn seeds, with increased protein aggregation over time in all DA neurons compared to the 24hr treatment as well as average signal intensity higher in GBA^N370S^, LRRK2^G2019S^ and SNCA^A53T^ DA neurons compared to the AHN neurons at all time points (Figure 3C).

**Figure 3:**
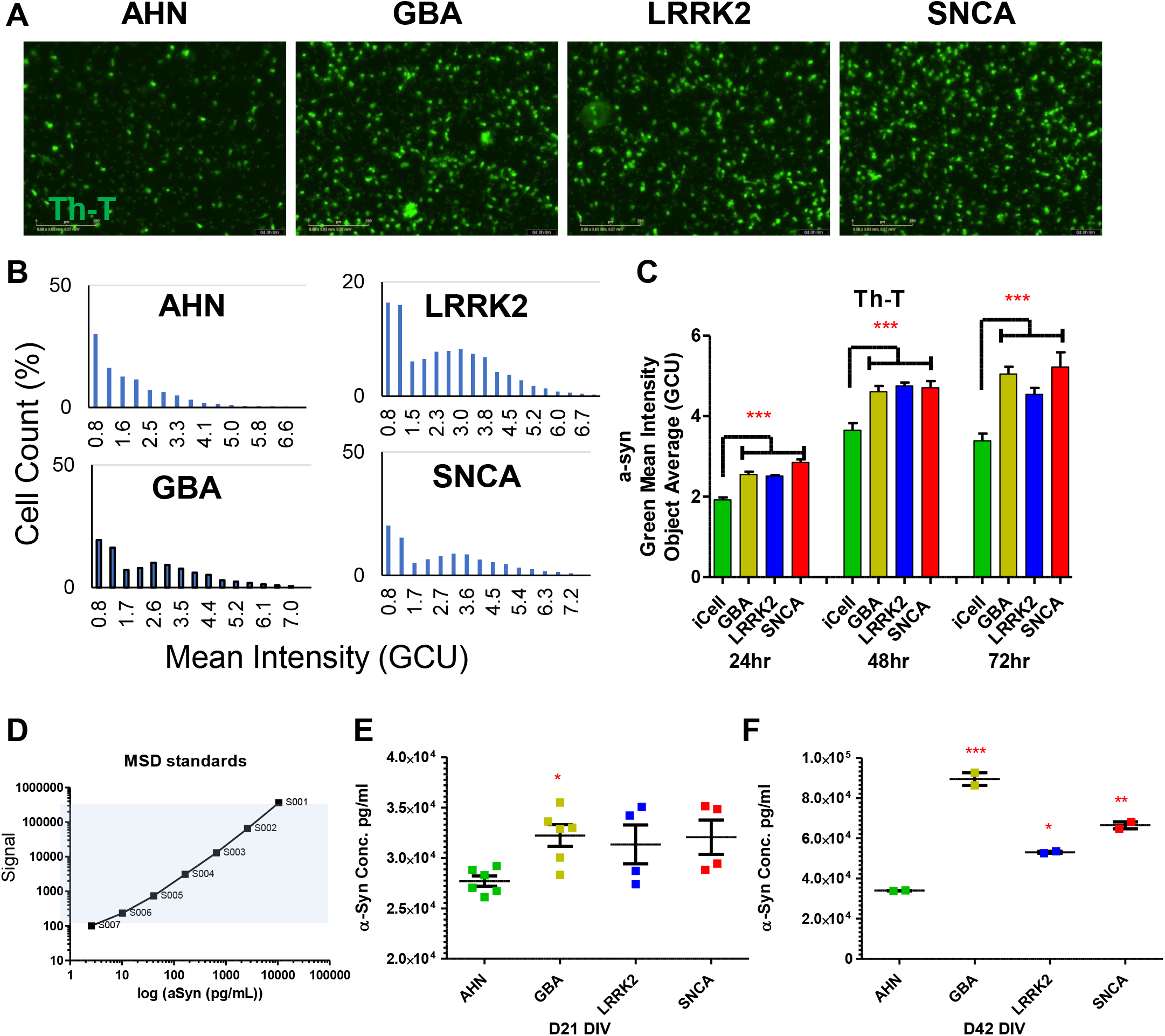
α-Syn protein accumulation in the DA neurons. (A) Thioflavin (Th-T) staining of α-Syn at 22 days post thaw in mutant and AHN DA neurons 24 hours after adding recombinant mouse α-Syn protein aggregate (4 μg/mL). (B) All α-Syn-positive cells were counted and Th-T signal intensity of individual neurons was plotted into bins of signal expression. (C) Kinetics of Th-T signal in DA neurons at 24, 48, and 72 hours after addition of active recombinant mouse α-Syn protein aggregate (n = 4, One-way ANOVA with Dunnett’s multiple comparisons test). (D) Standard curve of serially diluted α-Syn protein using the U-PLEX Human α-Syn Kit. Accumulation of α-Syn in mutant and AHN DA neurons at 21 (E) and 42 (F) days post thaw using the U-PLEX Human α-Synuclein Kit (n = 4, *P-value<0.05, **P-value<0.01, ***P-value<0.001 compared to AHN One-way ANOVA with Dunnett’s multiple comparisons test).

Th-T can bind to other amyloidogenic proteins in aggregates and may not be specific to the α-Syn aggregation. For confirmation of Th-T results, Meso Scale Discovery (MSD) antibodies were used for detection of α-Syn expression in the mutant and AHN neurons (Figure 3 D-F). The same amount of protein (0.2 μg) was used in each group and three replicates were run at each timepoint. Results from DA neurons cultured for 21 days post thaw showed more α-Syn expression in the *GBA*^N370S^ mutant cells compared to AHN neurons. *LRRK2*^G2019S^ and *SNCA*^A53T^ DA neurons also showed increases in α-Syn expression at day 21 but differences were not significant due to the variability among experimental replicates. In the later time point (42 days post thaw), significantly enhanced α-Syn expression was observed across all mutant DA neurons compared to the AHN neurons, with *GBA*^N370S^ being the highest and *LRRK2*^G2019S^ the lowest among the mutant DA neurons. These data show that mutant neurons have increased α-Syn expression and protein aggregation.

### Mutant DA neurons had higher mitochondrial stress

Accumulation of α-Syn has been associated with decreased mitochondrial membrane potential, and increased levels of mitochondrial oxidative stress in multiple cellular models^23^. Since *GBA*^N370S^ and *LRRK2*^G2019S^ display increased a-syn expression and aggregation, we hypothesized these neurons would be more prone to oxidative stress. To test our hypothesis, we cultured DA neurons for 21 days post thaw and evaluated mitochondrial oxidative stress using MitoSOX dye. Rotenone was used to artificially induce reactive oxygen species (ROS) in neurons as positive control group.

The average MitoSOX fluorescence per neuron was quantified (Figure 4A) and compared across AHN, GBA^N370S^, LRRK2^G2019S^ and SNCA^A53T^ DA neurons. All PD mutant DA neurons showed higher signal intensity per neuron, with significant increases in *GBA*^N370S^ and *SNCA*^A53T^ compared to the AHN neurons (Figure 4B). As expected, rotenone treatment increased the average MitoSOX intensity across all DA neurons compared to their untreated controls. More importantly, LRRK2^G2019S^ DA neurons showed significant increase compared to AHN under treatment, along with GBA^N370S^ and SNCA^A53T^ DA neurons. We also quantified the number of MitoSOX-positive cells in all groups and confirmed increased MitoSOX positive cells in the *GBA*^N370S^ and *SNCA*^A53T^ DA neurons compared to the AHN neurons. Rotenone treatment did not increase the number of MitoSOX positive cells in the mutant DA neurons but did increase the number of MitoSOX positive cells in AHN neurons (Figure 4C).

**Figure 4:**
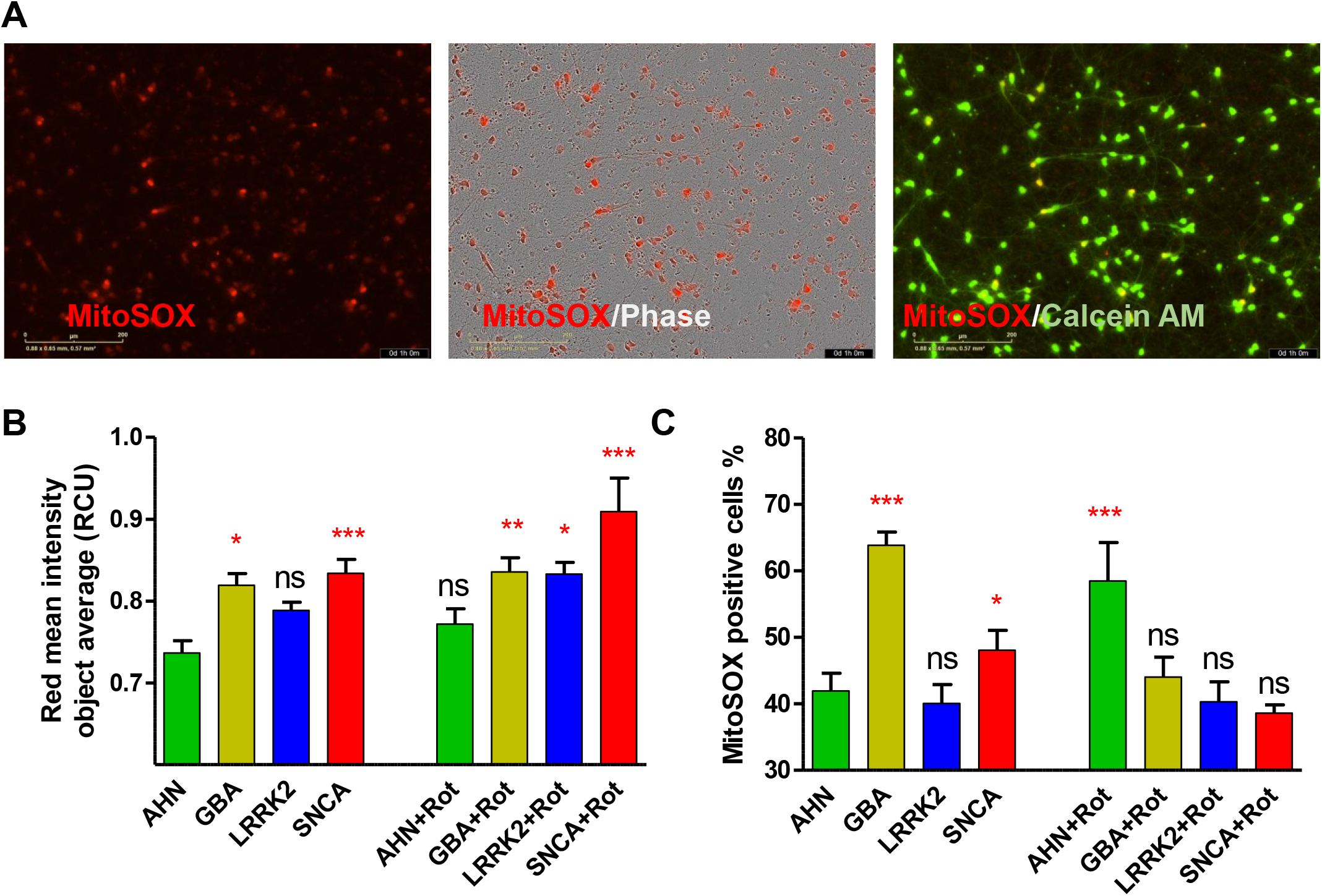
Increased mitochondrial oxidative stress in LRRK2^G2019S^, GBA^N370S^, and SNCA^A53T^ DA neurons. (A) Representative images of mutant and AHN DA neurons stained with MitoSOX dye (left), MitoSOX overlayed with phase image (middle), and MitoSOX co-stained with Calcein AM for live cells (right). (B, C) Quantitative data for the MitoSOX signal intensity (B, means ± SD) and percentage of the MitoSOX-positive cells in the mutant neurons compared to the AHN neurons (E). Rotenone (30-minute incubation) was added as a positive control to induce ROS in the mitochondria. (n=8-16, *P-value<0.05, **P-value<0.01, ***P-value<0.001 one way ANOVA with Dunnett’s test for mean comparisons against the AHN group).

### Mutant DA neurons demonstrated altered network activity and Ca^2+^ oscillations

DA neurons form neuronal networks when cultured *in vitro,* developing synchronized bursting activity that matures over time. To elucidate potential differences in neural network formation and maturation in *GBA*^N370S^, *LRRK2* ^G2019S^, *SNCA*^A53T^ and AHN neurons, we plated DA neurons on multielectrode array plates and monitored electrophysiological activity over time. Overall, DA neurons showed spontaneous activity almost immediately after plating (data not shown) and their activity increased with time in culture, peaking at three weeks (21 days post thaw) for all mutant DA neurons. In contrast, the peak spontaneous activity of AHN cells was delayed to 26 days post thaw and maintained out to day 35 (Figure 5A), the longest timepoint cultured. Altered network activity was observed across multiple parameters in mutant DA neurons. Across all DA neuron groups most of the activity occurred as network bursts (60-80%). However, by day 35 in culture the network burst percentage was remarkably decreased in mutant DA neurons (Figure 5B). General activity of neurons, described as mean firing rate (Hz), decreased over time in mutant DA neurons but increased for AHN neurons (Figure 5C). The most different activity parameter across DA neuron groups was the network burst frequency, which was higher in the mutant neurons at all time points compared to AHN neurons. *LRRK2*^G2019S^ neurons showed the highest burst frequency while *GBA*^N370S^ neuron burst frequency was only slightly higher than AHN neurons (Figure 5D). The synchrony index, a parameter indicating network maturity, increased over time in AHN DA neurons. In contrast, mutant DA neurons displayed more erratic synchrony over time with overall lower synchrony values at each time point compared to AHN cultures (Figure 5E).

**Figure 5:**
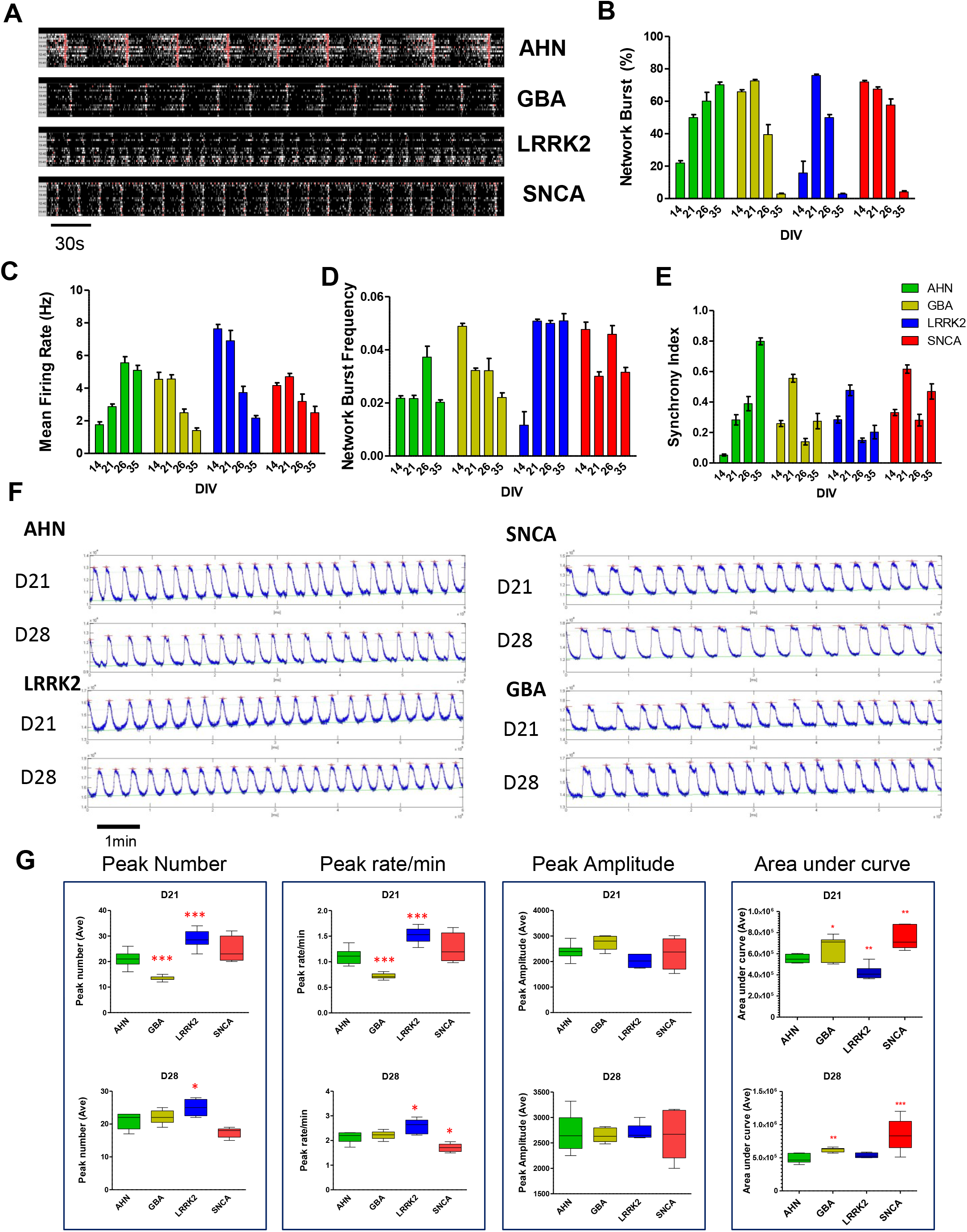
Neuronal network activity in LRRK2^G2019S^, GBA^N370S^, and SNCA^A53T^ DA neurons. (A) Representative MEA raster plots of DA neurons at 35 days post thaw showing neuronal activity and network. MEA activity a 14-, 21-, 26-, and 35-days post thaw was quantified and displayed as network burst percentage (B), mean firing rate (C), network burst frequency (D) and synchrony index (E). (F) Representative calcium traces recorded at 21- and 28-days post thaw for LRRK2^G2019S^, GBA^N370S^, and SNCA^A53T^, and AHN DA neurons cultured as spheroids. (G) Quantification of calcium traces for peak number, peak rate, peak amplitude and area under the curve summarized for both day 21 and 28 post plating (n≥6, *P-value<0.05, **P-value<0.01, ***P-value<0.001 one way ANOVA with Dunnett’s test for mean comparisons against the AHN group).

To further evaluate neural network dynamics between *GBA*^N370S^, *LRRK2*^G2019S^, *SNCA*^A53T^ and AHN neurons, we monitored calcium oscillations in DA neurons cultured as spheroids. Calcium oscillations were captured at day 21 and 28 in culture using a 10-minute baseline reading on the FDSS/μCELL instrument. Calcium traces showed altered oscillations in the mutant neurons compared to the AHN cultures (Figure 5F). Calcium oscillations across neurons showed diverse phenotypes with lower activity in *GBA*^N370S^ neurons, marked by lower peak numbers and peak rate, and higher activity in *LRRK2*^G2019S^ DA neurons (Figure 5G), suggesting hypo-excitability in *GBA*^N370S^ and hyperexcitability in the *LRRK2*^G2019S^ mutants. This observation is similar in both MEA and calcium oscillation results. *SNCA*^A53T^ neurons showed larger intracellular calcium transients, which is depicted as increased area under the peaks, indicating lower cytosolic buffering for calcium and thereby higher cytosolic calcium.

### TH and DDC dopamine synthesis enzymes highly expressed in the GBA^N370S^ and *LRRK2*^G2019S^ DA neurons leading to higher dopamine release

RNAseq analysis revealed that tyrosine hydroxylase (TH) and dopamine decarboxylase (DDC) gene transcripts were differentially regulated in the GBA^N370S^ and *LRRK2*^G2019S^ DA neurons (Figure 6A, Table S2). To further investigate changes in dopamine synthesis and release pathway, we detected protein expression of TH, DDC and other synthesis-related proteins. Western blot confirmed that TH protein expression was significantly increased in the *GBA*^N370S^ and *LRRK2*^G2019S^ mutant neurons, while its expression in the *SNCA*^A53T^ mutant cells showed a slight but not significant decrease (Figure 6B, 6C). Since these genes code essential enzymes in the dopamine synthesis pathway, we decided to evaluate the dopamine release in cultured DA neurons. At 21 days post thaw, dopamine release under basal resting conditions (isotonic HBSS buffer) showed that *GBA*^N370S^ and *LRRK2*^G2019S^ released slightly more dopamine (2-4 ng/mL). This increase was not significant compared to AHN neurons. However, when stimulated with KCl, *GBA*^N370S^ and *LRRK2*^G2019S^ neurons showed a significant increase in dopamine release compared to AHN neurons. Interestingly, *SNCA*^A53T^ dopamine release remained at the same level as AHN neurons regardless of the stimulation (Figure 6D, 6E). These results indicated that in the *GBA*^N370S^ and *LRRK2*^G2019S^ mutant DA neurons, dopamine synthesis and release are disrupted, producing and releasing more dopamine than normal cells.

**Figure 6:**
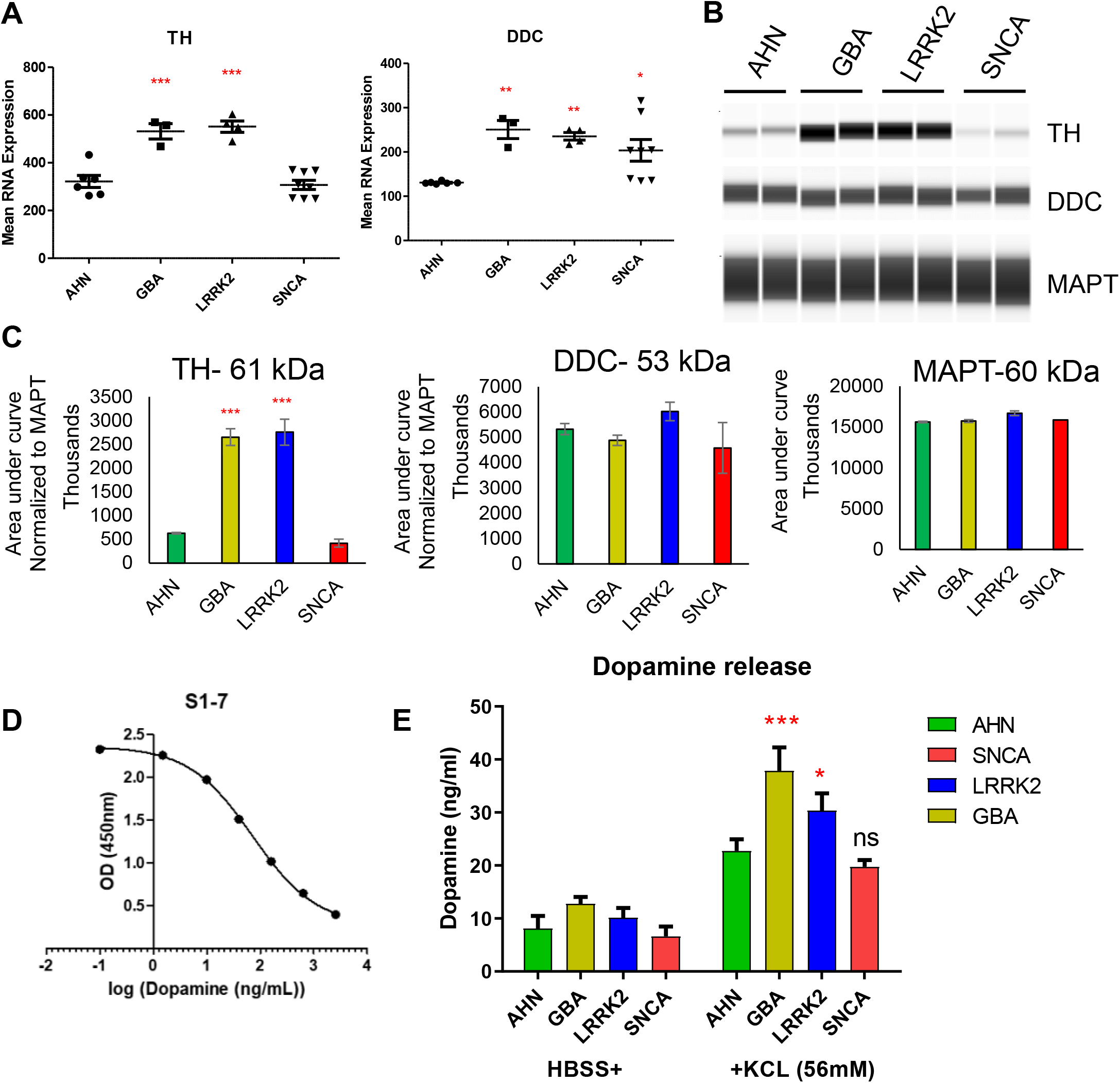
Increased TH expression and dopamine release in *GBA*^N370S^ and *LRRK2*^G2019S^ mutant DA neurons. (A) mRNA expression (FPKM) evaluated by RNAseq showed significantly increased TH and DDC in *GBA*^N370S^ and *LRRK2*^G2019S^ DA neurons at day 14 post thaw. (B) Western blot of TH and DDC protein expression detected at day 14 using the WES (ProteinSimple). Two replicates were run for each group. (C) Quantification of Western blot protein expression based on the area under the curve for each protein and normalized to the MAPT expression. (D) Standard curve of dopamine release assay was generated using standard samples of dopamine at defined concentrations. (E) At 21 days in culture mutant and AHN DA neurons were stimulated with HBSS or KCl and dopamine release was quantified using competitive ELISA (n=3, *P-value<0.05, **P-value<0.01, ***P-value<0.001 one way ANOVA with Dunnett’s test for mean comparisons against the AHN group).

### Mutant DA neurons showed decreased GCase activity

RNAseq data showed a reduction of *GBA* transcripts in both *GBA*^N370S^ and *LRRK2*^G2019S^ mutant neurons, but no significant change in the *SNCA*^A53T^ (data not shown), compared to AHN neurons (Figure 7A). To further investigate if the changes in the mRNA level are translated to the protein level, day 14 post thaw mutant and AHN DA neurons were collected and assayed for GBA protein expression. Interestingly, western blot analysis did not reveal significant differences in GBA protein expression across mutant and AHN groups (Figure 7B, 7C). Less GBA activity in the *GBA*^N370S^ and *LRRK2*^G2019S^ neurons were reported previously ^24,25^, therefore we next measured GCase enzyme activity in DA neurons. Fluorophore 4-methylumbelliferone (4-MU) was used as the substrate for GCase and optimized for concentration within the assay. We found that 10mM 4-MU gave maximum level of activity of the enzyme with human recombinant GCase (Figure 7D). 4-MU is also a substrate for other galactosidase enzymes, so taurocholate, an inhibitor of beta-galactosidase enzymes, was added to all conditions in the assay. For normalization purpose and to determine enzyme specificity, different concentrations of CBE, a specific blocker of the GCase enzyme, was also tested and 2mM CBE was chosen as the optimal concentration (Figure 7E). Figure 7F shows that enzyme activity has a linear correlation with the amount of protein in the assay, therefore 5μg of protein was selected for the final assay. Our results confirmed that at day 14 post thaw, GBA activity was significantly decreased in *GBA*^N370S^, *LRRK2* ^G2019S^, *SNCA*^A53T^ DA neurons compared to the AHN. *GBA*^N370S^ DA neurons showed the lowest activity while *SNCA*^A53T^ had the highest activity among the three mutations (Figure 7G).

**Figure 7:**
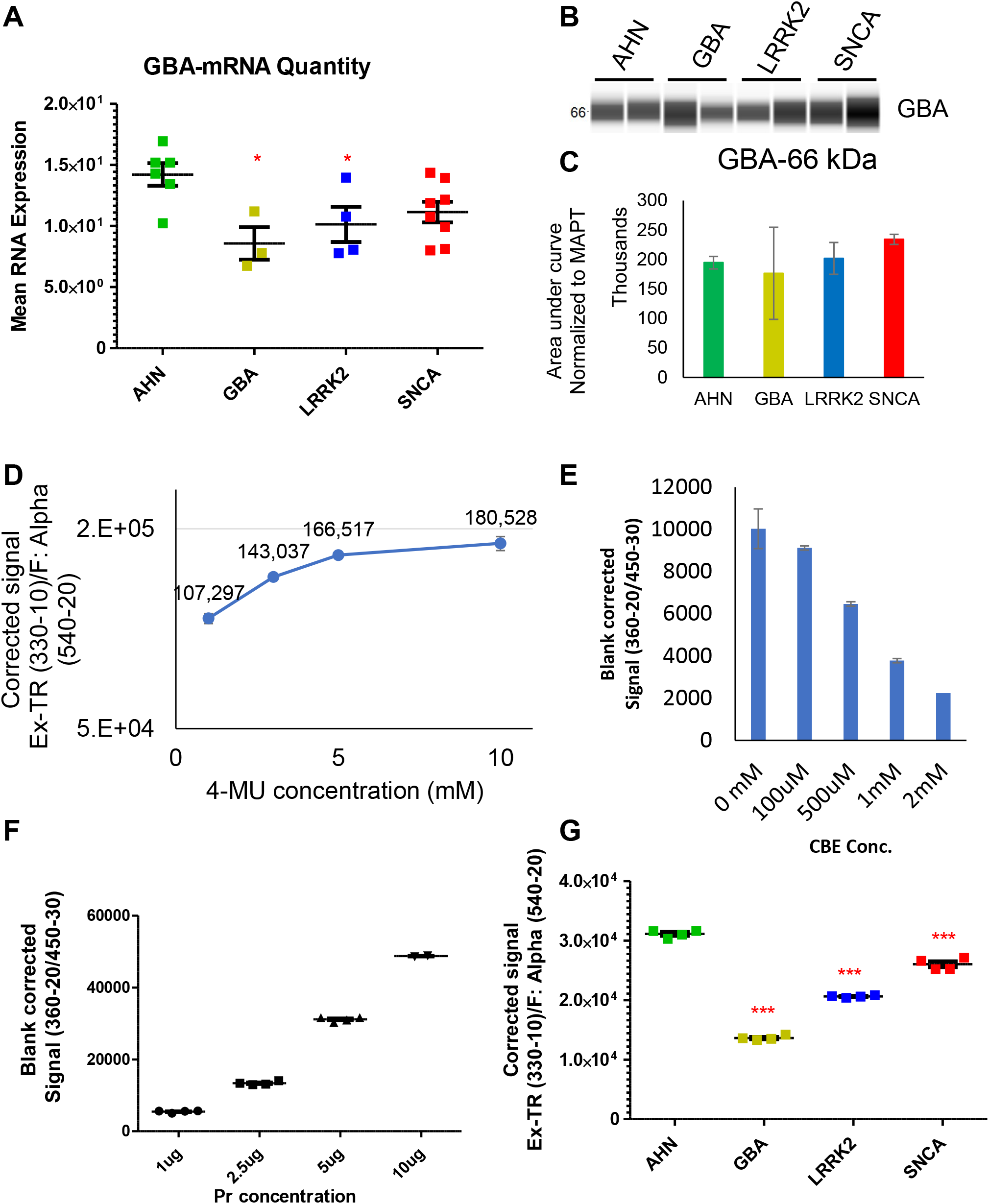
GCase enzyme activity in DA neurons. (A) GBA mRNA expression (FPKM) at day 14 post thaw for DA neurons showed significant activity reduction in GBA^N370S^ and LRRK2^G2019S^ mutant neurons compared to AHN neurons. (B) Western blot of GBA protein expression detected at 14 days post thaw using the WES (ProteinSimple). Two replicates were run for each group. (C) Western blot GBA protein expression quantified and normalized to MAPT expression. Optimum concentrations of 4-Methylumbelliferyl-β-D-glucopyranoside (4-MU) (D) and the GCase specific inhibitor, Conduritol B Epoxide (CBE) (E), were determined to be 10 mM and 2 mM, respectively. (F) GCase activity measured for different concentrations of protein lysate from AHN DA neurons. (G) Protein lysate (5 μg) from AHN and mutant DA neurons treated with or without CBE, were measured for GCase activity using 4-MU. Lower GCase activity was observed in disease DA neurons compared to AHN (n≥4, *P-value<0.05, **P-value<0.01, ***P-value<0.001 one way ANOVA with Dunnett’s test for mean comparisons against the AHN group).

### Chronic ABX treatment rescued the GCase activity and calcium oscillation in the *GBA*^N370S^ DA neurons

In a series of experiments, we tested 12 compounds previously shown to enhance the GCase activity in neurons (Table S3), for their ability to correct the aberrant GCase activity observed in the mutant DA neurons. First, we treated *GBA*^N370S^ and *LRRK2*^G2019S^ DA neurons with 12 compounds and compared GCase activity levels to AHN neurons (Figure S1A). This screening showed significant improvement of Gase activity using ABX (10 μM), cyclodextrin (100 μM), NN-DNJ (1 μM), and echinacoside (1 μM). These compounds were selected for further combinatorial screens (Figure S1B). In the second round of screenings, combinations of these four compounds were tested along with acute (3 day) and chronic (14 day) treatments of single compounds. None of the compound combinations showed superior performance compared to the single compound treatment (Figure S1B). However, chronic treatment with ABX (10μM) or NN-DNJ (1μM) significantly increased GCase activity in all three mutant DA neurons compared to the DMSO-treated control cells (133658±6086 (A-Ch), 108211±3269 (N-Ch), 90302±2301 (DMSO)) (Figure S1C). AHN neurons did not show significant change when treated with ABX. ABX treatment of *LRRK2*^G2019S^ DA neurons resulted in increases in GCase activity that was comparable to basal GCase activity in AHN DA neurons. ABX treatment was not sufficient for GCase activity recovery in *GBA*^N370S^ mutant neuron, showing significantly lower activity than AHN neurons (Figure S1C).

We next investigated if chronic exposure to ABX could rescue the calcium oscillation perturbations observed in *GBA*^N370S^, *LRRK2* ^G2019S^, and *SNCA*^A53T^ DA neurons. DA neurons were cultured as spheroids and exposed chronically (14 days) to ABX (10 μM) and assayed on 21 days post plating. Calcium oscillation showed an increase in the rate and number of calcium peaks in ABX treated neurons and decreased the area under the curve for *GBA*^N370S^ and *SNCA*^A53T^ neurospheres (Figure 8A-D). In *GBA*^N370S^ mutant cells chronic ABX rescued the peak number, rate and area under the curve to the levels observed in AHN neurons, but resulted in a hyperexcitable phenotype in *LRRK2*^G2019S^, *SNCA*^A53T^, and AHN neurons (Figure 8A-D).

**Figure 8:**
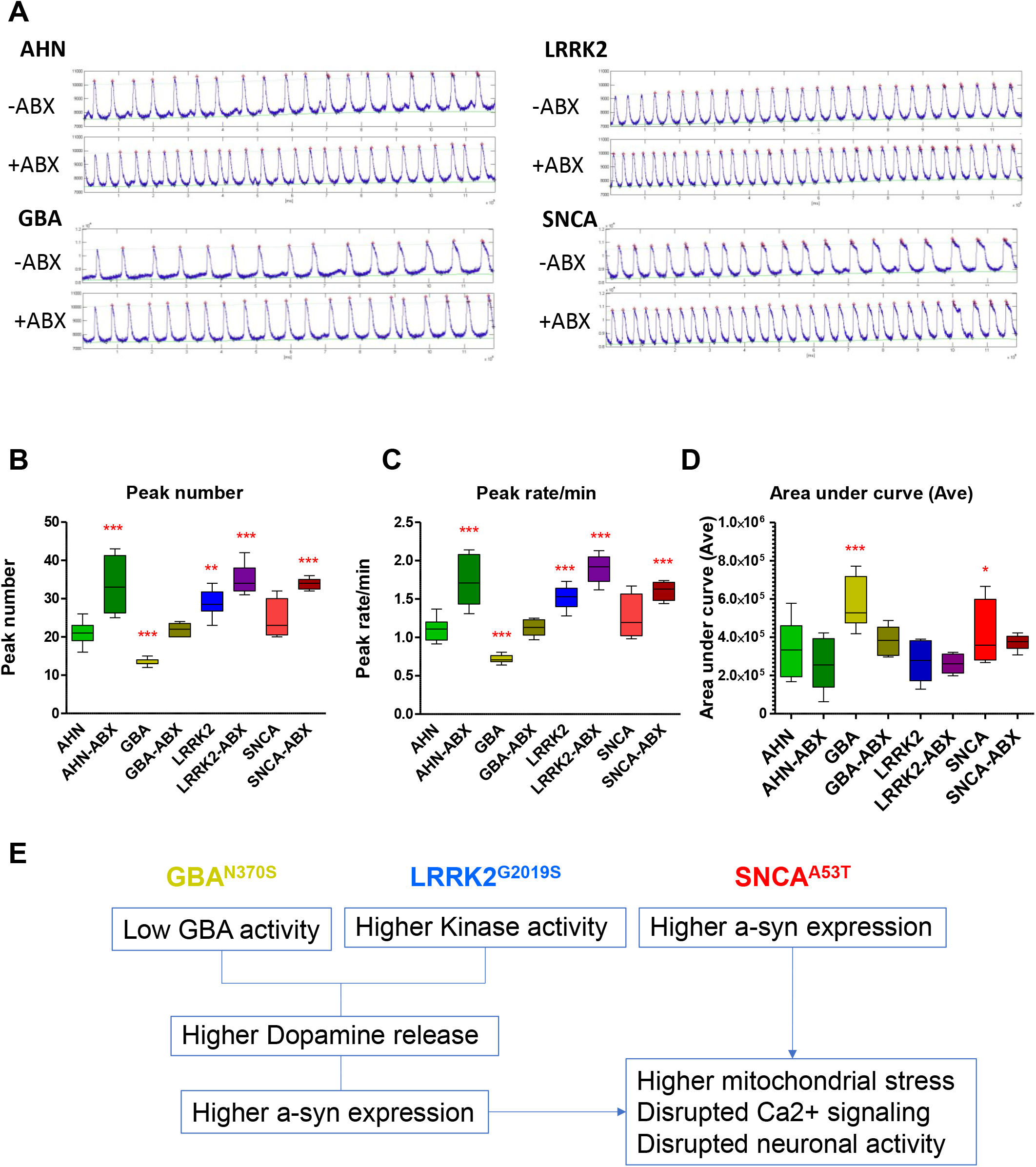
Calcium imaging of iPSC-derived DA neuron 3D spheroids. (A) Representative calcium oscillation traces recorded from 21 days post thaw DA neurons with or without 10 μM ABX for 14 days. Quantification of calcium traces for peak number (B), peak rate (C), and area under the curve (D). Compared to the untreated AHN DA neurons, treatment with ABX resulted in hyperexcitability of AHN, *LRRK2^G2019S^* and *SNCA*^A53T^ DA neurons but rescued the peak number and rate in GBA^N370S^ mutant DA neurons (n≥6, *P-value<0.05, **P-value<0.01, ***P-value<0.001 one way ANOVA with Dunnett’s test for mean comparisons against the AHN group). (E) Schematic of diverging phenotypes between GBA^N370S^, LRRK2^G2019S^ and SNCA^A53T^ DA neurons. GBA^N370S^ and LRRK2^G2019S^ DA neurons uniquely displayed abnormal dopamine synthesis and release which was not detected in SNCA^A53T^ DA neurons. Phenotypes across the three mutant DA neurons converged on high accumulation and production of α-Syn. Early developmental aberrations in the dopamine pathway and subsequent increases in mitochondrial stress may contribute to PD-related α-Syn accumulation, aberrant neural activity, and neurodegeneration.

## Discussion

In this study we described the generation of DA neurons from clinically diagnosed Parkinson’s disease patient-derived iPSCs containing either the *LRRK2*^G2019S^ or *GBA*^N370S^ mutation and compared their phenotypes to SNCA^A53T^ engineered and AHN DA neurons. Interestingly, LRRK2^G2019S^ and GBA^N370S^ DA neurons demonstrated increased dopamine release and enhanced expression of genes involved in dopamine synthesis and release which were not observed in SNCA^A53T^ mutant DA neurons ^26,27^. The common phenotypes across the mutant DA neurons include α-Syn accumulation, mis-regulation of network activity and calcium oscillations. In addition, patient-derived and engineered PD DA neurons showed decreased activity of GCase enzyme which is consistent with previous studies suggesting DA neurons have reduced capability for clearing misfolded and aggregated proteins ^9,28–31^. Together these data suggest a divergent etiology of PD pathogenesis between *GBA*^N370S^, *LRRK2*^G2019S^ and SNCA^A53T^ DA neurons, described in Figure 8E and discussed in further detail throughout the discussion.

Multi-factorial neurodegenerative diseases, including PD, have dysregulation of multiple common cellular pathways that leads to cellular malfunction and neuronal cell death. Neuroprotective agents have historically been the focus for therapeutic intervention, with the goal of slowing disease progression. However, at the time of disease symptom manifestation significant cell death has already occurred and remaining neurons display improper function, limiting the efficacy of neuroprotective drugs on disease symptoms. Due to the lack of the biomarkers for early detection of PD and the paucity of patient brain tissue across different ages and stages of the disease, it is difficult to identify and treat the root cause of the PD. Studies using stem cell derived DA neurons from PD patients, which retain their genetic predisposition for the disease including epigenetic and environmental factors, provides an accessible model for discovery of early cellular mechanisms that underly PD pathogenesis and DA neuron vulnerability to degeneration. For this study, iPSCs were differentiated into DA neurons using a batch-to-batch, consistent protocol resulting in similar midbrain DA neuron marker expression and purity across AHN, *GBA*^N370S^, *LRRK2* ^G2019S^, and *SNCA*^A53T^ lines. This consistency in DA neuron production was critical for identifying unique phenotypes that could provide a better mechanistic understanding between PD pathology and genetic risk factors.

Indicators of oxidative stress in DA neurons have been observed in PD patient brain samples ^11,32,33^, animal models ^34,35^ and neurons differentiated from iPSCs ^36–38^. In addition, mitochondrial stress is associated with α-Syn aggregation ^39,40^ and defects in protein homeostasis have been attributed to altered neuronal function, including impaired calcium signaling ^41,42^ and hyperexcitability ^43,44^. Despite progress in elucidating these disease phenotypes, the root cause of the disease remains elusive with no origin or mechanism for oxidative damage and mitochondrial stress in DA neurons. Using precise DA neuron differentiation of patient-derived iPSCs, these data show that *GBA*^N370S^ and *LRRK2*^G2019S^ patient derived neurons produce and release more dopamine at very early stages of development, suggesting that dopamine neurotoxicity may be an intrinsic underlying mechanism for gradual cell death in DA neurons.

In this study, we observed increased dopamine synthesis and release in LRRK2^G2019S^ and GBA^N370S^ DA neurons, with no change in dopamine release observed in SNCA^A53T^ DA neurons (Figure 6). Patient-derived iPSC-derived SNCA^A53T^ DA neurons have previously been shown to have normal dopamine release, suggesting that the phenotypes we report in LRRK2 ^G2019S^ and GBA^N370S^ DA neurons is novel and not reflective of different sources of mutation (patient vs. engineered mutation) ^26,27^.Dopamine, once released from DA neurons, is uptaken by dopamine receptors in post-synaptic neurons or surrounding glial cells. Modulation of dopamine synthesis and release is hypothesized as a contributing mechanism associated with PD neurodegeneration, but the underlying mechanism remains unclear. Dopamine degradation pathways have been reported to produce free radicals and quinones in neurons and glia, resulting in build-up of free cysteinyl-catechols and the deaminated metabolite, dihydroxy phenylacetic acid (DOPAC) ^45,46^. Injection of reserpine, which releases stored dopamine in the synaptic vesicles, was also found to increase ^32^. In addition, evidence indicating patient DA neurons are defective in vesicular monoamine transporter 2 (VMAT2)-mediated DA transport ^47,48^. Free cysteinyl-catechols have been shown to covalently bind to thiol groups of proteins and make them resistant to degradation pathways and perturb protein homeostasis. Conversely, excess dopamine converted by monoamine oxidase to DOPAC could generate oxidative stress and harm organelles, including mitochondria, which are crucial for cell metabolism and health ^12,49^. Although, generation of quinones, DOPAC, and cysteinylated intermediates have not been observed in iPSC-derived DA models, subsequent studies investigating these molecules within PD models may connect our observation of increased dopamine synthesis and release in patient DA neurons with the disruption of the UPR (unfolded protein response) and protein quality control in the neurons.

iPSC-derived DA neurons harboring LRRK2^G2019S^, GBA^N370S^, or SNCA^A53T^ mutations were found to have increased mitochondrial stress and higher α-Syn expression in this study. These data are consistent with previous studies in human and animal models. Interestingly, *GBA*^N370S^ mutant neurons showed the highest α-Syn accumulation of the three mutant neurons, while also having the lowest GCase activity. These data suggest a potential association between GCase activity and α-Syn protein turnover and aggregation, as depicted in the schematic diagram in Figure 8E.

Baseline calcium oscillation data on day 21 revealed altered patterns in all mutant DA neurons, with *SNCA*^A53T^ and *GBA*^N370S^ mutant neurons showing slow and broad spike patterns, which is an indication of higher cytosolic calcium and less buffering capacity. Conversely, *LRRK2*^G2019S^ mutant neurons had faster calcium oscillations which aligned with the hyperexcitable phenotype (increased network burst frequency) observed on MEA activity. One possible mechanism underlying this observation is that high expression of α-Syn has been shown to disrupt calcium homeostasis and potentially alter neuronal activity ^50^. This is supported by our observation of increased α-Syn accumulation in all mutant PD disease lines. Another possible contributor to altered calcium dynamics may be through expression of calcium-related genes or genes related to synaptic function. Indeed, our DEG RNAseq analysis revealed *LRRK2*^G2019S^, *GBA*^N370S^, and *SNCA*^A53T^ DA neurons all showed altered gene expression within calcium-related pathways. This is corroborated by a study in PARK2 mutant iPSC derived DA neurons, where increased expression of the T-type calcium channel contributed to the dysregulation of calcium homeostasis and resting calcium levels ^51^. Lower cytosolic calcium buffering, autonomous pacemaking, and Cav1.3 channel (L-type calcium channel) opening that triggers calcium-induced calcium release (CICR) could result in the generation of large calcium transients ^52^.

Our data showing dysregulation of dopamine release and genes related to dopamine synthesis suggests a possible association with α-Syn accumulation and calcium homeostasis. Overexpression of α-Syn has been shown to decrease the auto-inhibitory function of D2 receptors in DA neurons and increase intracellular and extracellular dopamine levels and tyrosine hydroxylase expression ^53^. Furthermore, calcium has been shown to directly and indirectly increase α-Syn pathology ^54–56^. Higher calcium transients promote α-Syn aggregation by enhancing calmodulin and membrane binding ^57^. High cytosolic calcium is known to activate calpains and increase cleavage of intracellular proteins ^58^, including α-Syn at its c-terminus, thus promoting its aggregation ^59^.

Notably, this study included a screen of compounds to rescue the deficiency of GCase activity observed in LRRK2^G2019S^ and GBA^N370S^ DA neurons (Figure S1). We found that chronic treatment with ABX, a molecule previously reported to stabilize GCase enzyme activity in Gaucher disease (GD) fibroblasts ^60^ and used in a clinical trial for PD treatment ^61^, could enhance the GCase activity in all mutant neurons. This effect was especially pronounced with *GBA*^N370S^ DA neurons which exhibited the largest reduction in GCase mRNA and activity across the mutant DA neurons.

Dysregulation of calcium homeostasis is reported to have a pivotal role in the pathogenesis of PD ^62^, therefore we tested the effects of ABX on calcium dynamics in all mutant neurons and AHN. Chronic ABX treatment resulted in an unfavorable phenotype, increased calcium activity and hyperactivity, in *LRRK2*^G2019S^, *SNCA*^A53T^ and AHN DA neurons. However, ABX returned *GBA*^N370S^ DA neuron calcium dynamics to a phenotype similar to untreated AHN DA neurons (Figure 8B-D). Our results indicate that phenotypic screening of iPSC-derived DA neurons across multiple assays will be essential for successful drug discovery platforms. This strategy can identify compounds that either have similar or divergent effects across different donors, as exemplified in this study where one compound (ABX) has potential benefit for one genetic background but may not be applicable to patients with different genetic backgrounds.

Our study demonstrates the capability of human iPSC-based disease modeling (“disease-in-a-dish”) to study pathogenic mechanisms of early onset PD. Based on the data presented in this study, we suggest a novel mechanism contributing to the progression of the disease phenotype in the *GBA*^N370S^ and *LRRK2*^G2019S^ DA neurons (Figure 8E). Specifically, alterations in DA synthesis and release begin a cascade of pathway changes that increase the susceptibility of LRRK2^G2019S^ and GBA^N370S^ DA neurons to undergo degeneration. This is different than previously known for *SNCA*^A53T^, which is thought to be triggered by aberrant α-Syn aggregation. Therefore, in the absence of extrinsic or extracellular factors, intrinsic cell-autonomous factors seem to be sufficient to trigger neurodegeneration of DA neurons, at least in familial cases captured in the time frame of *in vitro* cultured neurons. Our confidence in these data and our experimental strategy is rooted in the consistent differentiation of the PD relevant cell type across donors, the ability to maintain DA neurons long-term in culture, and the use of multiple patient cell lines per condition, which enabled us to control for the inherent variability of human pluripotent stem cell lines. Adding multiple patient iPSCs from both familial and sporadic cases would further help to generalize the disease phenotypes that we show here and subsequent studies that compare each patient cell line to their isogenic control cell line could help to minimize effects of genetic background differences. In summary, we have identified unique *in vitro* phenotypes associated with PD that could be further helpful in understanding the PD root cause and could serve as a valuable tool for searching new therapeutics.

## Acknowledgements

We acknowledge Parkinson’s Progression Markers Initiative (PPMI) and the Michael J. Fox Foundation for Parkinson’s research for their partnership with Fujifilm Cellular Dynamics Inc. This study is fully funded by Fujifilm Cellular Dynamics Inc.

## Author Contributions

A.F.: Design and conception of the study, differentiation of iPSCs, performing, collecting and summarizing data from ICC, FC, DA release assay, MitoSOX staining, WB, MEA, a-syn expression, GCase activity assay and Ca2+ imaging assay, small molecule screen, writing of manuscript. K.B. and L.Z.: RNAseq data analysis, interpreting the results, writing of manuscript. R.F.: Neurosphere Ca2+ imaging assay. R.B.: data interpretation, editing the manuscript. S.D.: iPSCs establishment and characterization, engineering SNCA A53T line, editing the manuscript. C.C.: design and interpretation of Ca2+ imaging data and small molecule screen, writing of manuscript. S.Sch.: data interpretation, writing and editing of manuscript. J.L.: design and conception of the study, data interpretation, writing and editing of manuscript.

## Declaration of Interest

All authors are Fujifilm employees and declare no competing financial interests. Correspondence and requests for materials should be addressed to A.F. or J.L.

## Lead Contact

Further information and requests for sources should be directed to, and will be fulfilled by, the Lead Contacts, Dr. Ali Fathi or Dr. Jing (Jeanie) Liu.

## Materials Availability

Lines generated in this study are available from the Lead Contact with a completed Materials Transfer Agreement or purchasing directly from Fujifilm Cellular Dynamics, Inc. (www.fujifilmcdi.com)

## Data and Code Availability

The published article includes all datasets generated or analyzed during this study. The datasets supporting this study are available from the lead contact, Dr. Ali Fathi or Dr. Jing (Jeanie) Liu upon request.

---

**Figure S1:**
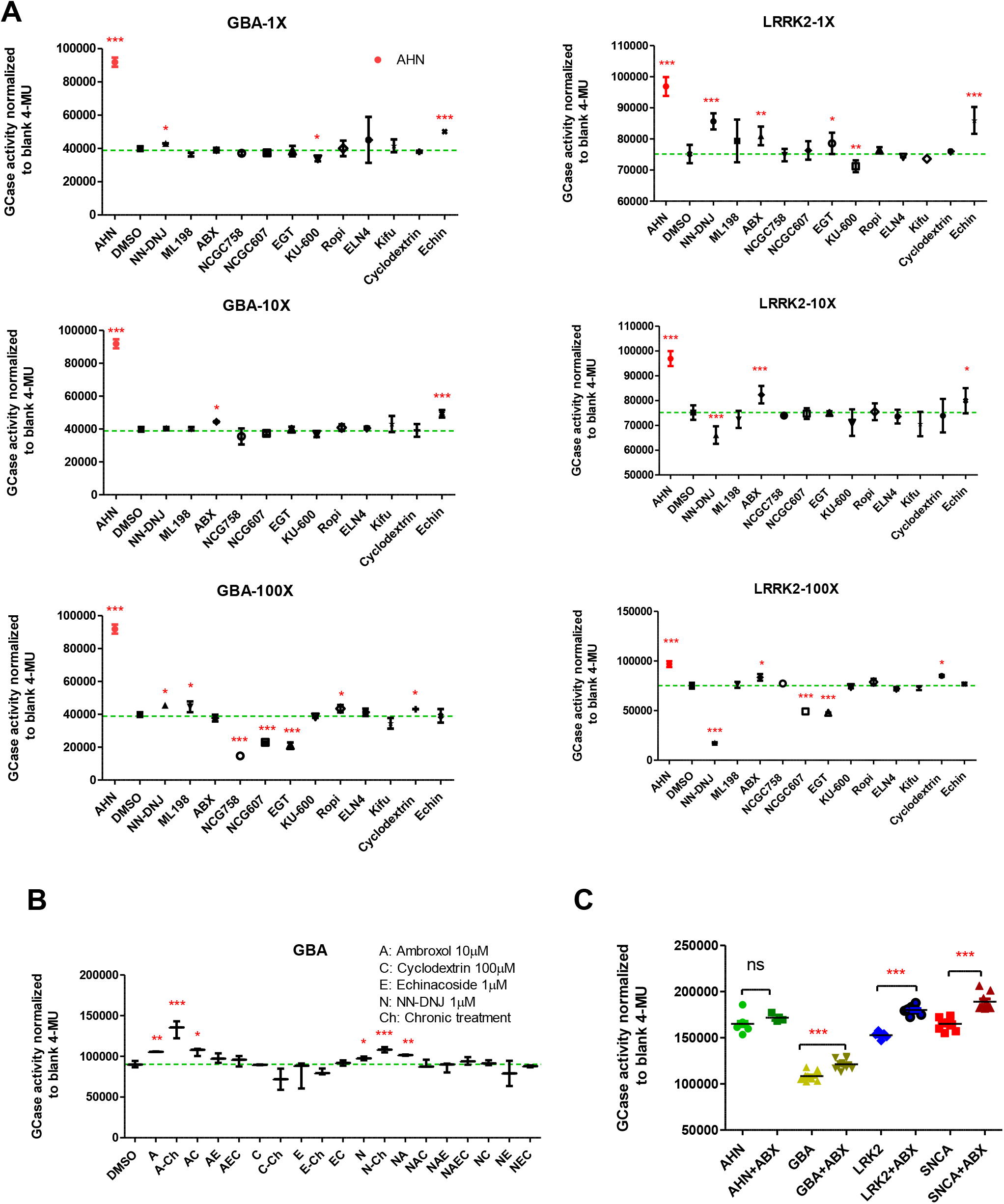
Compound screening for GCase activity in mutant DA neurons. GCase activity was measured in *GBA*^N370S^, *LRRK2*^G2019S^ and AHN DA neurons following treatment with various compounds (Table S3) reported to affect GCase activity. On day 18 post thaw DA neurons were dosed with various compounds at three concentrations (1X, 10X, and 100X). Three days after treatment (21 days post thaw) cells were lysed and total protein were determined. (A) Effects of 12 compounds at three concentrations on GCase activity in *GBA*^N370S^ and *LRRK2*^G2019S^ DA neurons with AHN as the control. (B) For combinatorial screen, 4 compounds displaying improved GCase activity were chosen from the initial screening. A matrix of compound combinations was tested in *GBA*^N370S^ cells. All combinations were tested for acute compound exposure (3 days) and single compound treatments were also tested for chronic exposure (two weeks). Combinatorial screen showed that chronic treatment with ABX (10uM) had the highest GCase activity in *GBA*^N370S^ cells. (C) Effects of ABX on 21 days post-thaw *LRRK2*^G2019S^, *SNCA*^A53T^, and *GBA*^N370S^ DA neurons was tested compared to AHN DA neurons. Treatment with ABX for two weeks significantly increased GCase activity in all mutant neurons. (Dotted lines indicate the GCase levels of the DMSO vehicle control. n=3-16, ***P-value<0.001 one way ANOVA with Dunnett’s test for mean comparisons against DMSO control.)

## References

1 Consortium, H. D. i. Induced pluripotent stem cells from patients with Huntington’s disease show CAG-repeat-expansion-associated phenotypes. Cell Stem Cell 11, 264–278 (2012). https://doi.org:10.1016/j.stem.2012.04.027

2 McGivern, J. V. et al. Spinal muscular atrophy astrocytes exhibit abnormal calcium regulation and reduced growth factor production. Glia 61, 1418–1428 (2013). https://doi.org:10.1002/glia.22522

3 Liu, H. et al. Spinal muscular atrophy patient-derived motor neurons exhibit hyperexcitability. Sci Rep 5, 12189 (2015). https://doi.org:10.1038/srep12189

4 Ye, L. et al. Induced pluripotent stem cells offer new approach to therapy in thalassemia and sickle cell anemia and option in prenatal diagnosis in genetic diseases. Proc Natl Acad Sci U S A 106, 9826–9830 (2009). https://doi.org:10.1073/pnas.0904689106

5 Ikonomou, L. et al. Stem Cells, Cell Therapies, and Bioengineering in Lung Biology and Disease 2021. Am J Physiol Lung Cell Mol Physiol (2022). https://doi.org:10.1152/ajplung.00113.2022

6 Garcia-Leon, J. A., Vitorica, J. & Gutierrez, A. Use of human pluripotent stem cell-derived cells for neurodegenerative disease modeling and drug screening platform. Future Med Chem 11, 1305–1322 (2019). https://doi.org:10.4155/fmc-2018-0520

7 Mishima, T., Fujioka, S., Fukae, J., Yuasa-Kawada, J. & Tsuboi, Y. Modeling Parkinson’s Disease and Atypical Parkinsonian Syndromes Using Induced Pluripotent Stem Cells. Int J Mol Sci 19 (2018). https://doi.org:10.3390/ijms19123870

8 Byers, B., Lee, H. L. & Reijo Pera, R. Modeling Parkinson’s disease using induced pluripotent stem cells. Curr Neurol Neurosci Rep 12, 237–242 (2012). https://doi.org:10.1007/s11910-012-0270-y

9 Clark, L. N. et al. Mutations in the Parkinson’s disease genes, Leucine Rich Repeat Kinase 2 (LRRK2) and Glucocerebrosidase (GBA), are not associated with essential tremor. Parkinsonism Relat Disord 16, 132–135 (2010). https://doi.org:10.1016/j.parkreldis.2009.05.008

10 Prots, I. et al. alpha-Synuclein oligomers induce early axonal dysfunction in human iPSC-based models of synucleinopathies. Proc Natl Acad Sci U S A 115, 7813–7818 (2018). https://doi.org:10.1073/pnas.1713129115

11 Sanders, L. H. & Timothy Greenamyre, J. Oxidative damage to macromolecules in human Parkinson disease and the rotenone model. Free Radic Biol Med 62, 111–120 (2013). https://doi.org:10.1016/j.freeradbiomed.2013.01.003

12 Burbulla, L. F. et al. Dopamine oxidation mediates mitochondrial and lysosomal dysfunction in Parkinson’s disease. Science 357, 1255–1261 (2017). https://doi.org:10.1126/science.aam9080

13 Little, D. et al. A single cell high content assay detects mitochondrial dysfunction in iPSC-derived neurons with mutations in SNCA. Sci Rep 8, 9033 (2018). https://doi.org:10.1038/s41598-018-27058-0

14 Cooper, O. et al. Pharmacological rescue of mitochondrial deficits in iPSC-derived neural cells from patients with familial Parkinson’s disease. Sci Transl Med 4, 141ra190 (2012). https://doi.org:10.1126/scitranslmed.3003985

15 Madabhushi, R., Pan, L. & Tsai, L. H. DNA damage and its links to neurodegeneration. Neuron 83, 266–282 (2014). https://doi.org:10.1016/j.neuron.2014.06.034

16 Vasquez, V. et al. Chromatin-Bound Oxidized alpha-Synuclein Causes Strand Breaks in Neuronal Genomes in in vitro Models of Parkinson’s Disease. J Alzheimers Dis 60, S133–S150 (2017). https://doi.org:10.3233/JAD-170342

17 Piper, D. A., Sastre, D. & Schule, B. Advancing Stem Cell Models of Alpha-Synuclein Gene Regulation in Neurodegenerative Disease. Front Neurosci 12, 199 (2018). https://doi.org:10.3389/fnins.2018.00199

18 Kriks, S. et al. Dopamine neurons derived from human ES cells efficiently engraft in animal models of Parkinson’s disease. Nature 480, 547–551 (2011). https://doi.org:10.1038/nature10648

19 Kim, D., Paggi, J. M., Park, C., Bennett, C. & Salzberg, S. L. Graph-based genome alignment and genotyping with HISAT2 and HISAT-genotype. Nat Biotechnol 37, 907–915 (2019). https://doi.org:10.1038/s41587-019-0201-4

20 Chen, L. et al. Unregulated cytosolic dopamine causes neurodegeneration associated with oxidative stress in mice. J Neurosci 28, 425–433 (2008). https://doi.org:10.1523/JNEUROSCI.3602-07.2008

21 Barzilai, A. et al. The molecular mechanism of dopamine-induced apoptosis: identification and characterization of genes that mediate dopamine toxicity. J Neural Transm Suppl, 59–76 (2000). https://doi.org:10.1007/978-3-7091-6301-6_4

22 Beriault, D. R. & Werstuck, G. H. Detection and quantification of endoplasmic reticulum stress in living cells using the fluorescent compound, Thioflavin T. Biochim Biophys Acta 1833, 2293–2301 (2013). https://doi.org:10.1016/j.bbamcr.2013.05.020

23 Park, J. H. et al. Alpha-synuclein-induced mitochondrial dysfunction is mediated via a sirtuin 3-dependent pathway. Mol Neurodegener 15, 5 (2020). https://doi.org:10.1186/s13024-019-0349-x

24 Kedariti, M. et al. LRRK2 kinase activity regulates GCase level and enzymatic activity differently depending on cell type in Parkinson’s disease. NPJ Parkinsons Dis 8, 92 (2022). https://doi.org:10.1038/s41531-022-00354-3

25 Gegg, M. E. et al. Glucocerebrosidase deficiency in substantia nigra of parkinson disease brains. Ann Neurol 72, 455–463 (2012). https://doi.org:10.1002/ana.23614

26 Thakur, P. et al. Modeling Parkinson’s disease pathology by combination of fibril seeds and alpha-synuclein overexpression in the rat brain. Proc Natl Acad Sci U S A 114, E8284–E8293 (2017). https://doi.org:10.1073/pnas.1710442114

27 Kurz, A. et al. A53T-alpha-synuclein overexpression impairs dopamine signaling and striatal synaptic plasticity in old mice. PLoS One 5, e11464 (2010). https://doi.org:10.1371/journal.pone.0011464

28 Brockmann, K. GBA-Associated Synucleinopathies: Prime Candidates for Alpha-Synuclein Targeting Compounds. Front Cell Dev Biol 8, 562522 (2020). https://doi.org:10.3389/fcell.2020.562522

29 Milenkovic, I., Blumenreich, S. & Futerman, A. H. GBA mutations, glucosylceramide and Parkinson’s disease. Curr Opin Neurobiol 72, 148–154 (2022).https://doi.org:10.1016/j.conb.2021.11.004

30 Schindlbeck, K. A. et al. LRRK2 and GBA Variants Exert Distinct Influences on Parkinson’s Disease-Specific Metabolic Networks. Cereb Cortex 30, 2867–2878 (2020). https://doi.org:10.1093/cercor/bhz280

31 Sidransky, E., Samaddar, T. & Tayebi, N. Mutations in GBA are associated with familial Parkinson disease susceptibility and age at onset. Neurology 73, 1424–1425, author reply 1425-1426 (2009). https://doi.org:10.1212/WNL.0b013e3181b28601

32 Spina, M. B. & Cohen, G. Dopamine turnover and glutathione oxidation: implications for Parkinson disease. Proc Natl Acad Sci U S A 86, 1398–1400 (1989). https://doi.org:10.1073/pnas.86.4.1398

33 Surmeier, D. J. & Sulzer, D. The pathology roadmap in Parkinson disease. Prion 7, 85–91 (2013). https://doi.org:10.4161/pri.23582

34 Bulteau, A. L. et al. Dysfunction of mitochondrial Lon protease and identification of oxidized protein in mouse brain following exposure to MPTP: Implications for Parkinson disease. Free Radic Biol Med 108, 236–246 (2017). https://doi.org:10.1016/j.freeradbiomed.2017.03.036

35 Hoang, T. et al. Neuronal NOS and cyclooxygenase-2 contribute to DNA damage in a mouse model of Parkinson disease. Free Radic Biol Med 47, 1049–1056 (2009). https://doi.org:10.1016/j.freeradbiomed.2009.07.013

36 Neely, M. D., Davison, C. A., Aschner, M. & Bowman, A. B. From the Cover: Manganese and Rotenone-Induced Oxidative Stress Signatures Differ in iPSC-Derived Human Dopamine Neurons. Toxicol Sci 159, 366–379 (2017). https://doi.org:10.1093/toxsci/kfx145

37 Nguyen, H. N. et al. LRRK2 mutant iPSC-derived DA neurons demonstrate increased susceptibility to oxidative stress. Cell Stem Cell 8, 267–280 (2011). https://doi.org:10.1016/j.stem.2011.01.013

38 Byers, B. et al. SNCA triplication Parkinson’s patient’s iPSC-derived DA neurons accumulate alpha-synuclein and are susceptible to oxidative stress. PLoS One 6, e26159 (2011). https://doi.org:10.1371/journal.pone.0026159

39 Dorszewska, J. et al. Oxidative stress factors in Parkinson’s disease. Neural Regen Res 16, 1383–1391 (2021). https://doi.org:10.4103/1673-5374.300980

40 Minakaki, G., Krainc, D. & Burbulla, L. F. The Convergence of Alpha-Synuclein, Mitochondrial, and Lysosomal Pathways in Vulnerability of Midbrain Dopaminergic Neurons in Parkinson’s Disease. Front Cell Dev Biol 8, 580634 (2020). https://doi.org:10.3389/fcell.2020.580634

41 Lieberman, O. J. et al. alpha-Synuclein-Dependent Calcium Entry Underlies Differential Sensitivity of Cultured SN and VTA Dopaminergic Neurons to a Parkinsonian Neurotoxin. eNeuro 4 (2017). https://doi.org:10.1523/ENEURO.0167-17.2017

42 Mattson, M. P. Parkinson’s disease: don’t mess with calcium. J Clin Invest 122, 1195–1198 (2012). https://doi.org:10.1172/JCI62835

43 Bishop, M. W. et al. Hyperexcitable substantia nigra dopamine neurons in PINK1- and HtrA2/Omi-deficient mice. J Neurophysiol 104, 3009–3020 (2010). https://doi.org:10.1152/jn.00466.2010

44 Wang, Y. et al. Noradrenergic lesion of the locus coeruleus increases apomorphine-induced circling behavior and the firing activity of substantia nigra pars reticulata neurons in a rat model of Parkinson’s disease. Brain Res 1310, 189–199 (2010). https://doi.org:10.1016/j.brainres.2009.10.070

45 Hastings, T. G., Lewis, D. A. & Zigmond, M. J. Role of oxidation in the neurotoxic effects of intrastriatal dopamine injections. Proc Natl Acad Sci U S A 93, 1956–1961 (1996). https://doi.org:10.1073/pnas.93.5.1956

46 Masato, A., Plotegher, N., Boassa, D. & Bubacco, L. Impaired dopamine metabolism in Parkinson’s disease pathogenesis. Mol Neurodegener 14, 35 (2019). https://doi.org:10.1186/s13024-019-0332-6

47 Lotharius, J. & Brundin, P. Impaired dopamine storage resulting from alpha-synuclein mutations may contribute to the pathogenesis of Parkinson’s disease. Hum Mol Genet 11, 2395–2407 (2002). https://doi.org:10.1093/hmg/11.20.2395

48 Pifl, C. et al. Is Parkinson’s disease a vesicular dopamine storage disorder? Evidence from a study in isolated synaptic vesicles of human and nonhuman primate striatum. J Neurosci 34, 8210–8218 (2014). https://doi.org:10.1523/JNEUROSCI.5456-13.2014

49 Goldstein, D. S., Kopin, I. J. & Sharabi, Y. Catecholamine autotoxicity. Implications for pharmacology and therapeutics of Parkinson disease and related disorders. Pharmacol Ther 144, 268–282 (2014). https://doi.org:10.1016/j.pharmthera.2014.06.006

50 Virdi, G. S. et al. Protein aggregation and calcium dysregulation are hallmarks of familial Parkinson’s disease in midbrain dopaminergic neurons. NPJ Parkinsons Dis 8, 162 (2022). https://doi.org:10.1038/s41531-022-00423-7

51 Tabata, Y. et al. T-type Calcium Channels Determine the Vulnerability of Dopaminergic Neurons to Mitochondrial Stress in Familial Parkinson Disease. Stem Cell Reports 11, 1171–1184 (2018). https://doi.org:10.1016/j.stemcr.2018.09.006

52 Zampese, E. & Surmeier, D. J. Calcium, Bioenergetics, and Parkinson’s Disease. Cells 9 (2020). https://doi.org:10.3390/cells9092045

53 Dagra, A. et al. alpha-Synuclein-induced dysregulation of neuronal activity contributes to murine dopamine neuron vulnerability. NPJ Parkinsons Dis7, 76 (2021). https://doi.org:10.1038/s41531-021-00210-w

54 Park, S. M. et al. Distinct roles of the N-terminal-binding domain and the C-terminal-solubilizing domain of alpha-synuclein, a molecular chaperone. J Biol Chem 277, 28512–28520 (2002). https://doi.org:10.1074/jbc.M111971200

55 Nath, S., Goodwin, J., Engelborghs, Y. & Pountney, D. L. Raised calcium promotes alpha-synuclein aggregate formation. Mol Cell Neurosci 46, 516–526 (2011). https://doi.org:10.1016/j.mcn.2010.12.004

56 Follett, J., Darlow, B., Wong, M. B., Goodwin, J. & Pountney, D. L. Potassium depolarization and raised calcium induces alpha-synuclein aggregates. Neurotox Res 23, 378–392 (2013). https://doi.org:10.1007/s12640-012-9366-z

57 Martinez, J., Moeller, I., Erdjument-Bromage, H., Tempst, P. & Lauring, B. Parkinson’s disease-associated alpha-synuclein is a calmodulin substrate. J Biol Chem 278, 17379–17387 (2003). https://doi.org:10.1074/jbc.M209020200

58 Liu, J., Liu, M. C. & Wang, K. K. Calpain in the CNS: from synaptic function to neurotoxicity. Sci Signal 1, re1 (2008). https://doi.org:10.1126/stke.114re1

59 Dufty, B. M. et al. Calpain-cleavage of alpha-synuclein: connecting proteolytic processing to disease-linked aggregation. Am J Pathol 170, 1725–1738 (2007). https://doi.org:10.2353/ajpath.2007.061232

60 Maegawa, G. H. et al. Identification and characterization of ambroxol as an enzyme enhancement agent for Gaucher disease. J Biol Chem 284, 23502–23516 (2009). https://doi.org:10.1074/jbc.M109.012393

61 Mullin, S. et al. Ambroxol for the Treatment of Patients With Parkinson Disease With and Without Glucocerebrosidase Gene Mutations: A Nonrandomized, Noncontrolled Trial. JAMA Neurol 77, 427–434 (2020). https://doi.org:10.1001/jamaneurol.2019.4611

62 Surmeier, D. J., Guzman, J. N. & Sanchez-Padilla, J. Calcium, cellular aging, and selective neuronal vulnerability in Parkinson’s disease. Cell Calcium 47, 175–182 (2010). https://doi.org:10.1016/j.ceca.2009.12.003

