## Supplementary material for "Diverging Parkinson’s Disease Pathology between patient-derived *GBA^N370S^, LRRK2^G2019S^* and engineered *SNCA^A53T^* iPSC-derived Dopaminergic Neurons": Table S1

| test_id | gene_id | gene | locus | status | AHN | GBA | log2(fold_change) | test_stat | p_value | q_value | significant |
|---|---|---|---|---|---|---|---|---|---|---|---|
| ENSG00000214548 | ENSG00000214548 | MEG3 | chr14:100779409-101027415 | OK | 0.438388 | 847.363 | 10.9166 | 7.05363 | 5.00E-05 | 0.00107006 | yes |
| ENSG00000225746 | ENSG00000225746 | MEG8 | chr14:100779409-101027415 | OK | 0.165683 | 24.6265 | 7.21564 | 0.851064 | 5.00E-05 | 0.00107006 | yes |
| ENSG00000134184 | ENSG00000134184 | GSTM1 | chr1:109656080-109709551 | OK | 0.00935455 | 0.857726 | 6.5187 | 0.194291 | 5.00E-05 | 0.00107006 | yes |
| ENSG00000140598 | ENSG00000140598 | EFL1 | chr15:82130229-82262763 | OK | 4.14493 | 370.202 | 6.48082 | 17.0574 | 5.00E-05 | 0.00107006 | yes |
| ENSG00000180758 | ENSG00000180758 | GPR157 | chr1:9100304-9129170 | OK | 0.239011 | 11.5787 | 5.59826 | 2.64361 | 5.00E-05 | 0.00107006 | yes |
| ENSG00000227617 | ENSG00000227617 | CERS6-AS1 | chr2:168455861-168913371 | OK | 0.35822 | 11.8827 | 5.05188 | 1.22731 | 5.00E-05 | 0.00107006 | yes |
| ENSG00000268883 | ENSG00000268883 | PNMA6B | chrX:153075768-153076968 | OK | 0.50514 | 15.0891 | 4.90068 | 6.21533 | 5.00E-05 | 0.00107006 | yes |
| ENSG00000109991 | ENSG00000109991 | P2RX3 | chr11:57338373-57370600 | OK | 0.158026 | 3.35138 | 4.40652 | 2.64191 | 5.00E-05 | 0.00107006 | yes |
| ENSG00000240342 | ENSG00000240342 | RPS2P5 | chr12:118149800-118372945 | OK | 8.5599 | 156.933 | 4.19641 | 5.67466 | 5.00E-05 | 0.00107006 | yes |
| ENSG00000146469 | ENSG00000146469 | VIP | chr6:152750797-152759765 | OK | 0.816407 | 11.9835 | 3.87562 | 6.32473 | 5.00E-05 | 0.00107006 | yes |
| ENSG00000223403 | ENSG00000223403 | MEG9 | chr14:101069910-101072937 | OK | 0.388392 | 5.23983 | 3.75394 | 2.86686 | 5.00E-05 | 0.00107006 | yes |
| ENSG00000159247 | ENSG00000159247 | TUBBP5 | chr9:138150074-138179774 | OK | 0.521212 | 6.69712 | 3.6836 | 4.55701 | 5.00E-05 | 0.00107006 | yes |
| ENSG00000102024 | ENSG00000102024 | PLS3 | chrX:115518181-115650861 | OK | 3.47662 | 41.5102 | 3.57771 | 9.62569 | 5.00E-05 | 0.00107006 | yes |
| ENSG00000184470 | ENSG00000184470 | TXNRD2 | chr22:19875516-20016808 | OK | 0.169473 | 1.87508 | 3.46782 | 0.624159 | 5.00E-05 | 0.00107006 | yes |
| ENSG00000183054 | ENSG00000183054 | RGPD6 | chr2:110513811-110577185 | OK | 0.38669 | 4.03061 | 3.38175 | 2.08509 | 5.00E-05 | 0.00107006 | yes |
| ENSG00000183793 | ENSG00000183793 | NPIPA5 | chr16:15363623-15381047 | OK | 1.22716 | 10.8276 | 3.14132 | 4.5175 | 5.00E-05 | 0.00107006 | yes |
| ENSG00000233098 | ENSG00000233098 | CCDC144NL-AS1 | chr17:20814619-21043760 | OK | 4.86986 | 37.621 | 2.94959 | 1.09083 | 5.00E-05 | 0.00107006 | yes |
| ENSG00000250138 | ENSG00000250138 | AC139495.3 | chr5:69631962-69636399 | OK | 0.45568 | 2.83059 | 2.63501 | 4.96257 | 5.00E-05 | 0.00107006 | yes |
| ENSG00000168743 | ENSG00000168743 | NPNT | chr4:105552619-106022478 | OK | 13.1701 | 79.025 | 2.58504 | 7.71083 | 5.00E-05 | 0.00107006 | yes |
| ENSG00000197980 | ENSG00000197980 | LEKR1 | chr3:156825480-157046129 | OK | 0.545148 | 3.01486 | 2.46737 | 2.09708 | 5.00E-05 | 0.00107006 | yes |
| ENSG00000181195 | ENSG00000181195 | PENK | chr8:56436673-56559823 | OK | 0.941177 | 5.14919 | 2.45181 | 2.4619 | 5.00E-05 | 0.00107006 | yes |
| ENSG00000247516 | ENSG00000247516 | MIR4458HG | chr5:8333480-8463095 | OK | 1.17162 | 5.78783 | 2.30451 | 3.23051 | 5.00E-05 | 0.00107006 | yes |
| ENSG00000269226 | ENSG00000269226 | TMSB15B | chrX:104063870-104076212 | OK | 0.515209 | 2.37978 | 2.2076 | 2.82557 | 5.00E-05 | 0.00107006 | yes |
| ENSG00000185324 | ENSG00000185324 | CDK10 | chr16:89680736-89701705 | OK | 11.0655 | 50.852 | 2.20023 | 3.38572 | 5.00E-05 | 0.00107006 | yes |
| ENSG00000160201 | ENSG00000160201 | U2AF1 | chr21:43092955-43107587 | OK | 3.20765 | 14.0112 | 2.12699 | 3.09225 | 5.00E-05 | 0.00107006 | yes |
| ENSG00000108379 | ENSG00000108379 | WNT3 | chr17:46762505-46833154 | OK | 3.36951 | 14.5203 | 2.10747 | 3.28765 | 5.00E-05 | 0.00107006 | yes |
| ENSG00000172058 | ENSG00000172058 | SERF1A | chr5:70900664-70918530 | OK | 0.465201 | 1.99803 | 2.10265 | 1.74209 | 5.00E-05 | 0.00107006 | yes |
| ENSG00000281181 | ENSG00000281181 | FP236383.3 | chr21:8437628-8438551 | OK | 16.0923 | 69.0772 | 2.10184 | 6.52903 | 5.00E-05 | 0.00107006 | yes |
| ENSG00000177551 | ENSG00000177551 | NHLH2 | chr1:115836376-115843917 | OK | 2.91362 | 12.2713 | 2.0744 | 5.30207 | 5.00E-05 | 0.00107006 | yes |
| ENSG00000152229 | ENSG00000152229 | PSTPIP2 | chr18:45983535-46072272 | OK | 0.412172 | 1.69775 | 2.0423 | 2.11315 | 5.00E-05 | 0.00107006 | yes |
| ENSG00000166575 | ENSG00000166575 | TMEM135 | chr11:87037843-87324359 | OK | 9.26877 | 36.2252 | 1.96654 | 8.52762 | 5.00E-05 | 0.00107006 | yes |
| ENSG00000276791 | ENSG00000276791 | AC092117.1 | chr16:2777318-2780568 | OK | 0.830558 | 3.20919 | 1.95006 | 3.90197 | 5.00E-05 | 0.00107006 | yes |
| ENSG00000249859 | ENSG00000249859 | PVT1 | chr8:127794532-128101253 | OK | 1.84958 | 6.85347 | 1.88963 | 1.62298 | 5.00E-05 | 0.00107006 | yes |
| ENSG00000142102 | ENSG00000142102 | PGGHG | chr11:289134-296107 | OK | 1.1411 | 3.88375 | 1.76702 | 1.91906 | 5.00E-05 | 0.00107006 | yes |
| ENSG00000253305 | ENSG00000253305 | PCDHGB6 | chr5:141330570-141512981 | OK | 20.647 | 69.1532 | 1.74386 | 3.89652 | 5.00E-05 | 0.00107006 | yes |
| ENSG00000064666 | ENSG00000064666 | CNN2 | chr19:1026580-1039068 | OK | 5.6224 | 18.4721 | 1.71609 | 4.72382 | 5.00E-05 | 0.00107006 | yes |
| ENSG00000227775 | ENSG00000227775 | AL031282.1 | chr1:1702729-1745992 | OK | 6.49717 | 21.2669 | 1.71072 | 2.93163 | 5.00E-05 | 0.00107006 | yes |
| ENSG00000141750 | ENSG00000141750 | STAC2 | chr17:39210535-39225872 | OK | 0.752849 | 2.39782 | 1.67129 | 3.16652 | 5.00E-05 | 0.00107006 | yes |
| ENSG00000125355 | ENSG00000125355 | TMEM255A | chrX:120258649-120311556 | OK | 5.86006 | 18.4103 | 1.65153 | 3.27406 | 5.00E-05 | 0.00107006 | yes |
| ENSG00000160307 | ENSG00000160307 | S100B | chr21:46598961-46605208 | OK | 2.40601 | 7.40235 | 1.62134 | 2.6439 | 5.00E-05 | 0.00107006 | yes |
| ENSG00000146674 | ENSG00000146674 | IGFBP3 | chr7:45912244-45921874 | OK | 0.927386 | 2.81642 | 1.60262 | 2.064 | 5.00E-05 | 0.00107006 | yes |
| ENSG00000241494 | ENSG00000241494 | AL355032.1 | chr14:101634453-101732522 | OK | 10.2978 | 31.1025 | 1.5947 | 2.70572 | 5.00E-05 | 0.00107006 | yes |
| ENSG00000100593 | ENSG00000100593 | ISM2 | chr14:77474393-77498850 | OK | 0.558759 | 1.62971 | 1.54431 | 1.78618 | 5.00E-05 | 0.00107006 | yes |
| ENSG00000237550 | ENSG00000237550 | RPL9P9 | chr15:82355141-82439153 | OK | 192.461 | 551.604 | 1.51907 | 6.17423 | 5.00E-05 | 0.00107006 | yes |
| ENSG00000165646 | ENSG00000165646 | SLC18A2 | chr10:117239599-117375467 | OK | 10.6676 | 30.3407 | 1.50802 | 3.69307 | 5.00E-05 | 0.00107006 | yes |
| ENSG00000175701 | ENSG00000175701 | LINC00116 | chr2:110211528-110285856 | OK | 24.838 | 68.7401 | 1.4686 | 3.33279 | 5.00E-05 | 0.00107006 | yes |

| | | | | | | | | | | | |
|---|---|---|---|---|---|---|---|---|---|---|---|
| ENSG00000134443 | ENSG00000134443 | GRP | chr18:59220167-59230774 | OK | 3.69769 | 10.1582 | 1.45795 | 2.38225 | 5.00E-05 | 0.00107006 | yes |
| ENSG00000260075 | ENSG00000260075 | NSFP1 | chr17:46372854-46487141 | OK | 1.77069 | 4.80831 | 1.44122 | 2.57591 | 5.00E-05 | 0.00107006 | yes |
| ENSG00000101198 | ENSG00000101198 | NKAIN4 | chr20:63240783-63272694 | OK | 4.27332 | 11.4118 | 1.41709 | 3.17104 | 5.00E-05 | 0.00107006 | yes |
| ENSG00000005981 | ENSG00000005981 | ASB4 | chr7:95478443-95540232 | OK | 1.12353 | 2.99811 | 1.41601 | 2.37621 | 5.00E-05 | 0.00107006 | yes |
| ENSG00000271824 | ENSG00000271824 | SMIM32 | chr5:136191467-136193134 | OK | 12.7809 | 33.5838 | 1.39377 | 4.83448 | 5.00E-05 | 0.00107006 | yes |
| ENSG00000119138 | ENSG00000119138 | KLF9 | chr9:70384596-70414624 | OK | 0.530614 | 1.35889 | 1.3567 | 2.63848 | 5.00E-05 | 0.00107006 | yes |
| ENSG00000168952 | ENSG00000168952 | STXBP6 | chr14:24809655-25050297 | OK | 1.3362 | 3.38481 | 1.34093 | 2.70027 | 5.00E-05 | 0.00107006 | yes |
| ENSG00000163064 | ENSG00000163064 | EN1 | chr2:118842170-118847678 | OK | 24.6013 | 61.0677 | 1.31167 | 4.60251 | 5.00E-05 | 0.00107006 | yes |
| ENSG00000172247 | ENSG00000172247 | C1QTNF4 | chr11:47589663-47594659 | OK | 24.6829 | 60.6223 | 1.29634 | 5.29835 | 5.00E-05 | 0.00107006 | yes |
| ENSG00000163932 | ENSG00000163932 | PRKCD | chr3:53156008-53192717 | OK | 3.14086 | 7.67768 | 1.28951 | 3.22201 | 5.00E-05 | 0.00107006 | yes |
| ENSG00000121769 | ENSG00000121769 | FABP3 | chr1:31365624-31376850 | OK | 38.9411 | 92.4464 | 1.24732 | 4.29993 | 5.00E-05 | 0.00107006 | yes |
| ENSG00000189343 | ENSG00000189343 | RPS2P46 | chr17:19417803-19492991 | OK | 17.284 | 40.5813 | 1.23138 | 3.69708 | 5.00E-05 | 0.00107006 | yes |
| ENSG00000260941 | ENSG00000260941 | LINC00622 | chr1:119597701-119599271 | OK | 1.58008 | 3.6781 | 1.21896 | 2.16564 | 5.00E-05 | 0.00107006 | yes |
| ENSG00000275993 | ENSG00000275993 | CU639417.1 | chr21:6111133-6123739 | OK | 3.50017 | 8.13005 | 1.21584 | 3.9803 | 5.00E-05 | 0.00107006 | yes |
| ENSG00000088882 | ENSG00000088882 | CPXM1 | chr20:2794068-2800637 | OK | 3.48864 | 8.07714 | 1.21118 | 3.26284 | 5.00E-05 | 0.00107006 | yes |
| ENSG00000006128 | ENSG00000006128 | TAC1 | chr7:97731907-97740472 | OK | 49.5625 | 113.565 | 1.1962 | 4.87939 | 5.00E-05 | 0.00107006 | yes |
| ENSG00000090539 | ENSG00000090539 | CHRD | chr3:184380072-184390736 | OK | 2.17088 | 4.90208 | 1.17512 | 1.67358 | 5.00E-05 | 0.00107006 | yes |
| ENSG00000142173 | ENSG00000142173 | COL6A2 | chr21:46098096-46132849 | OK | 1.8385 | 3.98763 | 1.117 | 2.0976 | 5.00E-05 | 0.00107006 | yes |
| ENSG00000270231 | ENSG00000270231 | NBPF8 | chr1:120436352-120467739 | OK | 2.4384 | 5.28648 | 1.11637 | 3.48646 | 5.00E-05 | 0.00107006 | yes |
| ENSG00000145248 | ENSG00000145248 | SLC10A4 | chr4:48483342-48489196 | OK | 2.44389 | 5.29637 | 1.11583 | 2.37597 | 5.00E-05 | 0.00107006 | yes |
| ENSG00000265972 | ENSG00000265972 | TXNIP | chr1:145992434-145996600 | OK | 4.94491 | 10.6572 | 1.10782 | 3.33781 | 5.00E-05 | 0.00107006 | yes |
| ENSG00000128573 | ENSG00000128573 | FOXP2 | chr7:114086326-114693772 | OK | 9.89369 | 21.2191 | 1.10078 | 1.8293 | 5.00E-05 | 0.00107006 | yes |
| ENSG00000196756 | ENSG00000196756 | SNHG17 | chr20:38420587-38435353 | OK | 14.9028 | 31.8771 | 1.09693 | 2.31949 | 5.00E-05 | 0.00107006 | yes |
| ENSG00000273136 | ENSG00000273136 | NBPF26 | chr1:120723922-120841481 | OK | 1.88521 | 4.00521 | 1.08715 | 2.4941 | 5.00E-05 | 0.00107006 | yes |
| ENSG00000179603 | ENSG00000179603 | GRM8 | chr7:126438597-127253294 | OK | 16.1859 | 34.2436 | 1.0811 | 3.19941 | 5.00E-05 | 0.00107006 | yes |
| ENSG00000104327 | ENSG00000104327 | CALB1 | chr8:90058607-90095475 | OK | 27.7843 | 58.4948 | 1.07404 | 4.48718 | 5.00E-05 | 0.00107006 | yes |
| ENSG00000170430 | ENSG00000170430 | MGMT | chr10:129467183-129768007 | OK | 4.34802 | 9.10728 | 1.06666 | 2.65699 | 5.00E-05 | 0.00107006 | yes |
| ENSG00000226067 | ENSG00000226067 | LINC00623 | chr1:120913150-121052167 | OK | 6.89633 | 14.4349 | 1.06566 | 2.11286 | 5.00E-05 | 0.00107006 | yes |
| ENSG00000279249 | ENSG00000279249 | AC007614.5 | chr16:49341662-49350554 | OK | 0.642771 | 1.32246 | 1.04084 | 2.51597 | 5.00E-05 | 0.00107006 | yes |
| ENSG00000167333 | ENSG00000167333 | TRIM68 | chr11:4598671-4608259 | OK | 3.2239 | 6.59711 | 1.03303 | 2.41145 | 5.00E-05 | 0.00107006 | yes |
| ENSG00000145824 | ENSG00000145824 | CXCL14 | chr5:135559576-135634874 | OK | 25.1484 | 51.4496 | 1.03269 | 4.07104 | 5.00E-05 | 0.00107006 | yes |
| ENSG00000111666 | ENSG00000111666 | CHPT1 | chr12:101696946-101744140 | OK | 7.0751 | 14.3116 | 1.01636 | 2.56181 | 5.00E-05 | 0.00107006 | yes |
| ENSG00000283293 | ENSG00000283293 | RN7SK | chr6:52995619-52995950 | OK | 151.816 | 306.083 | 1.01159 | 2.72699 | 5.00E-05 | 0.00107006 | yes |
| ENSG00000274012 | ENSG00000274012 | RN7SL2 | chr14:49861175-49864379 | OK | 15342.4 | 7658.46 | -1.0024 | -5.13477 | 5.00E-05 | 0.00107006 | yes |
| ENSG00000179455 | ENSG00000179455 | MKRN3 | chr15:23565677-23630075 | OK | 6.88363 | 3.43452 | -1.00306 | -2.52202 | 5.00E-05 | 0.00107006 | yes |
| ENSG00000203875 | ENSG00000203875 | SNHG5 | chr6:85660949-85678736 | OK | 56.0587 | 27.885 | -1.00745 | -2.21776 | 5.00E-05 | 0.00107006 | yes |
| ENSG00000130518 | ENSG00000130518 | KIAA1683 | chr19:18257096-18274509 | OK | 2.89909 | 1.41024 | -1.03965 | -2.12968 | 5.00E-05 | 0.00107006 | yes |
| ENSG00000051523 | ENSG00000051523 | CYBA | chr16:88643282-88651152 | OK | 19.3224 | 9.35456 | -1.04653 | -2.41915 | 5.00E-05 | 0.00107006 | yes |
| ENSG00000119772 | ENSG00000119772 | DNMT3A | chr2:25227854-25342590 | OK | 38.8005 | 18.7559 | -1.04873 | -2.52984 | 5.00E-05 | 0.00107006 | yes |
| ENSG00000267651 | ENSG00000267651 | AC015961.1 | chr18:37234118-37236242 | OK | 3.84234 | 1.85628 | -1.04957 | -2.18656 | 5.00E-05 | 0.00107006 | yes |
| ENSG00000178162 | ENSG00000178162 | FAR2P2 | chr2:130416754-130441413 | OK | 59.4296 | 28.4792 | -1.06127 | -2.30478 | 5.00E-05 | 0.00107006 | yes |
| ENSG00000036565 | ENSG00000036565 | SLC18A1 | chr8:20144854-20183206 | OK | 6.93097 | 3.2533 | -1.09115 | -2.38739 | 5.00E-05 | 0.00107006 | yes |
| ENSG00000227888 | ENSG00000227888 | FAM66A | chr8:12362018-12388296 | OK | 19.0138 | 8.77228 | -1.11602 | -2.57094 | 5.00E-05 | 0.00107006 | yes |
| ENSG00000272501 | ENSG00000272501 | AL662844.4 | chr6:31195199-31203968 | OK | 6.8669 | 3.16017 | -1.11966 | -2.98998 | 5.00E-05 | 0.00107006 | yes |
| ENSG00000187244 | ENSG00000187244 | BCAM | chr19:44809058-44821420 | OK | 16.6344 | 7.555 | -1.13867 | -3.08767 | 5.00E-05 | 0.00107006 | yes |
| ENSG00000154162 | ENSG00000154162 | CDH12 | chr5:21616261-22853622 | OK | 2.87732 | 1.29345 | -1.15351 | -1.99946 | 5.00E-05 | 0.00107006 | yes |
| ENSG00000144747 | ENSG00000144747 | TMF1 | chr3:68975213-69080408 | OK | 11.3396 | 5.06036 | -1.16405 | -3.00442 | 5.00E-05 | 0.00107006 | yes |

| | | | | | | | | | | | | |
|---|---|---|---|---|---|---|---|---|---|---|---|---|
| ENSG00000107731 | ENSG00000107731 | UNC5B | chr10:71212569-71302864 | OK | 3.57957 | 1.58858 | | -1.17205 | -3.39399 | 5.00E-05 | 0.00107006 | yes |
| ENSG00000064300 | ENSG00000064300 | NGFR | chr17:49494222-49583827 | OK | 5.57888 | 2.46776 | | -1.17677 | -2.39952 | 5.00E-05 | 0.00107006 | yes |
| ENSG00000115461 | ENSG00000115461 | IGFBP5 | chr2:216672104-216994079 | OK | 3.78436 | 1.65009 | | -1.19751 | -2.92152 | 5.00E-05 | 0.00107006 | yes |
| ENSG00000118523 | ENSG00000118523 | CTGF | chr6:131948175-132077393 | OK | 4.20005 | 1.8031 | | -1.21993 | -2.59437 | 5.00E-05 | 0.00107006 | yes |
| ENSG00000135549 | ENSG00000135549 | PKIB | chr6:122471916-122726373 | OK | 71.0882 | 30.3575 | | -1.22756 | -3.7793 | 5.00E-05 | 0.00107006 | yes |
| ENSG00000205922 | ENSG00000205922 | ONECUT3 | chr19:1752372-1780988 | OK | 3.70692 | 1.56971 | | -1.23973 | -3.90461 | 5.00E-05 | 0.00107006 | yes |
| ENSG00000116147 | ENSG00000116147 | TNR | chr1:175307217-175743770 | OK | 6.15185 | 2.58835 | | -1.24899 | -2.63278 | 5.00E-05 | 0.00107006 | yes |
| ENSG00000152402 | ENSG00000152402 | GUCY1A2 | chr11:106674011-107018524 | OK | 4.13469 | 1.73694 | | -1.25123 | -3.60651 | 5.00E-05 | 0.00107006 | yes |
| ENSG00000239268 | ENSG00000239268 | AC092691.1 | chr3:117672153-117997592 | OK | 105.179 | 44.0405 | | -1.25594 | -3.88854 | 5.00E-05 | 0.00107006 | yes |
| ENSG00000148143 | ENSG00000148143 | ZNF462 | chr9:106863096-107102988 | OK | 18.3992 | 7.63044 | | -1.26981 | -1.97658 | 5.00E-05 | 0.00107006 | yes |
| ENSG00000183323 | ENSG00000183323 | CCDC125 | chr5:69280174-69332809 | OK | 3.84288 | 1.59106 | | -1.2722 | -2.64831 | 5.00E-05 | 0.00107006 | yes |
| ENSG00000255495 | ENSG00000255495 | AC145124.1 | chr8:12194466-12196280 | OK | 5.5955 | 2.30696 | | -1.27827 | -2.44157 | 5.00E-05 | 0.00107006 | yes |
| ENSG00000232599 | ENSG00000232599 | AL008707.1 | chrX:125203804-125204338 | OK | 46.8613 | 19.2703 | | -1.28202 | -3.02948 | 5.00E-05 | 0.00107006 | yes |
| ENSG00000120738 | ENSG00000120738 | EGR1 | chr5:138465489-138469315 | OK | 4.36579 | 1.78081 | | -1.29371 | -3.10988 | 5.00E-05 | 0.00107006 | yes |
| ENSG00000261167 | ENSG00000261167 | AC107027.3 | chr3:131455125-131458598 | OK | 2.46159 | 0.990667 | | -1.31312 | -2.71602 | 5.00E-05 | 0.00107006 | yes |
| ENSG00000180229 | ENSG00000180229 | HERC2P3 | chr15:20379494-20506180 | OK | 21.7597 | 8.6854 | | -1.325 | -2.52051 | 5.00E-05 | 0.00107006 | yes |
| ENSG00000129317 | ENSG00000129317 | PUS7L | chr12:43718992-43758817 | OK | 4.87255 | 1.931 | | -1.33533 | -2.79977 | 5.00E-05 | 0.00107006 | yes |
| ENSG00000189060 | ENSG00000189060 | H1F0 | chr22:37805092-37807436 | OK | 41.16 | 16.3077 | | -1.33569 | -5.44541 | 5.00E-05 | 0.00107006 | yes |
| ENSG00000143320 | ENSG00000143320 | CRABP2 | chr1:156699605-156705816 | OK | 34.9508 | 13.8413 | | -1.33635 | -3.20912 | 5.00E-05 | 0.00107006 | yes |
| ENSG00000164330 | ENSG00000164330 | EBF1 | chr5:158695915-159099761 | OK | 21.2704 | 8.39661 | | -1.34097 | -2.8406 | 5.00E-05 | 0.00107006 | yes |
| ENSG00000135333 | ENSG00000135333 | EPHA7 | chr6:93240019-93419547 | OK | 2.28333 | 0.901151 | | -1.3413 | -2.91592 | 5.00E-05 | 0.00107006 | yes |
| ENSG00000181215 | ENSG00000181215 | C4orf50 | chr4:5897372-6018507 | OK | 1.95024 | 0.761385 | | -1.35695 | -3.54654 | 5.00E-05 | 0.00107006 | yes |
| ENSG00000184216 | ENSG00000184216 | IRAK1 | chrX:154010499-154019989 | OK | 22.1872 | 8.60496 | | -1.36649 | -3.24487 | 5.00E-05 | 0.00107006 | yes |
| ENSG00000090857 | ENSG00000090857 | PDPR | chr16:70113625-70187361 | OK | 9.62574 | 3.70643 | | -1.37687 | -4.10385 | 5.00E-05 | 0.00107006 | yes |
| ENSG00000161681 | ENSG00000161681 | SHANK1 | chr19:50661826-50719450 | OK | 58.5191 | 22.1727 | | -1.40012 | -2.36004 | 5.00E-05 | 0.00107006 | yes |
| ENSG00000214279 | ENSG00000214279 | SCART1 | chr10:133453927-133569835 | OK | 2.27746 | 0.833909 | | -1.44947 | -2.47053 | 5.00E-05 | 0.00107006 | yes |
| ENSG00000185551 | ENSG00000185551 | NR2F2 | chr15:95990581-96340263 | OK | 46.1361 | 16.8472 | | -1.45338 | -5.37037 | 5.00E-05 | 0.00107006 | yes |
| ENSG00000276550 | ENSG00000276550 | HERC2P2 | chr15:22495569-22590815 | OK | 29.0157 | 10.4997 | | -1.46649 | -2.4646 | 5.00E-05 | 0.00107006 | yes |
| ENSG00000198914 | ENSG00000198914 | POU3F3 | chr2:104853284-104926052 | OK | 4.98329 | 1.76829 | | -1.49475 | -3.54915 | 5.00E-05 | 0.00107006 | yes |
| ENSG00000198680 | ENSG00000198680 | TUSC1 | chr9:25676388-25678440 | OK | 17.5661 | 6.2233 | | -1.49704 | -4.6459 | 5.00E-05 | 0.00107006 | yes |
| ENSG00000109846 | ENSG00000109846 | CRYAB | chr11:111908564-111926872 | OK | 6.89256 | 2.42025 | | -1.50989 | -1.83681 | 5.00E-05 | 0.00107006 | yes |
| ENSG00000141756 | ENSG00000141756 | FKBP10 | chr17:41812679-41823217 | OK | 4.17825 | 1.44869 | | -1.52815 | -2.08214 | 5.00E-05 | 0.00107006 | yes |
| ENSG00000080224 | ENSG00000080224 | EPHA6 | chr3:96814580-97752460 | OK | 5.25375 | 1.81822 | | -1.53082 | -2.83751 | 5.00E-05 | 0.00107006 | yes |
| ENSG00000055332 | ENSG00000055332 | EIF2AK2 | chr2:37084450-37157065 | OK | 5.0962 | 1.71305 | | -1.57286 | -2.10054 | 5.00E-05 | 0.00107006 | yes |
| ENSG00000204525 | ENSG00000204525 | HLA-C | chr6:31268748-31357637 | OK | 13.7539 | 4.505 | | -1.61024 | -2.43357 | 5.00E-05 | 0.00107006 | yes |
| ENSG00000167178 | ENSG00000167178 | ISLR2 | chr15:74100310-74138540 | OK | 65.5525 | 21.4457 | | -1.61196 | -5.72213 | 5.00E-05 | 0.00107006 | yes |
| ENSG00000100867 | ENSG00000100867 | DHRS2 | chr14:23630114-23645639 | OK | 5.44084 | 1.76086 | | -1.62755 | -2.80722 | 5.00E-05 | 0.00107006 | yes |
| ENSG00000118596 | ENSG00000118596 | SLC16A7 | chr12:59596066-59789855 | OK | 1.6148 | 0.520419 | | -1.63361 | -1.23527 | 5.00E-05 | 0.00107006 | yes |
| ENSG00000267313 | ENSG00000267313 | KC6 | chr18:41465782-41632185 | OK | 9.72308 | 3.10206 | | -1.64818 | -2.32633 | 5.00E-05 | 0.00107006 | yes |
| ENSG00000284471 | ENSG00000284471 | CR769775.4 | chr9:61982795-61986934 | OK | 1.42348 | 0.452682 | | -1.65285 | -2.88808 | 5.00E-05 | 0.00107006 | yes |
| ENSG00000159450 | ENSG00000159450 | TCHH | chr1:152106316-152115454 | OK | 0.815336 | 0.258438 | | -1.65757 | -2.82036 | 5.00E-05 | 0.00107006 | yes |
| ENSG00000105894 | ENSG00000105894 | PTN | chr7:137227340-137343865 | OK | 10.8429 | 3.39484 | | -1.67533 | -3.94605 | 5.00E-05 | 0.00107006 | yes |
| ENSG00000113209 | ENSG00000113209 | PCDHB5 | chr5:141100241-141249365 | OK | 14.0212 | 4.2734 | | -1.71415 | -2.46946 | 5.00E-05 | 0.00107006 | yes |
| ENSG00000180787 | ENSG00000180787 | ZFP3 | chr17:5078247-5096374 | OK | 0.617086 | 0.182977 | | -1.75381 | -2.4627 | 5.00E-05 | 0.00107006 | yes |
| ENSG00000146858 | ENSG00000146858 | ZC3HAV1L | chr7:139025705-139036029 | OK | 2.05597 | 0.603666 | | -1.768 | -2.42018 | 5.00E-05 | 0.00107006 | yes |
| ENSG00000100234 | ENSG00000100234 | TIMP3 | chr22:32512551-33058372 | OK | 1.48087 | 0.429615 | | -1.78533 | -2.41956 | 5.00E-05 | 0.00107006 | yes |
| ENSG00000198467 | ENSG00000198467 | TPM2 | chr9:35681991-35691020 | OK | 4.28185 | 1.23491 | | -1.79383 | -2.31032 | 5.00E-05 | 0.00107006 | yes |

| | | | | | | | | | | | | |
|---|---|---|---|---|---|---|---|---|---|---|---|---|
| ENSG00000160200 | ENSG00000160200 | CBS | chr21:43053190-43076943 | OK | 15.9844 | 4.49078 | | -1.83163 | -4.08661 | 5.00E-05 | 0.00107006 | yes |
| ENSG00000159674 | ENSG00000159674 | SPON2 | chr4:1166931-1208962 | OK | 3.39165 | 0.948698 | | -1.83797 | -1.64134 | 5.00E-05 | 0.00107006 | yes |
| ENSG00000088827 | ENSG00000088827 | SIGLEC1 | chr20:3686969-3707128 | OK | 1.41649 | 0.394975 | | -1.84249 | -3.31624 | 5.00E-05 | 0.00107006 | yes |
| ENSG00000254872 | ENSG00000254872 | AC139749.1 | chr11:1049879-1055749 | OK | 1.0219 | 0.28071 | | -1.86411 | -2.06412 | 5.00E-05 | 0.00107006 | yes |
| ENSG00000228223 | ENSG00000228223 | HCG11 | chr6:26521708-26527404 | OK | 0.794949 | 0.215221 | | -1.88505 | -2.8851 | 5.00E-05 | 0.00107006 | yes |
| ENSG00000176438 | ENSG00000176438 | SYNE3 | chr14:95407265-95475836 | OK | 0.952342 | 0.25721 | | -1.88853 | -2.23472 | 5.00E-05 | 0.00107006 | yes |
| ENSG00000135828 | ENSG00000135828 | RNASEL | chr1:182573633-182589256 | OK | 2.13484 | 0.57024 | | -1.90449 | -3.66391 | 5.00E-05 | 0.00107006 | yes |
| ENSG00000130635 | ENSG00000130635 | COL5A1 | chr9:134641773-134844843 | OK | 0.747171 | 0.198192 | | -1.91454 | -1.15477 | 5.00E-05 | 0.00107006 | yes |
| ENSG00000284700 | ENSG00000284700 | AL049637.2 | chr1:50423608-50425316 | OK | 5.37327 | 1.35529 | | -1.9872 | -2.75058 | 5.00E-05 | 0.00107006 | yes |
| ENSG00000168542 | ENSG00000168542 | COL3A1 | chr2:188974319-189012746 | OK | 1.58018 | 0.395871 | | -1.99699 | -1.44197 | 5.00E-05 | 0.00107006 | yes |
| ENSG00000237161 | ENSG00000237161 | AC068446.1 | chr15:21293652-21295201 | OK | 6.31329 | 1.57649 | | -2.00168 | -3.6697 | 5.00E-05 | 0.00107006 | yes |
| ENSG00000279672 | ENSG00000279672 | AP006621.5 | chr11:777577-784297 | OK | 12.7881 | 3.15087 | | -2.02098 | -3.44099 | 5.00E-05 | 0.00107006 | yes |
| ENSG00000161509 | ENSG00000161509 | GRIN2C | chr17:74842022-74861504 | OK | 0.920559 | 0.224785 | | -2.03396 | -2.00245 | 5.00E-05 | 0.00107006 | yes |
| ENSG00000255052 | ENSG00000255052 | FAM66D | chr8:12115781-12177550 | OK | 5.09386 | 1.20513 | | -2.07957 | -2.70786 | 5.00E-05 | 0.00107006 | yes |
| ENSG00000133433 | ENSG00000133433 | GSTT2B | chr22:23957413-23961186 | OK | 12.6087 | 2.92298 | | -2.10891 | -4.02246 | 5.00E-05 | 0.00107006 | yes |
| ENSG00000163219 | ENSG00000163219 | ARHGAP25 | chr2:68679600-68826833 | OK | 1.16437 | 0.259995 | | -2.163 | -1.0094 | 5.00E-05 | 0.00107006 | yes |
| ENSG00000145945 | ENSG00000145945 | FAM50B | chr6:3831932-3855737 | OK | 4.14553 | 0.904464 | | -2.19642 | -3.59981 | 5.00E-05 | 0.00107006 | yes |
| ENSG00000113140 | ENSG00000113140 | SPARC | chr5:151661095-151724782 | OK | 14.1543 | 3.08698 | | -2.19697 | -3.67276 | 5.00E-05 | 0.00107006 | yes |
| ENSG00000132688 | ENSG00000132688 | NES | chr1:156668762-156677397 | OK | 12.5229 | 2.72081 | | -2.20246 | -7.56692 | 5.00E-05 | 0.00107006 | yes |
| ENSG00000236682 | ENSG00000236682 | AC068282.1 | chr2:127389129-127400580 | OK | 1.68256 | 0.363466 | | -2.21077 | -3.16635 | 5.00E-05 | 0.00107006 | yes |
| ENSG00000173698 | ENSG00000173698 | ADGRG2 | chrX:18989308-19122637 | OK | 4.6952 | 1.01348 | | -2.21187 | -5.34206 | 5.00E-05 | 0.00107006 | yes |
| ENSG00000042493 | ENSG00000042493 | CAPG | chr2:85394747-85418432 | OK | 0.859008 | 0.177048 | | -2.27853 | -0.609566 | 5.00E-05 | 0.00107006 | yes |
| ENSG00000019582 | ENSG00000019582 | CD74 | chr5:150401636-150412929 | OK | 4.49015 | 0.921785 | | -2.28426 | -2.40388 | 5.00E-05 | 0.00107006 | yes |
| ENSG00000178394 | ENSG00000178394 | HTR1A | chr5:63957892-63981043 | OK | 1.7432 | 0.354754 | | -2.29684 | -2.00387 | 5.00E-05 | 0.00107006 | yes |
| ENSG00000260409 | ENSG00000260409 | AC012414.5 | chr15:20729746-20756183 | OK | 5.52346 | 1.12335 | | -2.29777 | -3.95692 | 5.00E-05 | 0.00107006 | yes |
| ENSG00000268350 | ENSG00000268350 | FAM156A | chrX:52926401-52995472 | OK | 3.61638 | 0.710854 | | -2.34692 | -1.93034 | 5.00E-05 | 0.00107006 | yes |
| ENSG00000164692 | ENSG00000164692 | COL1A2 | chr7:94394560-94431232 | OK | 2.74769 | 0.539733 | | -2.3479 | -1.82372 | 5.00E-05 | 0.00107006 | yes |
| ENSG00000247809 | ENSG00000247809 | NR2F2-AS1 | chr15:95990581-96340263 | OK | 3.32413 | 0.603461 | | -2.46164 | -0.773014 | 5.00E-05 | 0.00107006 | yes |
| ENSG00000284610 | ENSG00000284610 | AC107918.4 | chr8:12006187-12015193 | OK | 3.53184 | 0.6346 | | -2.4765 | -3.21195 | 5.00E-05 | 0.00107006 | yes |
| ENSG00000275496 | ENSG00000275496 | CU633906.1 | chr21:6228965-6267317 | OK | 4.57415 | 0.78985 | | -2.53385 | -2.33447 | 5.00E-05 | 0.00107006 | yes |
| ENSG00000018280 | ENSG00000018280 | SLC11A1 | chr2:218382028-218396894 | OK | 0.717922 | 0.122704 | | -2.54865 | -0.557671 | 5.00E-05 | 0.00107006 | yes |
| ENSG00000259316 | ENSG00000259316 | AC087632.1 | chr15:64165516-64387687 | OK | 0.884922 | 0.131512 | | -2.75036 | -0.352895 | 5.00E-05 | 0.00107006 | yes |
| ENSG00000269994 | ENSG00000269994 | AL513318.2 | chr9:87008454-87042126 | OK | 2.61143 | 0.387637 | | -2.75206 | -3.6874 | 5.00E-05 | 0.00107006 | yes |
| ENSG00000184956 | ENSG00000184956 | MUC6 | chr11:1012820-1036706 | OK | 1.57235 | 0.233251 | | -2.75296 | -1.9428 | 5.00E-05 | 0.00107006 | yes |
| ENSG00000157227 | ENSG00000157227 | MMP14 | chr14:22836556-22849027 | OK | 1.56984 | 0.222188 | | -2.82077 | -1.41666 | 5.00E-05 | 0.00107006 | yes |
| ENSG00000125730 | ENSG00000125730 | C3 | chr19:6677703-6737603 | OK | 2.34916 | 0.329685 | | -2.83298 | -0.97236 | 5.00E-05 | 0.00107006 | yes |
| ENSG00000275895 | ENSG00000275895 | U2AF1L5 | chr21:6484622-6499261 | OK | 11.8692 | 1.5048 | | -2.97959 | -3.53446 | 5.00E-05 | 0.00107006 | yes |
| ENSG00000215374 | ENSG00000215374 | FAM66B | chr8:7301610-7355354 | OK | 3.29175 | 0.407992 | | -3.01224 | -3.57017 | 5.00E-05 | 0.00107006 | yes |
| ENSG00000117983 | ENSG00000117983 | MUC5B | chr11:1223065-1262172 | OK | 0.8443 | 0.0949708 | | -3.1522 | -0.903215 | 5.00E-05 | 0.00107006 | yes |
| ENSG00000134954 | ENSG00000134954 | ETS1 | chr11:128458760-128587558 | OK | 0.868997 | 0.0950843 | | -3.19207 | -1.01956 | 5.00E-05 | 0.00107006 | yes |
| ENSG00000163046 | ENSG00000163046 | ANKRD30BL | chr2:132147590-132257969 | OK | 2.16164 | 0.233644 | | -3.20974 | -2.44849 | 5.00E-05 | 0.00107006 | yes |
| ENSG00000171848 | ENSG00000171848 | RRM2 | chr2:10120697-10211725 | OK | 0.82652 | 0.0862116 | | -3.2611 | -0.598533 | 5.00E-05 | 0.00107006 | yes |
| ENSG00000247765 | ENSG00000247765 | AC068446.2 | chr15:21298232-21325241 | OK | 3.17538 | 0.297549 | | -3.41573 | -4.4101 | 5.00E-05 | 0.00107006 | yes |
| ENSG00000167608 | ENSG00000167608 | TMC4 | chr19:54160107-54173250 | OK | 1.15111 | 0.105881 | | -3.4425 | -0.825333 | 5.00E-05 | 0.00107006 | yes |
| ENSG00000104879 | ENSG00000104879 | CKM | chr19:45306413-45322977 | OK | 2.93067 | 0.257511 | | -3.50852 | -3.73339 | 5.00E-05 | 0.00107006 | yes |
| ENSG00000176728 | ENSG00000176728 | TTTY14 | chrY:18872500-19077416 | OK | 13.6599 | 1.06789 | | -3.67711 | -3.33736 | 5.00E-05 | 0.00107006 | yes |
| ENSG00000105825 | ENSG00000105825 | TFPI2 | chr7:93591572-93911265 | OK | 5.16122 | 0.399739 | | -3.69058 | -3.18432 | 5.00E-05 | 0.00107006 | yes |

| | | | | | | | | | | | |
|---|---|---|---|---|---|---|---|---|---|---|---|
| ENSG00000230333 | ENSG00000230333 | AC004160.1 | chr7:11180901-11832198 | OK | 1.36568 | 0.102144 | -3.74094 | -0.567387 | 5.00E-05 | 0.00107006 | yes |
| ENSG00000143632 | ENSG00000143632 | ACTA1 | chr1:229431244-229434098 | OK | 5.29051 | 0.347956 | -3.92643 | -3.97171 | 5.00E-05 | 0.00107006 | yes |
| ENSG00000131398 | ENSG00000131398 | KCNC3 | chr19:50311936-50382982 | OK | 10.8111 | 0.619515 | -4.12523 | -5.41115 | 5.00E-05 | 0.00107006 | yes |
| ENSG00000142621 | ENSG00000142621 | FHAD1 | chr1:15247271-15400283 | OK | 1.01223 | 0.0564049 | -4.16557 | -0.44115 | 5.00E-05 | 0.00107006 | yes |
| ENSG00000124333 | ENSG00000124333 | VAMP7 | chrX:155881292-155943769 | OK | 5.01018 | 0.263176 | -4.25076 | -2.85585 | 5.00E-05 | 0.00107006 | yes |
| ENSG00000188153 | ENSG00000188153 | COL4A5 | chrX:108439843-108697545 | OK | 0.640927 | 0.0285944 | -4.48636 | -0.367228 | 5.00E-05 | 0.00107006 | yes |
| ENSG00000231473 | ENSG00000231473 | LINC00441 | chr13:48296512-48303661 | OK | 9.10271 | 0.357346 | -4.6709 | -3.43849 | 5.00E-05 | 0.00107006 | yes |
| ENSG00000205663 | ENSG00000205663 | FAM239B | chrX:3891437-3920746 | OK | 7.98673 | 0.282996 | -4.81875 | -3.77139 | 5.00E-05 | 0.00107006 | yes |
| ENSG00000283196 | ENSG00000283196 | AC006453.2 | chr2:89533184-89585652 | OK | 5.48502 | 0.186652 | -4.87708 | -3.06429 | 5.00E-05 | 0.00107006 | yes |
| ENSG00000127399 | ENSG00000127399 | LRRC61 | chr7:150243915-150338150 | OK | 12.71 | 0.404072 | -4.97521 | -4.81036 | 5.00E-05 | 0.00107006 | yes |
| ENSG00000108821 | ENSG00000108821 | COL1A1 | chr17:50183288-50201632 | OK | 4.83775 | 0.123841 | -5.28778 | -1.32384 | 5.00E-05 | 0.00107006 | yes |
| ENSG00000230021 | ENSG00000230021 | AL669831.3 | chr1:586070-859446 | OK | 8.34561 | 0.176426 | -5.56388 | -0.815029 | 5.00E-05 | 0.00107006 | yes |
| ENSG00000188372 | ENSG00000188372 | ZP3 | chr7:76389333-76442071 | OK | 0.0998041 | 5.69088 | 5.83341 | 1.45766 | 0.0001 | 0.0019628 | yes |
| ENSG00000245750 | ENSG00000245750 | DRAIC | chr15:69463025-69571440 | OK | 1.17945 | 4.43745 | 1.91162 | 2.41044 | 0.0001 | 0.0019628 | yes |
| ENSG00000187091 | ENSG00000187091 | PLCD1 | chr3:38007495-38029762 | OK | 4.87008 | 9.99349 | 1.03704 | 2.00747 | 0.0001 | 0.0019628 | yes |
| ENSG00000156052 | ENSG00000156052 | GNAQ | chr9:77716086-78031458 | OK | 83.4128 | 40.8336 | -1.03051 | -2.24008 | 0.0001 | 0.0019628 | yes |
| ENSG00000071537 | ENSG00000071537 | SEL1L | chr14:81471548-81533861 | OK | 34.8574 | 16.9078 | -1.04378 | -2.06065 | 0.0001 | 0.0019628 | yes |
| ENSG00000214425 | ENSG00000214425 | LRRC37A4P | chr17:45506740-45563230 | OK | 2.95142 | 1.35469 | -1.12345 | -2.2818 | 0.0001 | 0.0019628 | yes |
| ENSG00000080854 | ENSG00000080854 | IGSF9B | chr11:133908563-133956985 | OK | 18.6022 | 8.24242 | -1.17433 | -1.88167 | 0.0001 | 0.0019628 | yes |
| ENSG00000011028 | ENSG00000011028 | MRC2 | chr17:62626030-62693597 | OK | 1.78255 | 0.486031 | -1.87482 | -1.49726 | 0.0001 | 0.0019628 | yes |
| ENSG00000153404 | ENSG00000153404 | PLEKHG4B | chr5:92150-189972 | OK | 0.567035 | 0.145422 | -1.9632 | -1.71071 | 0.0001 | 0.0019628 | yes |
| ENSG00000259905 | ENSG00000259905 | PWRN1 | chr15:24493136-24652130 | OK | 1.70109 | 0.299522 | -2.50573 | -1.2396 | 0.0001 | 0.0019628 | yes |
| ENSG00000198125 | ENSG00000198125 | MB | chr22:35606763-35637951 | OK | 2.43205 | 0.363279 | -2.74302 | -1.39079 | 0.0001 | 0.0019628 | yes |
| ENSG00000282458 | ENSG00000282458 | WASH5P | chr19:60950-71626 | OK | 3.91996 | 0.297044 | -3.72209 | -1.44635 | 0.0001 | 0.0019628 | yes |
| ENSG00000113805 | ENSG00000113805 | CNTN3 | chr3:74262567-74521140 | OK | 0.227378 | 1.52208 | 2.74288 | 1.93675 | 0.00015 | 0.00276892 | yes |
| ENSG00000157445 | ENSG00000157445 | CACNA2D3 | chr3:54122546-55074557 | OK | 2.69052 | 7.05596 | 1.39096 | 2.41311 | 0.00015 | 0.00276892 | yes |
| ENSG00000229119 | ENSG00000229119 | AC026403.1 | chr5:166382304-166382599 | OK | 43.203 | 110.274 | 1.35189 | 2.16797 | 0.00015 | 0.00276892 | yes |
| ENSG00000234745 | ENSG00000234745 | HLA-B | chr6:31268748-31357637 | OK | 8.64657 | 19.415 | 1.16697 | 1.50762 | 0.00015 | 0.00276892 | yes |
| ENSG00000196172 | ENSG00000196172 | ZNF681 | chr19:23739194-23758891 | OK | 0.742006 | 1.5959 | 1.10487 | 1.43437 | 0.00015 | 0.00276892 | yes |
| ENSG00000157617 | ENSG00000157617 | C2CD2 | chr21:41885111-41953890 | OK | 2.09068 | 1.02055 | -1.03463 | -1.66091 | 0.00015 | 0.00276892 | yes |
| ENSG00000141569 | ENSG00000141569 | TRIM65 | chr17:75880334-75897003 | OK | 3.21727 | 1.50563 | -1.09547 | -1.78264 | 0.00015 | 0.00276892 | yes |
| ENSG00000154856 | ENSG00000154856 | APCDD1 | chr18:10454627-10489948 | OK | 11.1391 | 4.5468 | -1.29271 | -2.19437 | 0.00015 | 0.00276892 | yes |
| ENSG00000180422 | ENSG00000180422 | LINC00304 | chr16:89159145-89164245 | OK | 2.64424 | 1.00171 | -1.40039 | -2.06371 | 0.00015 | 0.00276892 | yes |
| ENSG00000269001 | ENSG00000269001 | AC092070.2 | chr19:53197110-53214522 | OK | 2.30431 | 0.729971 | -1.65843 | -1.58245 | 0.00015 | 0.00276892 | yes |
| ENSG00000196071 | ENSG00000196071 | OR2L13 | chr1:248095183-248101103 | OK | 1.75866 | 0.554975 | -1.66398 | -2.28103 | 0.00015 | 0.00276892 | yes |
| ENSG00000255085 | ENSG00000255085 | AF186192.2 | chr8:144700352-144708517 | OK | 8.52748 | 2.51651 | -1.7607 | -2.35386 | 0.00015 | 0.00276892 | yes |
| ENSG00000267368 | ENSG00000267368 | UPK3BL1 | chr7:102482444-102679295 | OK | 15.2992 | 3.77145 | -2.02027 | -2.37857 | 0.00015 | 0.00276892 | yes |
| ENSG00000275395 | ENSG00000275395 | FCGBP | chr19:39863322-39906323 | OK | 0.672428 | 0.110422 | -2.60635 | -2.67853 | 0.00015 | 0.00276892 | yes |
| ENSG00000213185 | ENSG00000213185 | FAM24B | chr10:122832148-122879641 | OK | 4.10782 | 0.620237 | -2.72748 | -2.35464 | 0.00015 | 0.00276892 | yes |
| ENSG00000121966 | ENSG00000121966 | CXCR4 | chr2:136114348-136118165 | OK | 1.45131 | 0.177285 | -3.03322 | -2.6782 | 0.00015 | 0.00276892 | yes |
| ENSG00000234665 | ENSG00000234665 | AL512625.3 | chr9:62856998-62900104 | OK | 5.81071 | 0.277574 | -4.38777 | -2.32745 | 0.00015 | 0.00276892 | yes |
| ENSG00000175170 | ENSG00000175170 | FAM182B | chr20:25763465-25868225 | OK | 1.8368 | 6.53357 | 1.83067 | 2.1915 | 0.0002 | 0.00351582 | yes |
| ENSG00000215256 | ENSG00000215256 | DHRS4-AS1 | chr14:23938730-24006408 | OK | 3.02982 | 7.04737 | 1.21785 | 2.09397 | 0.0002 | 0.00351582 | yes |
| ENSG00000135269 | ENSG00000135269 | TES | chr7:116209233-116508541 | OK | 5.89355 | 12.7414 | 1.11231 | 2.3287 | 0.0002 | 0.00351582 | yes |
| ENSG00000186493 | ENSG00000186493 | C5orf38 | chr5:2752130-2755397 | OK | 21.6577 | 45.7091 | 1.0776 | 1.68312 | 0.0002 | 0.00351582 | yes |
| ENSG00000173715 | ENSG00000173715 | C11orf80 | chr11:66744450-66846546 | OK | 12.2271 | 24.4592 | 1.0003 | 1.97533 | 0.0002 | 0.00351582 | yes |
| ENSG00000103485 | ENSG00000103485 | QPRT | chr16:29662978-29698699 | OK | 3.93422 | 1.93 | -1.02747 | -1.95298 | 0.0002 | 0.00351582 | yes |

| | | | | | | | | | | | |
|---|---|---|---|---|---|---|---|---|---|---|---|
| ENSG00000215156 | ENSG00000215156 | AC138409.1 | chr5:34164697-34244796 | OK | 2.76927 | 1.3291 | -1.05905 | -2.12929 | 0.0002 | 0.00351582 | yes |
| ENSG00000262585 | ENSG00000262585 | LINC01979 | chr17:79915251-79926725 | OK | 0.882516 | 0.364709 | -1.27487 | -2.11042 | 0.0002 | 0.00351582 | yes |
| ENSG00000147257 | ENSG00000147257 | GPC3 | chrX:133535744-133985895 | OK | 4.53416 | 1.86717 | -1.27998 | -1.92295 | 0.0002 | 0.00351582 | yes |
| ENSG00000130513 | ENSG00000130513 | GDF15 | chr19:18374730-18389176 | OK | 3.68884 | 1.44639 | -1.35071 | -1.88248 | 0.0002 | 0.00351582 | yes |
| ENSG00000181585 | ENSG00000181585 | TMIE | chr3:46701332-46710886 | OK | 2.30976 | 0.895867 | -1.36638 | -2.15096 | 0.0002 | 0.00351582 | yes |
| ENSG00000041982 | ENSG00000041982 | TNC | chr9:115019577-115118257 | OK | 1.90496 | 0.399066 | -2.25506 | -0.709574 | 0.0002 | 0.00351582 | yes |
| ENSG00000166292 | ENSG00000166292 | TMEM100 | chr17:55719626-55732121 | OK | 0.775039 | 0.128209 | -2.59577 | -1.02429 | 0.0002 | 0.00351582 | yes |
| ENSG00000182578 | ENSG00000182578 | CSF1R | chr5:150053290-150113372 | OK | 0.830789 | 0.0626386 | -3.72936 | -0.702027 | 0.0002 | 0.00351582 | yes |
| ENSG00000176769 | ENSG00000176769 | TCERG1L | chr10:131092390-131311721 | OK | 0.274039 | 1.11449 | 2.02393 | 2.54071 | 0.00025 | 0.00421621 | yes |
| ENSG00000197321 | ENSG00000197321 | SVIL | chr10:29409401-29736781 | OK | 17.0144 | 60.4681 | 1.82941 | 9.50715 | 0.00025 | 0.00421621 | yes |
| ENSG00000274276 | ENSG00000274276 | CBSL | chr21:6444868-6468040 | OK | 4.97476 | 14.8017 | 1.57306 | 1.68285 | 0.00025 | 0.00421621 | yes |
| ENSG00000260769 | ENSG00000260769 | AC007614.4 | chr16:49336682-49338993 | OK | 1.81695 | 3.73939 | 1.04129 | 2.02996 | 0.00025 | 0.00421621 | yes |
| ENSG00000134352 | ENSG00000134352 | IL6ST | chr5:55935094-55994993 | OK | 13.9135 | 6.90106 | -1.01159 | -1.77843 | 0.00025 | 0.00421621 | yes |
| ENSG00000259953 | ENSG00000259953 | AL138756.1 | chr9:112032554-112037730 | OK | 2.95874 | 1.34605 | -1.13625 | -2.00185 | 0.00025 | 0.00421621 | yes |
| ENSG00000145685 | ENSG00000145685 | LHFPL2 | chr5:78485214-78770021 | OK | 1.91504 | 0.846128 | -1.17843 | -1.35495 | 0.00025 | 0.00421621 | yes |
| ENSG00000096060 | ENSG00000096060 | FKBP5 | chr6:35573584-35728583 | OK | 1.04248 | 0.41464 | -1.33009 | -2.12049 | 0.00025 | 0.00421621 | yes |
| ENSG00000223509 | ENSG00000223509 | AC135983.2 | chr15:32519847-32639949 | OK | 5.95528 | 1.5898 | -1.90532 | -2.59809 | 0.00025 | 0.00421621 | yes |
| ENSG00000142347 | ENSG00000142347 | MYO1F | chr19:8520789-8577577 | OK | 0.843072 | 4.62068 | 2.45438 | 1.56579 | 0.0003 | 0.00488215 | yes |
| ENSG00000036448 | ENSG00000036448 | MYOM2 | chr8:2045039-2165552 | OK | 6.76057 | 15.5766 | 1.20416 | 2.15114 | 0.0003 | 0.00488215 | yes |
| ENSG00000093072 | ENSG00000093072 | ADA2 | chr22:17178789-17221989 | OK | 1.01645 | 2.1823 | 1.10231 | 1.69624 | 0.0003 | 0.00488215 | yes |
| ENSG00000205571 | ENSG00000205571 | SMN2 | chr5:70049611-70078522 | OK | 7.22162 | 14.7185 | 1.02724 | 1.88415 | 0.0003 | 0.00488215 | yes |
| ENSG00000198093 | ENSG00000198093 | ZNF649 | chr19:51745171-51905040 | OK | 8.93721 | 3.87278 | -1.20646 | -1.94187 | 0.0003 | 0.00488215 | yes |
| ENSG00000253771 | ENSG00000253771 | TPTE2P1 | chr13:24924676-24968487 | OK | 6.06768 | 2.09157 | -1.53656 | -1.96781 | 0.0003 | 0.00488215 | yes |
| ENSG00000162620 | ENSG00000162620 | LRRIQ3 | chr1:74026014-74198187 | OK | 0.790982 | 0.12637 | -2.64599 | -0.785494 | 0.0003 | 0.00488215 | yes |
| ENSG00000278200 | ENSG00000278200 | LINC01971 | chr17:81480522-81481570 | OK | 0.58324 | 2.77414 | 2.24988 | 2.56375 | 0.00035 | 0.00550383 | yes |
| ENSG00000144331 | ENSG00000144331 | ZNF385B | chr2:179441981-179861505 | OK | 0.422701 | 1.61032 | 1.92964 | 1.24432 | 0.00035 | 0.00550383 | yes |
| ENSG00000154645 | ENSG00000154645 | CHODL | chr21:17901262-18267373 | OK | 0.677455 | 1.46888 | 1.11652 | 1.70913 | 0.00035 | 0.00550383 | yes |
| ENSG00000198948 | ENSG00000198948 | MFAP3L | chr4:169981746-170033031 | OK | 4.42908 | 9.04787 | 1.03057 | 1.68249 | 0.00035 | 0.00550383 | yes |
| ENSG00000230606 | ENSG00000230606 | AC092683.1 | chr2:97416164-97434847 | OK | 15.0651 | 7.13036 | -1.07917 | -2.23652 | 0.00035 | 0.00550383 | yes |
| ENSG00000215458 | ENSG00000215458 | AATBC | chr21:43805757-43812567 | OK | 1.74986 | 0.794886 | -1.13842 | -1.89292 | 0.00035 | 0.00550383 | yes |
| ENSG00000100842 | ENSG00000100842 | EFS | chr14:23356401-23365752 | OK | 0.573515 | 0.0824173 | -2.79881 | -2.23467 | 0.00035 | 0.00550383 | yes |
| ENSG00000259129 | ENSG00000259129 | LINC00648 | chr14:47764953-47795092 | OK | 1.17155 | 0.1621 | -2.85346 | -0.984117 | 0.00035 | 0.00550383 | yes |
| ENSG00000006116 | ENSG00000006116 | CACNG3 | chr16:24255552-24362801 | OK | 0.465885 | 1.16664 | 1.32432 | 2.03212 | 0.0004 | 0.00614885 | yes |
| ENSG00000233198 | ENSG00000233198 | RNF224 | chr9:137227270-137229638 | OK | 2.08265 | 0.874555 | -1.2518 | -1.93219 | 0.0004 | 0.00614885 | yes |
| ENSG00000007174 | ENSG00000007174 | DNAH9 | chr17:11598430-11997510 | OK | 3.9245 | 1.60288 | -1.29184 | -1.40166 | 0.0004 | 0.00614885 | yes |
| ENSG00000283886 | ENSG00000283886 | BX664615.2 | chr9:39816541-40106661 | OK | 3.1721 | 1.24071 | -1.35428 | -2.06301 | 0.0004 | 0.00614885 | yes |
| ENSG00000109099 | ENSG00000109099 | PMP22 | chr17:15229776-15265707 | OK | 0.599029 | 0.103551 | -2.53228 | -0.719641 | 0.0004 | 0.00614885 | yes |
| ENSG00000110680 | ENSG00000110680 | CALCA | chr11:14904996-15082342 | OK | 1.05636 | 0.0416682 | -4.66401 | -0.708439 | 0.0004 | 0.00614885 | yes |
| ENSG00000151892 | ENSG00000151892 | GFRA1 | chr10:116056924-116273467 | OK | 11.1671 | 22.6696 | 1.0215 | 1.42627 | 0.00045 | 0.00675493 | yes |
| ENSG00000198121 | ENSG00000198121 | LPAR1 | chr9:110873262-111038458 | OK | 1.24355 | 0.521276 | -1.25435 | -1.65157 | 0.00045 | 0.00675493 | yes |
| ENSG00000282936 | ENSG00000282936 | AC004706.4 | chr17:6578147-6651634 | OK | 5.78485 | 2.38536 | -1.27807 | -2.10678 | 0.00045 | 0.00675493 | yes |
| ENSG00000198225 | ENSG00000198225 | FKBP1C | chr6:63211445-63213024 | OK | 1.76734 | 0.519612 | -1.76607 | -2.2156 | 0.00045 | 0.00675493 | yes |
| ENSG00000237945 | ENSG00000237945 | LINC00649 | chr21:33503930-33977691 | OK | 1.61033 | 0.339067 | -2.24771 | -0.407269 | 0.00045 | 0.00675493 | yes |
| ENSG00000140470 | ENSG00000140470 | ADAMTS17 | chr15:99970214-100342005 | OK | 0.207147 | 1.20211 | 2.53685 | 0.969418 | 0.0005 | 0.00734648 | yes |
| ENSG00000229644 | ENSG00000229644 | NAMPTP1 | chr10:36521720-36524234 | OK | 1.40515 | 0.648427 | -1.1157 | -1.7594 | 0.0005 | 0.00734648 | yes |
| ENSG00000172575 | ENSG00000172575 | RASGRP1 | chr15:38488102-38565575 | OK | 1.00568 | 0.324316 | -1.6327 | -0.989201 | 0.0005 | 0.00734648 | yes |
| ENSG00000149131 | ENSG00000149131 | SERPING1 | chr11:57597386-57614853 | OK | 2.53286 | 0.972302 | -1.38129 | -1.41136 | 0.00055 | 0.0079124 | yes |

| | | | | | | | | | | | | |
|---|---|---|---|---|---|---|---|---|---|---|---|---|
| ENSG00000227028 | ENSG00000227028 | SLC8A1-AS1 | chr2:39786452-40611053 | OK | 10.17 | 3.19094 | | -1.67226 | -0.558482 | 0.00055 | 0.0079124 | yes |
| ENSG00000261512 | ENSG00000261512 | AC092368.3 | chr16:46622860-46624451 | OK | 1.29633 | 0.383418 | | -1.75744 | -1.91631 | 0.00055 | 0.0079124 | yes |
| ENSG00000283199 | ENSG00000283199 | FP565324.1 | chr13:113953704-113973997 | OK | 0.780665 | 0.118273 | | -2.72258 | -1.48195 | 0.00055 | 0.0079124 | yes |
| ENSG00000224858 | ENSG00000224858 | RPL29P11 | chr3:36993331-37050918 | OK | 94.3274 | 11.2596 | | -3.06652 | -2.6112 | 0.00055 | 0.0079124 | yes |
| ENSG00000256073 | ENSG00000256073 | URB1-AS1 | chr21:32393129-32393960 | OK | 1.75778 | 4.64845 | | 1.40299 | 1.97288 | 0.0006 | 0.00848922 | yes |
| ENSG00000183831 | ENSG00000183831 | ANKRD45 | chr1:173500463-173669862 | OK | 3.36481 | 1.5526 | | -1.11584 | -1.86707 | 0.0006 | 0.00848922 | yes |
| ENSG00000227855 | ENSG00000227855 | DPY19L2P3 | chr7:29598794-29742594 | OK | 3.07799 | 1.37181 | | -1.16591 | -1.81553 | 0.0006 | 0.00848922 | yes |
| ENSG00000114013 | ENSG00000114013 | CD86 | chr3:122055365-122121139 | OK | 0.87039 | 0.246767 | | -1.81851 | -1.03895 | 0.0006 | 0.00848922 | yes |
| ENSG00000162496 | ENSG00000162496 | DHRS3 | chr1:12567909-12617731 | OK | 0.749179 | 0.117702 | | -2.67017 | -1.23948 | 0.0006 | 0.00848922 | yes |
| ENSG00000166831 | ENSG00000166831 | RBPMS2 | chr15:64739891-64775587 | OK | 1.13095 | 2.52555 | | 1.15906 | 1.94637 | 0.00065 | 0.00905328 | yes |
| ENSG00000139132 | ENSG00000139132 | FGD4 | chr12:32399528-32646050 | OK | 18.8836 | 9.13966 | | -1.04692 | -1.66675 | 0.00065 | 0.00905328 | yes |
| ENSG00000078403 | ENSG00000078403 | MLLT10 | chr10:21513477-21743630 | OK | 22.5897 | 10.3299 | | -1.12883 | -1.92573 | 0.00065 | 0.00905328 | yes |
| ENSG00000205632 | ENSG00000205632 | LINC01310 | chr22:48866769-48898386 | OK | 0.622557 | 0.214315 | | -1.53847 | -2.0332 | 0.00065 | 0.00905328 | yes |
| ENSG00000101160 | ENSG00000101160 | CTSZ | chr20:58995184-59007247 | OK | 1.14946 | 0.245129 | | -2.22934 | -1.54695 | 0.00065 | 0.00905328 | yes |
| ENSG00000125414 | ENSG00000125414 | MYH2 | chr17:10383131-10623886 | OK | 0.696698 | 0.0106431 | | -6.03254 | -0.334677 | 0.00065 | 0.00905328 | yes |
| ENSG00000125733 | ENSG00000125733 | TRIP10 | chr19:6737924-6751526 | OK | 1.96443 | 4.1031 | | 1.0626 | 1.63913 | 0.0007 | 0.00960252 | yes |

| test_id | gene_id | gene | locus | status | AHN | LRRK2 | log2(fold_change) | test_stat | p_value | q_value | significant |
|---|---|---|---|---|---|---|---|---|---|---|---|
| ENSG00000189223 | ENSG00000189223 | PAX8-AS1 | chr2:113211521-113278950 | OK | 0.029824 | 6.53101 | 7.77469 | 0.407468 | 5.00E-05 | 0.00107006 | yes |
| ENSG00000134184 | ENSG00000134184 | GSTM1 | chr1:109656080-109709551 | OK | 0.00935455 | 0.687121 | 6.19875 | 0.18465 | 5.00E-05 | 0.00107006 | yes |
| ENSG00000227617 | ENSG00000227617 | CERS6-AS1 | chr2:168455861-168913371 | OK | 0.35822 | 15.6903 | 5.45288 | 1.31838 | 5.00E-05 | 0.00107006 | yes |
| ENSG00000188219 | ENSG00000188219 | POTEE | chr2:131104846-131355769 | OK | 0.0282158 | 0.956196 | 5.08273 | 0.578042 | 5.00E-05 | 0.00107006 | yes |
| ENSG00000214548 | ENSG00000214548 | MEG3 | chr14:100779409-101027415 | OK | 0.438388 | 11.7807 | 4.74807 | 2.73428 | 5.00E-05 | 0.00107006 | yes |
| ENSG00000093010 | ENSG00000093010 | COMT | chr22:19875516-20016808 | OK | 0.919501 | 24.5693 | 4.73986 | 1.92682 | 5.00E-05 | 0.00107006 | yes |
| ENSG00000240342 | ENSG00000240342 | RPS2P5 | chr12:118149800-118372945 | OK | 8.5599 | 225.553 | 4.71973 | 6.33107 | 5.00E-05 | 0.00107006 | yes |
| ENSG00000184470 | ENSG00000184470 | TXNRD2 | chr22:19875516-20016808 | OK | 0.169473 | 4.28706 | 4.66086 | 0.851238 | 5.00E-05 | 0.00107006 | yes |
| ENSG00000188372 | ENSG00000188372 | ZP3 | chr7:76389333-76442071 | OK | 0.0998041 | 2.37661 | 4.57366 | 1.1262 | 5.00E-05 | 0.00107006 | yes |
| ENSG00000196604 | ENSG00000196604 | POTEF | chr2:130074029-130129222 | OK | 0.0909477 | 2.01349 | 4.46852 | 3.01318 | 5.00E-05 | 0.00107006 | yes |
| ENSG00000234449 | ENSG00000234449 | FAM239A | chrX:3853009-3882317 | OK | 0.20821 | 4.4807 | 4.42761 | 3.49018 | 5.00E-05 | 0.00107006 | yes |
| ENSG00000183662 | ENSG00000183662 | FAM19A1 | chr3:68004215-68545625 | OK | 1.1801 | 16.3346 | 3.79095 | 5.3978 | 5.00E-05 | 0.00107006 | yes |
| ENSG00000276805 | ENSG00000276805 | AL133216.2 | chr10:38403188-38452153 | OK | 0.0487457 | 0.636895 | 3.70771 | 0.78036 | 5.00E-05 | 0.00107006 | yes |
| ENSG00000183793 | ENSG00000183793 | NPIPA5 | chr16:15363623-15381047 | OK | 1.22716 | 14.5824 | 3.57084 | 5.21395 | 5.00E-05 | 0.00107006 | yes |
| ENSG00000109991 | ENSG00000109991 | P2RX3 | chr11:57338373-57370600 | OK | 0.158026 | 1.54745 | 3.29166 | 1.94419 | 5.00E-05 | 0.00107006 | yes |
| ENSG00000227827 | ENSG00000227827 | AC138969.2 | chr16:16356223-16377507 | OK | 0.165416 | 1.5148 | 3.19495 | 5.81043 | 5.00E-05 | 0.00107006 | yes |
| ENSG00000102195 | ENSG00000102195 | GPR50 | chrX:151175191-151181465 | OK | 0.647708 | 5.88263 | 3.18305 | 5.03135 | 5.00E-05 | 0.00107006 | yes |
| ENSG00000224557 | ENSG00000224557 | HLA-DPB2 | chr6:33112450-33129084 | OK | 0.15404 | 1.27408 | 3.04808 | 1.90933 | 5.00E-05 | 0.00107006 | yes |
| ENSG00000254681 | ENSG00000254681 | PKD1P5 | chr16:18358085-18401940 | OK | 1.62636 | 13.3512 | 3.03725 | 2.32794 | 5.00E-05 | 0.00107006 | yes |
| ENSG00000176769 | ENSG00000176769 | TCERG1L | chr10:131092390-131311721 | OK | 0.274039 | 2.17077 | 2.98576 | 3.93225 | 5.00E-05 | 0.00107006 | yes |
| ENSG00000120669 | ENSG00000120669 | SOHLH2 | chr13:36168207-36297842 | OK | 0.315417 | 2.49177 | 2.98184 | 2.41243 | 5.00E-05 | 0.00107006 | yes |
| ENSG00000186564 | ENSG00000186564 | FOXD2 | chr1:47436016-47440691 | OK | 0.107405 | 0.816405 | 2.92623 | 3.35368 | 5.00E-05 | 0.00107006 | yes |
| ENSG00000113805 | ENSG00000113805 | CNTN3 | chr3:74262567-74521140 | OK | 0.227378 | 1.61907 | 2.832 | 2.08506 | 5.00E-05 | 0.00107006 | yes |
| ENSG00000237424 | ENSG00000237424 | FOXD2-AS1 | chr1:47432132-47434641 | OK | 0.109593 | 0.765635 | 2.8045 | 2.538 | 5.00E-05 | 0.00107006 | yes |
| ENSG00000108379 | ENSG00000108379 | WNT3 | chr17:46762505-46833154 | OK | 3.36951 | 22.9078 | 2.76523 | 4.49685 | 5.00E-05 | 0.00107006 | yes |
| ENSG00000112246 | ENSG00000112246 | SIM1 | chr6:100385014-100464929 | OK | 0.121461 | 0.736313 | 2.59983 | 1.38904 | 5.00E-05 | 0.00107006 | yes |
| ENSG00000198064 | ENSG00000198064 | NPIPB13 | chr16:30222936-30254510 | OK | 0.12645 | 0.721315 | 2.51207 | 0.989315 | 5.00E-05 | 0.00107006 | yes |
| ENSG00000103449 | ENSG00000103449 | SALL1 | chr16:51135974-51151367 | OK | 0.125025 | 0.707624 | 2.50077 | 1.15377 | 5.00E-05 | 0.00107006 | yes |
| ENSG00000137310 | ENSG00000137310 | TCF19 | chr6:31158541-31180731 | OK | 0.613722 | 3.35728 | 2.45164 | 3.16378 | 5.00E-05 | 0.00107006 | yes |
| ENSG00000264204 | ENSG00000264204 | AGAP7P | chr10:46109620-46131358 | OK | 0.133915 | 0.721732 | 2.43015 | 2.12287 | 5.00E-05 | 0.00107006 | yes |
| ENSG00000135406 | ENSG00000135406 | PRPH | chr12:49292630-49331731 | OK | 0.357354 | 1.82408 | 2.35174 | 1.59816 | 5.00E-05 | 0.00107006 | yes |
| ENSG00000269993 | ENSG00000269993 | KC877982.1 | chrX:151182385-151182855 | OK | 1.32011 | 6.69061 | 2.34148 | 2.50084 | 5.00E-05 | 0.00107006 | yes |
| ENSG00000168032 | ENSG00000168032 | ENTPD3 | chr3:40313801-40453329 | OK | 0.331994 | 1.67999 | 2.33923 | 1.95911 | 5.00E-05 | 0.00107006 | yes |
| ENSG00000206503 | ENSG00000206503 | HLA-A | chr6:29941259-29945884 | OK | 6.29077 | 31.6505 | 2.33092 | 6.85024 | 5.00E-05 | 0.00107006 | yes |
| ENSG00000142173 | ENSG00000142173 | COL6A2 | chr21:46098096-46132849 | OK | 1.8385 | 9.22915 | 2.32767 | 4.41518 | 5.00E-05 | 0.00107006 | yes |
| ENSG00000146469 | ENSG00000146469 | VIP | chr6:152750797-152759765 | OK | 0.816407 | 4.05402 | 2.31199 | 3.48798 | 5.00E-05 | 0.00107006 | yes |
| ENSG00000216866 | ENSG00000216866 | RPS2P55 | chrX:40934981-40935864 | OK | 0.371244 | 1.83478 | 2.30517 | 1.94333 | 5.00E-05 | 0.00107006 | yes |
| ENSG00000093072 | ENSG00000093072 | ADA2 | chr22:17178789-17221989 | OK | 1.01645 | 4.99187 | 2.29604 | 3.834 | 5.00E-05 | 0.00107006 | yes |
| ENSG00000054803 | ENSG00000054803 | CBLN4 | chr20:55997439-56005472 | OK | 0.170019 | 0.821567 | 2.27268 | 2.55403 | 5.00E-05 | 0.00107006 | yes |
| ENSG00000102362 | ENSG00000102362 | SYTL4 | chrX:100644165-100732123 | OK | 0.393859 | 1.90242 | 2.27209 | 1.84716 | 5.00E-05 | 0.00107006 | yes |
| ENSG00000178038 | ENSG00000178038 | ALS2CL | chr3:46668996-46693704 | OK | 0.35044 | 1.64488 | 2.23074 | 2.28892 | 5.00E-05 | 0.00107006 | yes |
| ENSG00000261572 | ENSG00000261572 | AC097639.1 | chr3:32236687-32238578 | OK | 0.193095 | 0.899542 | 2.21988 | 2.19573 | 5.00E-05 | 0.00107006 | yes |
| ENSG00000280145 | ENSG00000280145 | CU638689.4 | chr21:6630181-6670695 | OK | 0.292592 | 1.34455 | 2.20016 | 1.2635 | 5.00E-05 | 0.00107006 | yes |
| ENSG00000100300 | ENSG00000100300 | TSPO | chr22:43151513-43163242 | OK | 3.53698 | 16.2026 | 2.19563 | 3.96217 | 5.00E-05 | 0.00107006 | yes |
| ENSG00000261227 | ENSG00000261227 | AC140912.1 | chr16:73232054-73233970 | OK | 1.92541 | 8.81963 | 2.19555 | 3.93938 | 5.00E-05 | 0.00107006 | yes |
| ENSG00000168743 | ENSG00000168743 | NPNT | chr4:105552619-106022478 | OK | 13.1701 | 55.5223 | 2.0758 | 5.63385 | 5.00E-05 | 0.00107006 | yes |

| | | | | | | | | | | | | |
|---|---|---|---|---|---|---|---|---|---|---|---|---|
| ENSG00000231806 | ENSG00000231806 | PCAT7 | chr9:94555068-94593793 | OK | 0.498154 | 2.03765 | | 2.03224 | 2.14186 | 5.00E-05 | 0.00107006 | yes |
| ENSG00000130208 | ENSG00000130208 | APOC1 | chr19:44905753-44919349 | OK | 3.05414 | 12.2603 | | 2.00516 | 1.89492 | 5.00E-05 | 0.00107006 | yes |
| ENSG00000151929 | ENSG00000151929 | BAG3 | chr10:119651369-119677819 | OK | 1.25417 | 5.02725 | | 2.00304 | 4.00954 | 5.00E-05 | 0.00107006 | yes |
| ENSG00000170893 | ENSG00000170893 | TRH | chr3:129974304-129977938 | OK | 3.83414 | 15.2404 | | 1.99092 | 5.13849 | 5.00E-05 | 0.00107006 | yes |
| ENSG00000233098 | ENSG00000233098 | CCDC144NL-AS1 | chr17:20814619-21043760 | OK | 4.86986 | 19.0027 | | 1.96425 | 0.700076 | 5.00E-05 | 0.00107006 | yes |
| ENSG00000134443 | ENSG00000134443 | GRP | chr18:59220167-59230774 | OK | 3.69769 | 14.3496 | | 1.95631 | 3.35945 | 5.00E-05 | 0.00107006 | yes |
| ENSG00000204381 | ENSG00000204381 | LAYN | chr11:111540279-111561745 | OK | 0.241822 | 0.937326 | | 1.95461 | 1.02446 | 5.00E-05 | 0.00107006 | yes |
| ENSG00000278200 | ENSG00000278200 | LINC01971 | chr17:81480522-81481570 | OK | 0.58324 | 2.21779 | | 1.92696 | 2.15876 | 5.00E-05 | 0.00107006 | yes |
| ENSG00000187688 | ENSG00000187688 | TRPV2 | chr17:16414523-16437003 | OK | 0.728132 | 2.74974 | | 1.91702 | 1.67131 | 5.00E-05 | 0.00107006 | yes |
| ENSG00000196843 | ENSG00000196843 | ARID5A | chr2:96536742-96552638 | OK | 0.36285 | 1.36276 | | 1.90909 | 2.02149 | 5.00E-05 | 0.00107006 | yes |
| ENSG00000142102 | ENSG00000142102 | PGGHG | chr11:289134-296107 | OK | 1.1411 | 4.243 | | 1.89466 | 2.12713 | 5.00E-05 | 0.00107006 | yes |
| ENSG00000166105 | ENSG00000166105 | GLB1L3 | chr11:134274244-134319564 | OK | 0.180882 | 0.67195 | | 1.8933 | 1.27008 | 5.00E-05 | 0.00107006 | yes |
| ENSG00000277701 | ENSG00000277701 | AC159540.2 | chr2:97281903-97291780 | OK | 0.8921 | 3.2931 | | 1.88417 | 1.87315 | 5.00E-05 | 0.00107006 | yes |
| ENSG00000175170 | ENSG00000175170 | FAM182B | chr20:25763465-25868225 | OK | 1.8368 | 6.64046 | | 1.85409 | 2.04337 | 5.00E-05 | 0.00107006 | yes |
| ENSG00000198075 | ENSG00000198075 | SULT1C4 | chr2:108377910-108388057 | OK | 1.46054 | 5.22666 | | 1.83939 | 2.8615 | 5.00E-05 | 0.00107006 | yes |
| ENSG00000138772 | ENSG00000138772 | ANXA3 | chr4:78551518-78610451 | OK | 0.587708 | 2.09403 | | 1.83311 | 1.45778 | 5.00E-05 | 0.00107006 | yes |
| ENSG00000185187 | ENSG00000185187 | SIGIRR | chr11:405715-417455 | OK | 1.24733 | 4.42687 | | 1.82744 | 1.80047 | 5.00E-05 | 0.00107006 | yes |
| ENSG00000138378 | ENSG00000138378 | STAT4 | chr2:191021525-191151596 | OK | 1.03762 | 3.59189 | | 1.79146 | 1.96769 | 5.00E-05 | 0.00107006 | yes |
| ENSG00000163932 | ENSG00000163932 | PRKCD | chr3:53156008-53192717 | OK | 3.14086 | 10.6197 | | 1.75751 | 4.51768 | 5.00E-05 | 0.00107006 | yes |
| ENSG00000065675 | ENSG00000065675 | PRKCQ | chr10:6427142-6580301 | OK | 0.181627 | 0.613065 | | 1.75506 | 1.90177 | 5.00E-05 | 0.00107006 | yes |
| ENSG00000177551 | ENSG00000177551 | NHLH2 | chr1:115836376-115843917 | OK | 2.91362 | 9.82836 | | 1.75414 | 4.20691 | 5.00E-05 | 0.00107006 | yes |
| ENSG00000089169 | ENSG00000089169 | RPH3A | chr12:112570379-112898881 | OK | 3.25478 | 10.967 | | 1.75253 | 3.15839 | 5.00E-05 | 0.00107006 | yes |
| ENSG00000064666 | ENSG00000064666 | CNN2 | chr19:1026580-1039068 | OK | 5.6224 | 18.5069 | | 1.7188 | 4.7093 | 5.00E-05 | 0.00107006 | yes |
| ENSG00000185324 | ENSG00000185324 | CDK10 | chr16:89680736-89701705 | OK | 11.0655 | 36.2544 | | 1.71208 | 2.32731 | 5.00E-05 | 0.00107006 | yes |
| ENSG00000145708 | ENSG00000145708 | CRHBP | chr5:76952712-76981158 | OK | 0.783817 | 2.56512 | | 1.71044 | 1.97194 | 5.00E-05 | 0.00107006 | yes |
| ENSG00000168952 | ENSG00000168952 | STXBP6 | chr14:24809655-25050297 | OK | 1.3362 | 4.36992 | | 1.70947 | 3.41163 | 5.00E-05 | 0.00107006 | yes |
| ENSG00000141750 | ENSG00000141750 | STAC2 | chr17:39210535-39225872 | OK | 0.752849 | 2.46194 | | 1.70936 | 3.22967 | 5.00E-05 | 0.00107006 | yes |
| ENSG00000144550 | ENSG00000144550 | CPNE9 | chr3:9703806-9729908 | OK | 0.405466 | 1.32466 | | 1.70797 | 1.46802 | 5.00E-05 | 0.00107006 | yes |
| ENSG00000128274 | ENSG00000128274 | A4GALT | chr22:42692120-42721298 | OK | 1.06261 | 3.46294 | | 1.70439 | 2.53641 | 5.00E-05 | 0.00107006 | yes |
| ENSG00000152229 | ENSG00000152229 | PSTPIP2 | chr18:45983535-46072272 | OK | 0.412172 | 1.34159 | | 1.70263 | 1.73006 | 5.00E-05 | 0.00107006 | yes |
| ENSG00000145428 | ENSG00000145428 | RNF175 | chr4:153710124-153760235 | OK | 3.89336 | 12.4208 | | 1.67367 | 2.97705 | 5.00E-05 | 0.00107006 | yes |
| ENSG00000130950 | ENSG00000130950 | NUTM2F | chr9:94318195-94328644 | OK | 0.272981 | 0.86914 | | 1.67079 | 2.02255 | 5.00E-05 | 0.00107006 | yes |
| ENSG00000159247 | ENSG00000159247 | TUBBP5 | chr9:138150074-138179774 | OK | 0.521212 | 1.64144 | | 1.65502 | 1.86381 | 5.00E-05 | 0.00107006 | yes |
| ENSG00000188488 | ENSG00000188488 | SERPINA5 | chr14:94561090-94624646 | OK | 0.3028 | 0.948812 | | 1.64776 | 0.830832 | 5.00E-05 | 0.00107006 | yes |
| ENSG00000152214 | ENSG00000152214 | RIT2 | chr18:42743226-43115691 | OK | 10.3868 | 32.4447 | | 1.64323 | 4.60262 | 5.00E-05 | 0.00107006 | yes |
| ENSG00000181195 | ENSG00000181195 | PENK | chr8:56436673-56559823 | OK | 0.941177 | 2.90475 | | 1.62588 | 1.52389 | 5.00E-05 | 0.00107006 | yes |
| ENSG00000278266 | ENSG00000278266 | AC079949.2 | chr12:127147148-127150081 | OK | 0.997401 | 3.06722 | | 1.62068 | 3.23341 | 5.00E-05 | 0.00107006 | yes |
| ENSG00000129451 | ENSG00000129451 | KLK10 | chr19:51012738-51020175 | OK | 0.698276 | 2.14654 | | 1.62015 | 1.65631 | 5.00E-05 | 0.00107006 | yes |
| ENSG00000058866 | ENSG00000058866 | DGKG | chr3:186046307-186362237 | OK | 1.95123 | 5.99407 | | 1.61915 | 1.59596 | 5.00E-05 | 0.00107006 | yes |
| ENSG00000049089 | ENSG00000049089 | COL9A2 | chr1:40300486-40317816 | OK | 1.50579 | 4.59979 | | 1.61104 | 2.34428 | 5.00E-05 | 0.00107006 | yes |
| ENSG00000101463 | ENSG00000101463 | SYNDIG1 | chr20:24469198-24666616 | OK | 0.7268 | 2.21823 | | 1.60978 | 2.44863 | 5.00E-05 | 0.00107006 | yes |
| ENSG00000104313 | ENSG00000104313 | EYA1 | chr8:71155456-71362232 | OK | 0.191016 | 0.577392 | | 1.59586 | 0.962831 | 5.00E-05 | 0.00107006 | yes |
| ENSG00000115085 | ENSG00000115085 | ZAP70 | chr2:97713559-97739862 | OK | 0.383966 | 1.14918 | | 1.58156 | 1.27258 | 5.00E-05 | 0.00107006 | yes |
| ENSG00000104888 | ENSG00000104888 | SLC17A7 | chr19:49429400-49442360 | OK | 0.493272 | 1.46411 | | 1.56957 | 2.22817 | 5.00E-05 | 0.00107006 | yes |
| ENSG00000241494 | ENSG00000241494 | AL355032.1 | chr14:101634453-101732522 | OK | 10.2978 | 30.4945 | | 1.56621 | 2.68459 | 5.00E-05 | 0.00107006 | yes |
| ENSG00000189343 | ENSG00000189343 | RPS2P46 | chr17:19417803-19492991 | OK | 17.284 | 50.3222 | | 1.54176 | 4.72321 | 5.00E-05 | 0.00107006 | yes |
| ENSG00000183036 | ENSG00000183036 | PCP4 | chr21:39867316-39929397 | OK | 12.8401 | 37.1109 | | 1.53119 | 3.72025 | 5.00E-05 | 0.00107006 | yes |

| | | | | | | | | | | | |
|---|---|---|---|---|---|---|---|---|---|---|---|
| ENSG00000258498 | ENSG00000258498 | DIO3OS | chr14:101552220-101560431 | OK | 0.610981 | 1.73162 | 1.50292 | 1.30904 | 5.00E-05 | 0.00107006 | yes |
| ENSG00000166816 | ENSG00000166816 | LDHD | chr16:75111859-75116771 | OK | 1.58827 | 4.46506 | 1.49122 | 2.41693 | 5.00E-05 | 0.00107006 | yes |
| ENSG00000147571 | ENSG00000147571 | CRH | chr8:66176381-66178725 | OK | 13.5973 | 38.1045 | 1.48664 | 5.19316 | 5.00E-05 | 0.00107006 | yes |
| ENSG00000169891 | ENSG00000169891 | REPS2 | chrX:16946690-17153280 | OK | 0.949547 | 2.6591 | 1.48562 | 1.95654 | 5.00E-05 | 0.00107006 | yes |
| ENSG00000236279 | ENSG00000236279 | CLEC2L | chr7:139523855-139544984 | OK | 3.02763 | 8.31519 | 1.45756 | 2.6999 | 5.00E-05 | 0.00107006 | yes |
| ENSG00000105509 | ENSG00000105509 | HAS1 | chr19:51713111-51723994 | OK | 0.503414 | 1.37648 | 1.45117 | 1.59465 | 5.00E-05 | 0.00107006 | yes |
| ENSG00000162595 | ENSG00000162595 | DIRAS3 | chr1:67701465-68233120 | OK | 7.29873 | 19.8722 | 1.44503 | 1.99221 | 5.00E-05 | 0.00107006 | yes |
| ENSG00000215256 | ENSG00000215256 | DHRS4-AS1 | chr14:23938730-24006408 | OK | 3.02982 | 8.18444 | 1.43365 | 2.44732 | 5.00E-05 | 0.00107006 | yes |
| ENSG00000107736 | ENSG00000107736 | CDH23 | chr10:71396933-71815947 | OK | 1.15294 | 3.09608 | 1.42513 | 2.48988 | 5.00E-05 | 0.00107006 | yes |
| ENSG00000132622 | ENSG00000132622 | HSPA12B | chr20:3732666-3753111 | OK | 0.767703 | 2.0597 | 1.42382 | 2.48179 | 5.00E-05 | 0.00107006 | yes |
| ENSG00000076706 | ENSG00000076706 | MCAM | chr11:119206275-119321521 | OK | 6.81953 | 18.271 | 1.42181 | 1.6363 | 5.00E-05 | 0.00107006 | yes |
| ENSG00000237943 | ENSG00000237943 | PRKCQ-AS1 | chr10:6580418-6616452 | OK | 0.42174 | 1.12229 | 1.41202 | 0.724417 | 5.00E-05 | 0.00107006 | yes |
| ENSG00000205664 | ENSG00000205664 | BX890604.1 | chrX:3817527-3843857 | OK | 17.6127 | 46.7974 | 1.40981 | 3.60308 | 5.00E-05 | 0.00107006 | yes |
| ENSG00000197747 | ENSG00000197747 | S100A10 | chr1:151982914-151994390 | OK | 2.14494 | 5.66338 | 1.40073 | 1.58608 | 5.00E-05 | 0.00107006 | yes |
| ENSG00000146674 | ENSG00000146674 | IGFBP3 | chr7:45912244-45921874 | OK | 0.927386 | 2.43916 | 1.39515 | 1.76998 | 5.00E-05 | 0.00107006 | yes |
| ENSG00000123496 | ENSG00000123496 | IL13RA2 | chrX:115003974-115019977 | OK | 8.71986 | 22.8092 | 1.38724 | 3.85846 | 5.00E-05 | 0.00107006 | yes |
| ENSG00000176907 | ENSG00000176907 | TCIM | chr8:40153454-40155308 | OK | 1.10437 | 2.88813 | 1.38691 | 2.37411 | 5.00E-05 | 0.00107006 | yes |
| ENSG00000186493 | ENSG00000186493 | C5orf38 | chr5:2752130-2755397 | OK | 21.6577 | 56.467 | 1.38253 | 2.53508 | 5.00E-05 | 0.00107006 | yes |
| ENSG00000227775 | ENSG00000227775 | AL031282.1 | chr1:1702729-1745992 | OK | 6.49717 | 16.8413 | 1.37412 | 1.92727 | 5.00E-05 | 0.00107006 | yes |
| ENSG00000140945 | ENSG00000140945 | CDH13 | chr16:82626802-83807834 | OK | 3.5617 | 9.22309 | 1.37268 | 1.68731 | 5.00E-05 | 0.00107006 | yes |
| ENSG00000175701 | ENSG00000175701 | LINC00116 | chr2:110211528-110285856 | OK | 24.838 | 64.1329 | 1.36851 | 3.11984 | 5.00E-05 | 0.00107006 | yes |
| ENSG00000100302 | ENSG00000100302 | RASD2 | chr22:35540867-35554001 | OK | 1.55308 | 3.99159 | 1.36183 | 3.24546 | 5.00E-05 | 0.00107006 | yes |
| ENSG00000141934 | ENSG00000141934 | PLPP2 | chr19:281039-291504 | OK | 2.99492 | 7.67281 | 1.35724 | 2.43745 | 5.00E-05 | 0.00107006 | yes |
| ENSG00000136147 | ENSG00000136147 | PHF11 | chr13:49495609-49528987 | OK | 1.44643 | 3.70334 | 1.35633 | 1.36782 | 5.00E-05 | 0.00107006 | yes |
| ENSG00000081052 | ENSG00000081052 | COL4A4 | chr2:227002710-227164113 | OK | 0.400856 | 1.02526 | 1.35483 | 2.922 | 5.00E-05 | 0.00107006 | yes |
| ENSG00000172889 | ENSG00000172889 | EGFL7 | chr9:136648609-136672678 | OK | 3.02653 | 7.73998 | 1.35466 | 2.70271 | 5.00E-05 | 0.00107006 | yes |
| ENSG00000131015 | ENSG00000131015 | ULBP2 | chr6:149941999-149949235 | OK | 1.52079 | 3.86226 | 1.34463 | 2.23561 | 5.00E-05 | 0.00107006 | yes |
| ENSG00000088882 | ENSG00000088882 | CPXM1 | chr20:2794068-2800637 | OK | 3.48864 | 8.8415 | 1.34163 | 3.6452 | 5.00E-05 | 0.00107006 | yes |
| ENSG00000281181 | ENSG00000281181 | FP236383.3 | chr21:8437628-8438551 | OK | 16.0923 | 40.5828 | 1.3345 | 3.89177 | 5.00E-05 | 0.00107006 | yes |
| ENSG00000218739 | ENSG00000218739 | CEBPZOS | chr2:37167819-37324808 | OK | 8.21951 | 20.699 | 1.33243 | 2.1576 | 5.00E-05 | 0.00107006 | yes |
| ENSG00000164326 | ENSG00000164326 | CARTPT | chr5:71719162-71721048 | OK | 813.022 | 2043.33 | 1.32955 | 6.35275 | 5.00E-05 | 0.00107006 | yes |
| ENSG00000101438 | ENSG00000101438 | SLC32A1 | chr20:38724461-38729372 | OK | 3.30077 | 8.25688 | 1.32279 | 3.48556 | 5.00E-05 | 0.00107006 | yes |
| ENSG00000187091 | ENSG00000187091 | PLCD1 | chr3:38007495-38029762 | OK | 4.87008 | 12.046 | 1.30653 | 2.52831 | 5.00E-05 | 0.00107006 | yes |
| ENSG00000163794 | ENSG00000163794 | UCN | chr2:27282391-27308445 | OK | 3.83583 | 9.4607 | 1.30241 | 2.19588 | 5.00E-05 | 0.00107006 | yes |
| ENSG00000125863 | ENSG00000125863 | MKKS | chr20:10401008-10434222 | OK | 9.07397 | 22.3266 | 1.29896 | 4.48676 | 5.00E-05 | 0.00107006 | yes |
| ENSG00000150556 | ENSG00000150556 | LYPD6B | chr2:149038106-149215262 | OK | 0.906639 | 2.22677 | 1.29635 | 1.39883 | 5.00E-05 | 0.00107006 | yes |
| ENSG00000203485 | ENSG00000203485 | INF2 | chr14:104689605-104722535 | OK | 4.02746 | 9.8234 | 1.28635 | 2.57647 | 5.00E-05 | 0.00107006 | yes |
| ENSG00000170430 | ENSG00000170430 | MGMT | chr10:129467183-129768007 | OK | 4.34802 | 10.5993 | 1.28554 | 3.26973 | 5.00E-05 | 0.00107006 | yes |
| ENSG00000088992 | ENSG00000088992 | TESC | chr12:117038922-117099479 | OK | 1.18556 | 2.87348 | 1.27723 | 1.44602 | 5.00E-05 | 0.00107006 | yes |
| ENSG00000107742 | ENSG00000107742 | SPOCK2 | chr10:72059034-72089032 | OK | 6.41859 | 15.5545 | 1.27701 | 2.35821 | 5.00E-05 | 0.00107006 | yes |
| ENSG00000198759 | ENSG00000198759 | EGFL6 | chrX:13569604-13633575 | OK | 2.88346 | 6.95928 | 1.27114 | 3.09219 | 5.00E-05 | 0.00107006 | yes |
| ENSG00000149531 | ENSG00000149531 | FRG1BP | chr20:30377371-30399257 | OK | 2.51911 | 6.05436 | 1.26506 | 1.91113 | 5.00E-05 | 0.00107006 | yes |
| ENSG00000160307 | ENSG00000160307 | S100B | chr21:46598961-46605208 | OK | 2.40601 | 5.7387 | 1.25408 | 1.98703 | 5.00E-05 | 0.00107006 | yes |
| ENSG00000256235 | ENSG00000256235 | SMIM3 | chr5:150777945-150796734 | OK | 1.1151 | 2.65593 | 1.25205 | 2.30243 | 5.00E-05 | 0.00107006 | yes |
| ENSG00000188732 | ENSG00000188732 | FAM221A | chr7:23680129-23703249 | OK | 4.68358 | 11.1029 | 1.24526 | 2.21331 | 5.00E-05 | 0.00107006 | yes |
| ENSG00000166831 | ENSG00000166831 | RBPMS2 | chr15:64739891-64775587 | OK | 1.13095 | 2.66928 | 1.23891 | 2.06208 | 5.00E-05 | 0.00107006 | yes |
| ENSG00000111666 | ENSG00000111666 | CHPT1 | chr12:101696946-101744140 | OK | 7.0751 | 16.6918 | 1.23832 | 3.03251 | 5.00E-05 | 0.00107006 | yes |

| | | | | | | | | | | | | |
|---|---|---|---|---|---|---|---|---|---|---|---|---|
| ENSG00000154027 | ENSG00000154027 | AK5 | chr1:77282050-77559969 | OK | 17.1152 | 39.96 | | 1.22328 | 3.07137 | 5.00E-05 | 0.00107006 | yes |
| ENSG00000205542 | ENSG00000205542 | TMSB4X | chrX:12975107-12977227 | OK | 2311.19 | 5333.91 | | 1.20656 | 5.2386 | 5.00E-05 | 0.00107006 | yes |
| ENSG00000253305 | ENSG00000253305 | PCDHGB6 | chr5:141330570-141512981 | OK | 20.647 | 47.5064 | | 1.20219 | 2.47362 | 5.00E-05 | 0.00107006 | yes |
| ENSG00000105355 | ENSG00000105355 | PLIN3 | chr19:4838331-4867768 | OK | 2.74343 | 6.31055 | | 1.20179 | 2.58657 | 5.00E-05 | 0.00107006 | yes |
| ENSG00000260941 | ENSG00000260941 | LINC00622 | chr1:119597701-119599271 | OK | 1.58008 | 3.62739 | | 1.19894 | 2.12825 | 5.00E-05 | 0.00107006 | yes |
| ENSG00000125968 | ENSG00000125968 | ID1 | chr20:31605282-31606515 | OK | 4.3627 | 9.99999 | | 1.1967 | 2.44577 | 5.00E-05 | 0.00107006 | yes |
| ENSG00000229119 | ENSG00000229119 | AC026403.1 | chr5:166382304-166382599 | OK | 43.203 | 97.8219 | | 1.17903 | 1.94486 | 5.00E-05 | 0.00107006 | yes |
| ENSG00000164758 | ENSG00000164758 | MED30 | chr8:117520712-117540262 | OK | 6.15211 | 13.8539 | | 1.17114 | 2.36629 | 5.00E-05 | 0.00107006 | yes |
| ENSG00000160161 | ENSG00000160161 | CILP2 | chr19:19538247-19546659 | OK | 0.683731 | 1.53353 | | 1.16535 | 1.95442 | 5.00E-05 | 0.00107006 | yes |
| ENSG00000056998 | ENSG00000056998 | GYG2 | chrX:2828787-2882820 | OK | 13.1003 | 29.2501 | | 1.15885 | 3.20673 | 5.00E-05 | 0.00107006 | yes |
| ENSG00000114023 | ENSG00000114023 | FAM162A | chr3:122384175-122416051 | OK | 6.88415 | 15.259 | | 1.14831 | 2.31906 | 5.00E-05 | 0.00107006 | yes |
| ENSG00000142178 | ENSG00000142178 | SIK1 | chr21:43414514-43427128 | OK | 1.62797 | 3.55727 | | 1.12769 | 2.75329 | 5.00E-05 | 0.00107006 | yes |
| ENSG00000121769 | ENSG00000121769 | FABP3 | chr1:31365624-31376850 | OK | 38.9411 | 84.9207 | | 1.12482 | 3.87182 | 5.00E-05 | 0.00107006 | yes |
| ENSG00000090539 | ENSG00000090539 | CHRD | chr3:184380072-184390736 | OK | 2.17088 | 4.69691 | | 1.11343 | 1.64968 | 5.00E-05 | 0.00107006 | yes |
| ENSG00000157782 | ENSG00000157782 | CABP1 | chr12:120629610-120667324 | OK | 3.21226 | 6.93931 | | 1.1112 | 2.18419 | 5.00E-05 | 0.00107006 | yes |
| ENSG00000119138 | ENSG00000119138 | KLF9 | chr9:70384596-70414624 | OK | 0.530614 | 1.14466 | | 1.10918 | 2.11345 | 5.00E-05 | 0.00107006 | yes |
| ENSG00000184160 | ENSG00000184160 | ADRA2C | chr4:3766347-3768526 | OK | 3.80479 | 8.20663 | | 1.10897 | 2.84622 | 5.00E-05 | 0.00107006 | yes |
| ENSG00000115641 | ENSG00000115641 | FHL2 | chr2:105357711-105438513 | OK | 2.45806 | 5.28775 | | 1.10513 | 2.02033 | 5.00E-05 | 0.00107006 | yes |
| ENSG00000137507 | ENSG00000137507 | LRRC32 | chr11:76657055-76670747 | OK | 0.401286 | 0.863116 | | 1.10492 | 1.2411 | 5.00E-05 | 0.00107006 | yes |
| ENSG00000237550 | ENSG00000237550 | RPL9P9 | chr15:82355141-82439153 | OK | 192.461 | 412.495 | | 1.09981 | 4.21439 | 5.00E-05 | 0.00107006 | yes |
| ENSG00000163273 | ENSG00000163273 | NPPC | chr2:231921819-231926403 | OK | 2.96095 | 6.27018 | | 1.08244 | 1.7121 | 5.00E-05 | 0.00107006 | yes |
| ENSG00000170340 | ENSG00000170340 | B3GNT2 | chr2:62196112-62224731 | OK | 2.09137 | 4.42444 | | 1.08104 | 2.5709 | 5.00E-05 | 0.00107006 | yes |
| ENSG00000260822 | ENSG00000260822 | AC004656.1 | chrX:24545515-24550466 | OK | 2.03957 | 4.31041 | | 1.07956 | 3.08629 | 5.00E-05 | 0.00107006 | yes |
| ENSG00000173715 | ENSG00000173715 | C11orf80 | chr11:66744450-66846546 | OK | 12.2271 | 25.7745 | | 1.07587 | 2.10736 | 5.00E-05 | 0.00107006 | yes |
| ENSG00000226210 | ENSG00000226210 | AC215219.1 | chr12:14521-32015 | OK | 7.0769 | 14.9006 | | 1.07419 | 3.23329 | 5.00E-05 | 0.00107006 | yes |
| ENSG00000283886 | ENSG00000283886 | BX664615.2 | chr9:39816541-40106661 | OK | 3.1721 | 6.61872 | | 1.06111 | 2.29947 | 5.00E-05 | 0.00107006 | yes |
| ENSG00000103196 | ENSG00000103196 | CRISPLD2 | chr16:84819983-84920768 | OK | 0.489297 | 1.0206 | | 1.06063 | 1.02717 | 5.00E-05 | 0.00107006 | yes |
| ENSG00000130707 | ENSG00000130707 | ASS1 | chr9:130444928-130501274 | OK | 19.3717 | 40.3887 | | 1.06001 | 3.46266 | 5.00E-05 | 0.00107006 | yes |
| ENSG00000265972 | ENSG00000265972 | TXNIP | chr1:145992434-145996600 | OK | 4.94491 | 10.3035 | | 1.05913 | 2.58944 | 5.00E-05 | 0.00107006 | yes |
| ENSG00000279249 | ENSG00000279249 | AC007614.5 | chr16:49341662-49350554 | OK | 0.642771 | 1.33906 | | 1.05884 | 2.549 | 5.00E-05 | 0.00107006 | yes |
| ENSG00000260244 | ENSG00000260244 | AC104083.1 | chr4:155734447-155737062 | OK | 2.0681 | 4.29551 | | 1.05453 | 2.45579 | 5.00E-05 | 0.00107006 | yes |
| ENSG00000113504 | ENSG00000113504 | SLC12A7 | chr5:1050375-1112035 | OK | 3.68146 | 7.64048 | | 1.05338 | 2.13872 | 5.00E-05 | 0.00107006 | yes |
| ENSG00000184831 | ENSG00000184831 | APOO | chrX:23833352-23939507 | OK | 21.2125 | 43.9695 | | 1.05159 | 2.94969 | 5.00E-05 | 0.00107006 | yes |
| ENSG00000176485 | ENSG00000176485 | PLA2G16 | chr11:63573194-63621671 | OK | 3.54083 | 7.33411 | | 1.05053 | 1.94088 | 5.00E-05 | 0.00107006 | yes |
| ENSG00000163820 | ENSG00000163820 | FYCO1 | chr3:45917898-45995824 | OK | 0.490879 | 1.01599 | | 1.04945 | 1.4902 | 5.00E-05 | 0.00107006 | yes |
| ENSG00000107551 | ENSG00000107551 | RASSF4 | chr10:44811023-44995891 | OK | 4.56241 | 9.44158 | | 1.04923 | 1.94635 | 5.00E-05 | 0.00107006 | yes |
| ENSG00000178821 | ENSG00000178821 | TMEM52 | chr1:1917589-1919273 | OK | 4.50109 | 9.31262 | | 1.04891 | 1.92256 | 5.00E-05 | 0.00107006 | yes |
| ENSG00000214485 | ENSG00000214485 | RPL7P1 | chr5:150053290-150113372 | OK | 33.0486 | 68.2459 | | 1.04615 | 3.33382 | 5.00E-05 | 0.00107006 | yes |
| ENSG00000180440 | ENSG00000180440 | SERTM1 | chr13:36673911-36697839 | OK | 4.68164 | 9.66535 | | 1.04581 | 3.36183 | 5.00E-05 | 0.00107006 | yes |
| ENSG00000182718 | ENSG00000182718 | ANXA2 | chr15:60347133-60402883 | OK | 7.82341 | 16.1488 | | 1.04555 | 2.45909 | 5.00E-05 | 0.00107006 | yes |
| ENSG00000156110 | ENSG00000156110 | ADK | chr10:74151184-74709303 | OK | 14.0495 | 28.9376 | | 1.04243 | 3.27176 | 5.00E-05 | 0.00107006 | yes |
| ENSG00000117971 | ENSG00000117971 | CHRNB4 | chr15:78624118-78735495 | OK | 2.005 | 4.12275 | | 1.04 | 2.12848 | 5.00E-05 | 0.00107006 | yes |
| ENSG00000260769 | ENSG00000260769 | AC007614.4 | chr16:49336682-49338993 | OK | 1.81695 | 3.73332 | | 1.03894 | 2.0226 | 5.00E-05 | 0.00107006 | yes |
| ENSG00000233670 | ENSG00000233670 | PIRT | chr17:10822474-10838445 | OK | 1.06714 | 2.19212 | | 1.03858 | 2.22899 | 5.00E-05 | 0.00107006 | yes |
| ENSG00000113645 | ENSG00000113645 | WWC1 | chr5:168291650-168472303 | OK | 4.56007 | 9.34625 | | 1.03533 | 1.73309 | 5.00E-05 | 0.00107006 | yes |
| ENSG00000101282 | ENSG00000101282 | RSPO4 | chr20:958451-1002264 | OK | 1.42419 | 2.91384 | | 1.03278 | 1.76156 | 5.00E-05 | 0.00107006 | yes |
| ENSG00000174951 | ENSG00000174951 | FUT1 | chr19:48748010-48755390 | OK | 0.693226 | 1.40889 | | 1.02316 | 1.38955 | 5.00E-05 | 0.00107006 | yes |

| | | | | | | | | | | | |
|---|---|---|---|---|---|---|---|---|---|---|---|
| ENSG00000167711 | ENSG00000167711 | SERPINF2 | chr17:1742835-1755268 | OK | 0.891831 | 1.81144 | 1.0223 | 1.41785 | 5.00E-05 | 0.00107006 | yes |
| ENSG00000170293 | ENSG00000170293 | CMTM8 | chr3:32238678-32370325 | OK | 1.97394 | 3.99727 | 1.01794 | 1.65975 | 5.00E-05 | 0.00107006 | yes |
| ENSG00000197355 | ENSG00000197355 | UAP1L1 | chr9:137077500-137084539 | OK | 1.96763 | 3.98317 | 1.01746 | 2.10529 | 5.00E-05 | 0.00107006 | yes |
| ENSG00000135063 | ENSG00000135063 | FAM189A2 | chr9:69324571-69392455 | OK | 2.46648 | 4.98123 | 1.01405 | 2.00603 | 5.00E-05 | 0.00107006 | yes |
| ENSG00000164434 | ENSG00000164434 | FABP7 | chr6:122779474-122784074 | OK | 4.9196 | 9.93474 | 1.01394 | 2.15492 | 5.00E-05 | 0.00107006 | yes |
| ENSG00000186897 | ENSG00000186897 | C1QL4 | chr12:49332410-49337188 | OK | 3.88448 | 7.80364 | 1.00642 | 2.4853 | 5.00E-05 | 0.00107006 | yes |
| ENSG00000117691 | ENSG00000117691 | NENF | chr1:212432886-212446379 | OK | 17.9537 | 35.9778 | 1.00282 | 2.98979 | 5.00E-05 | 0.00107006 | yes |
| ENSG00000167333 | ENSG00000167333 | TRIM68 | chr11:4598671-4608259 | OK | 3.2239 | 6.44923 | 1.00032 | 2.29078 | 5.00E-05 | 0.00107006 | yes |
| ENSG00000126070 | ENSG00000126070 | AGO3 | chr1:35930717-36072500 | OK | 6.4875 | 3.2325 | -1.00501 | -1.54719 | 5.00E-05 | 0.00107006 | yes |
| ENSG00000157240 | ENSG00000157240 | FZD1 | chr7:91264363-91271326 | OK | 11.8699 | 5.90835 | -1.00648 | -4.17119 | 5.00E-05 | 0.00107006 | yes |
| ENSG00000169515 | ENSG00000169515 | CCDC8 | chr19:46410371-46413584 | OK | 1.17686 | 0.579843 | -1.02121 | -1.68311 | 5.00E-05 | 0.00107006 | yes |
| ENSG00000182674 | ENSG00000182674 | KCNB2 | chr8:72537390-72938349 | OK | 3.61009 | 1.77857 | -1.02131 | -2.4892 | 5.00E-05 | 0.00107006 | yes |
| ENSG00000174501 | ENSG00000174501 | ANKRD36C | chr2:95836918-95991831 | OK | 19.4764 | 9.574 | -1.02453 | -2.24463 | 5.00E-05 | 0.00107006 | yes |
| ENSG00000160145 | ENSG00000160145 | KALRN | chr3:124080022-124726325 | OK | 40.3664 | 19.8352 | -1.02509 | -1.80955 | 5.00E-05 | 0.00107006 | yes |
| ENSG00000076928 | ENSG00000076928 | ARHGEF1 | chr19:41883160-41930150 | OK | 29.4336 | 14.4626 | -1.02513 | -2.39203 | 5.00E-05 | 0.00107006 | yes |
| ENSG00000159263 | ENSG00000159263 | SIM2 | chr21:36698772-36749917 | OK | 2.32602 | 1.13946 | -1.02951 | -1.9607 | 5.00E-05 | 0.00107006 | yes |
| ENSG00000144460 | ENSG00000144460 | NYAP2 | chr2:225399709-225654018 | OK | 2.58098 | 1.25946 | -1.03511 | -2.62346 | 5.00E-05 | 0.00107006 | yes |
| ENSG00000080854 | ENSG00000080854 | IGSF9B | chr11:133908563-133956985 | OK | 18.6022 | 9.03637 | -1.04166 | -1.52527 | 5.00E-05 | 0.00107006 | yes |
| ENSG00000186395 | ENSG00000186395 | KRT10 | chr17:40818116-40836274 | OK | 24.8917 | 12.0808 | -1.04295 | -1.91711 | 5.00E-05 | 0.00107006 | yes |
| ENSG00000143614 | ENSG00000143614 | GATAD2B | chr1:153789029-153946696 | OK | 23.0683 | 11.1845 | -1.04442 | -1.84272 | 5.00E-05 | 0.00107006 | yes |
| ENSG00000273079 | ENSG00000273079 | GRIN2B | chr12:13437941-13981957 | OK | 14.409 | 6.97191 | -1.04735 | -1.6064 | 5.00E-05 | 0.00107006 | yes |
| ENSG00000137770 | ENSG00000137770 | CTDSPL2 | chr15:44427233-44536923 | OK | 14.5224 | 6.99239 | -1.05443 | -1.61636 | 5.00E-05 | 0.00107006 | yes |
| ENSG00000223756 | ENSG00000223756 | TSSC2 | chr11:3380960-3408978 | OK | 8.07841 | 3.8632 | -1.06428 | -1.77957 | 5.00E-05 | 0.00107006 | yes |
| ENSG00000146707 | ENSG00000146707 | POMZP3 | chr7:76549359-76627982 | OK | 14.6694 | 6.99893 | -1.0676 | -2.37285 | 5.00E-05 | 0.00107006 | yes |
| ENSG00000187323 | ENSG00000187323 | DCC | chr18:52340171-53568283 | OK | 27.1099 | 12.9323 | -1.06784 | -2.20516 | 5.00E-05 | 0.00107006 | yes |
| ENSG00000188738 | ENSG00000188738 | FSIP2 | chr2:185719873-185833290 | OK | 3.90559 | 1.85583 | -1.07347 | -1.26894 | 5.00E-05 | 0.00107006 | yes |
| ENSG00000153094 | ENSG00000153094 | BCL2L11 | chr2:111119377-111168447 | OK | 1.93353 | 0.918485 | -1.07391 | -0.960922 | 5.00E-05 | 0.00107006 | yes |
| ENSG00000083168 | ENSG00000083168 | KAT6A | chr8:41929478-42051990 | OK | 21.3847 | 10.1515 | -1.07489 | -2.10717 | 5.00E-05 | 0.00107006 | yes |
| ENSG00000221843 | ENSG00000221843 | C2orf16 | chr2:27537385-27582721 | OK | 1.81771 | 0.861685 | -1.07689 | -3.28087 | 5.00E-05 | 0.00107006 | yes |
| ENSG00000170381 | ENSG00000170381 | SEMA3E | chr7:83363905-83649010 | OK | 2.6503 | 1.255 | -1.07847 | -1.54974 | 5.00E-05 | 0.00107006 | yes |
| ENSG00000051523 | ENSG00000051523 | CYBA | chr16:88643282-88651152 | OK | 19.3224 | 9.14697 | -1.07891 | -2.52328 | 5.00E-05 | 0.00107006 | yes |
| ENSG00000069020 | ENSG00000069020 | MAST4 | chr5:66596360-67169595 | OK | 7.66831 | 3.58412 | -1.09729 | -1.7813 | 5.00E-05 | 0.00107006 | yes |
| ENSG00000247765 | ENSG00000247765 | AC068446.2 | chr15:21298232-21325241 | OK | 3.17538 | 1.48027 | -1.10107 | -2.15963 | 5.00E-05 | 0.00107006 | yes |
| ENSG00000185129 | ENSG00000185129 | PURA | chr5:140072856-140125619 | OK | 2.53048 | 1.17774 | -1.10339 | -1.44813 | 5.00E-05 | 0.00107006 | yes |
| ENSG00000250486 | ENSG00000250486 | FAM218A | chr4:164954445-164977668 | OK | 5.60879 | 2.60613 | -1.10578 | -2.55518 | 5.00E-05 | 0.00107006 | yes |
| ENSG00000156052 | ENSG00000156052 | GNAQ | chr9:77716086-78031458 | OK | 83.4128 | 38.7437 | -1.10631 | -2.64278 | 5.00E-05 | 0.00107006 | yes |
| ENSG00000154975 | ENSG00000154975 | CA10 | chr17:51630312-52160017 | OK | 17.56 | 8.13823 | -1.10951 | -2.74476 | 5.00E-05 | 0.00107006 | yes |
| ENSG00000080603 | ENSG00000080603 | SRCAP | chr16:30697706-30776307 | OK | 22.293 | 10.3264 | -1.11026 | -1.4807 | 5.00E-05 | 0.00107006 | yes |
| ENSG00000156253 | ENSG00000156253 | RWDD2B | chr21:29004383-29019378 | OK | 9.05073 | 4.18499 | -1.11281 | -2.54979 | 5.00E-05 | 0.00107006 | yes |
| ENSG00000070018 | ENSG00000070018 | LRP6 | chr12:12049843-12267012 | OK | 13.8295 | 6.3845 | -1.1151 | -1.58121 | 5.00E-05 | 0.00107006 | yes |
| ENSG00000181585 | ENSG00000181585 | TMIE | chr3:46701332-46710886 | OK | 2.30976 | 1.06545 | -1.11628 | -1.82265 | 5.00E-05 | 0.00107006 | yes |
| ENSG00000103226 | ENSG00000103226 | NOMO3 | chr16:16232494-16294814 | OK | 15.2695 | 7.04345 | -1.1163 | -1.45799 | 5.00E-05 | 0.00107006 | yes |
| ENSG00000170122 | ENSG00000170122 | FOXD4 | chr9:116230-118204 | OK | 2.00952 | 0.926142 | -1.11754 | -1.80684 | 5.00E-05 | 0.00107006 | yes |
| ENSG00000113140 | ENSG00000113140 | SPARC | chr5:151661095-151724782 | OK | 14.1543 | 6.51571 | -1.11925 | -2.03373 | 5.00E-05 | 0.00107006 | yes |
| ENSG00000185668 | ENSG00000185668 | POU3F1 | chr1:38044610-38046794 | OK | 8.13667 | 3.73891 | -1.12182 | -2.89846 | 5.00E-05 | 0.00107006 | yes |
| ENSG00000183474 | ENSG00000183474 | GTF2H2C | chr5:69560207-69594723 | OK | 15.6448 | 7.15413 | -1.12883 | -2.13055 | 5.00E-05 | 0.00107006 | yes |
| ENSG00000067182 | ENSG00000067182 | TNFRSF1A | chr12:6328756-6342114 | OK | 2.0249 | 0.921176 | -1.1363 | -1.01516 | 5.00E-05 | 0.00107006 | yes |

| | | | | | | | | | | | |
|---|---|---|---|---|---|---|---|---|---|---|---|
| ENSG00000148737 | ENSG00000148737 | TCF7L2 | chr10:112950249-113167678 | OK | 2.05227 | 0.932544 | -1.13798 | -0.854357 | 5.00E-05 | 0.00107006 | yes |
| ENSG00000112319 | ENSG00000112319 | EYA4 | chr6:133240597-133895553 | OK | 1.08971 | 0.495143 | -1.13803 | -1.22472 | 5.00E-05 | 0.00107006 | yes |
| ENSG00000132688 | ENSG00000132688 | NES | chr1:156668762-156677397 | OK | 12.5229 | 5.68279 | -1.13989 | -4.46354 | 5.00E-05 | 0.00107006 | yes |
| ENSG00000071537 | ENSG00000071537 | SEL1L | chr14:81471548-81533861 | OK | 34.8574 | 15.7501 | -1.1461 | -2.27116 | 5.00E-05 | 0.00107006 | yes |
| ENSG00000144747 | ENSG00000144747 | TMF1 | chr3:68975213-69080408 | OK | 11.3396 | 5.12019 | -1.14709 | -2.74737 | 5.00E-05 | 0.00107006 | yes |
| ENSG00000120594 | ENSG00000120594 | PLXDC2 | chr10:19816238-20289856 | OK | 14.1772 | 6.39832 | -1.14781 | -3.99578 | 5.00E-05 | 0.00107006 | yes |
| ENSG00000130338 | ENSG00000130338 | TULP4 | chr6:158232235-158511828 | OK | 52.8138 | 23.8181 | -1.14886 | -1.50823 | 5.00E-05 | 0.00107006 | yes |
| ENSG00000107731 | ENSG00000107731 | UNC5B | chr10:71212569-71302864 | OK | 3.57957 | 1.59969 | -1.16199 | -2.94793 | 5.00E-05 | 0.00107006 | yes |
| ENSG00000203666 | ENSG00000203666 | EFCAB2 | chr1:244969349-245127164 | OK | 13.335 | 5.95197 | -1.16378 | -1.68983 | 5.00E-05 | 0.00107006 | yes |
| ENSG00000161835 | ENSG00000161835 | GRASP | chr12:52006939-52015889 | OK | 4.32249 | 1.92142 | -1.16969 | -1.63136 | 5.00E-05 | 0.00107006 | yes |
| ENSG00000182752 | ENSG00000182752 | PAPPA | chr9:116153803-116402322 | OK | 1.11285 | 0.493055 | -1.17444 | -1.35057 | 5.00E-05 | 0.00107006 | yes |
| ENSG00000026508 | ENSG00000026508 | CD44 | chr11:35138869-35232402 | OK | 0.657309 | 0.290595 | -1.17756 | -0.360647 | 5.00E-05 | 0.00107006 | yes |
| ENSG00000090857 | ENSG00000090857 | PDPR | chr16:70113625-70187361 | OK | 9.62574 | 4.25452 | -1.1779 | -3.74019 | 5.00E-05 | 0.00107006 | yes |
| ENSG00000119772 | ENSG00000119772 | DNMT3A | chr2:25227854-25342590 | OK | 38.8005 | 17.0695 | -1.18465 | -2.69007 | 5.00E-05 | 0.00107006 | yes |
| ENSG00000139132 | ENSG00000139132 | FGD4 | chr12:32399528-32646050 | OK | 18.8836 | 8.30396 | -1.18526 | -1.65945 | 5.00E-05 | 0.00107006 | yes |
| ENSG00000267651 | ENSG00000267651 | AC015961.1 | chr18:37234118-37236242 | OK | 3.84234 | 1.678 | -1.19524 | -2.42004 | 5.00E-05 | 0.00107006 | yes |
| ENSG00000204524 | ENSG00000204524 | ZNF805 | chr19:57240684-57255135 | OK | 2.78522 | 1.21345 | -1.19868 | -2.14754 | 5.00E-05 | 0.00107006 | yes |
| ENSG00000274602 | ENSG00000274602 | PI4KAP1 | chr22:18533645-18577968 | OK | 23.1646 | 9.99679 | -1.21238 | -3.8609 | 5.00E-05 | 0.00107006 | yes |
| ENSG00000151229 | ENSG00000151229 | SLC2A13 | chr12:39626166-40106089 | OK | 20.0032 | 8.51932 | -1.23142 | -4.27323 | 5.00E-05 | 0.00107006 | yes |
| ENSG00000164093 | ENSG00000164093 | PITX2 | chr4:110617422-110642123 | OK | 33.7479 | 14.3417 | -1.23458 | -3.82574 | 5.00E-05 | 0.00107006 | yes |
| ENSG00000203875 | ENSG00000203875 | SNHG5 | chr6:85660949-85678736 | OK | 56.0587 | 23.6967 | -1.24225 | -2.81332 | 5.00E-05 | 0.00107006 | yes |
| ENSG00000242086 | ENSG00000242086 | LINC00969 | chr3:195658061-195741123 | OK | 40.7679 | 17.1671 | -1.24778 | -2.74416 | 5.00E-05 | 0.00107006 | yes |
| ENSG00000165966 | ENSG00000165966 | PDZRN4 | chr12:41188447-41574590 | OK | 15.6775 | 6.46726 | -1.27747 | -4.40186 | 5.00E-05 | 0.00107006 | yes |
| ENSG00000166987 | ENSG00000166987 | MBD6 | chr12:57520709-57547331 | OK | 25.5798 | 10.5454 | -1.27839 | -1.27743 | 5.00E-05 | 0.00107006 | yes |
| ENSG00000227888 | ENSG00000227888 | FAM66A | chr8:12362018-12388296 | OK | 19.0138 | 7.74229 | -1.29621 | -2.80265 | 5.00E-05 | 0.00107006 | yes |
| ENSG00000154620 | ENSG00000154620 | TMSB4Y | chrY:13703566-13706024 | OK | 6.85263 | 2.7763 | -1.30349 | -2.81117 | 5.00E-05 | 0.00107006 | yes |
| ENSG00000146858 | ENSG00000146858 | ZC3HAV1L | chr7:139025705-139036029 | OK | 2.05597 | 0.826264 | -1.31515 | -1.96115 | 5.00E-05 | 0.00107006 | yes |
| ENSG00000118407 | ENSG00000118407 | FILIP1 | chr6:75285013-75493738 | OK | 1.24077 | 0.497546 | -1.31833 | -2.09265 | 5.00E-05 | 0.00107006 | yes |
| ENSG00000177426 | ENSG00000177426 | TGIF1 | chr18:3411607-3459978 | OK | 1.33287 | 0.527366 | -1.33766 | -0.812883 | 5.00E-05 | 0.00107006 | yes |
| ENSG00000125675 | ENSG00000125675 | GRIA3 | chrX:123184152-123490915 | OK | 4.95274 | 1.9589 | -1.33818 | -1.98799 | 5.00E-05 | 0.00107006 | yes |
| ENSG00000186517 | ENSG00000186517 | ARHGAP30 | chr1:161046945-161069970 | OK | 0.691595 | 0.272021 | -1.34621 | -1.24561 | 5.00E-05 | 0.00107006 | yes |
| ENSG00000223855 | ENSG00000223855 | HRAT92 | chr7:520390-525232 | OK | 0.807789 | 0.315441 | -1.35661 | -1.92086 | 5.00E-05 | 0.00107006 | yes |
| ENSG00000239268 | ENSG00000239268 | AC092691.1 | chr3:117672153-117997592 | OK | 105.179 | 41.0415 | -1.35768 | -4.13721 | 5.00E-05 | 0.00107006 | yes |
| ENSG00000214425 | ENSG00000214425 | LRRC37A4P | chr17:45506740-45563230 | OK | 2.95142 | 1.13388 | -1.38014 | -2.46801 | 5.00E-05 | 0.00107006 | yes |
| ENSG00000185900 | ENSG00000185900 | POMK | chr8:43093505-43123434 | OK | 25.103 | 9.56724 | -1.39168 | -4.41475 | 5.00E-05 | 0.00107006 | yes |
| ENSG00000274422 | ENSG00000274422 | AC245060.5 | chr22:22283927-22287220 | OK | 1.96689 | 0.749301 | -1.3923 | -2.55188 | 5.00E-05 | 0.00107006 | yes |
| ENSG00000181215 | ENSG00000181215 | C4orf50 | chr4:5897372-6018507 | OK | 1.95024 | 0.742949 | -1.39232 | -3.71321 | 5.00E-05 | 0.00107006 | yes |
| ENSG00000261186 | ENSG00000261186 | LINC01238 | chr2:242087350-242088457 | OK | 3.97193 | 1.50015 | -1.40473 | -2.11516 | 5.00E-05 | 0.00107006 | yes |
| ENSG00000148143 | ENSG00000148143 | ZNF462 | chr9:106863096-107102988 | OK | 18.3992 | 6.88455 | -1.41821 | -2.11943 | 5.00E-05 | 0.00107006 | yes |
| ENSG00000178394 | ENSG00000178394 | HTR1A | chr5:63957892-63981043 | OK | 1.7432 | 0.645788 | -1.4326 | -1.55175 | 5.00E-05 | 0.00107006 | yes |
| ENSG00000130518 | ENSG00000130518 | KIAA1683 | chr19:18257096-18274509 | OK | 2.89909 | 1.06066 | -1.45064 | -2.73219 | 5.00E-05 | 0.00107006 | yes |
| ENSG00000257800 | ENSG00000257800 | FNBP1P1 | chr2:74120679-74123218 | OK | 3.3079 | 1.20749 | -1.4539 | -2.64057 | 5.00E-05 | 0.00107006 | yes |
| ENSG00000229178 | ENSG00000229178 | AC233280.1 | chr3:195655564-195657927 | OK | 1.54299 | 0.555711 | -1.47332 | -1.85429 | 5.00E-05 | 0.00107006 | yes |
| ENSG00000106018 | ENSG00000106018 | VIPR2 | chr7:159006521-159144957 | OK | 1.87866 | 0.67513 | -1.47646 | -2.28227 | 5.00E-05 | 0.00107006 | yes |
| ENSG00000235244 | ENSG00000235244 | DANT2 | chrX:115917270-115969089 | OK | 1.47212 | 0.527627 | -1.4803 | -2.07477 | 5.00E-05 | 0.00107006 | yes |
| ENSG00000226759 | ENSG00000226759 | DAB1-AS1 | chr1:56929209-58546802 | OK | 1.09686 | 0.392246 | -1.48355 | -0.432672 | 5.00E-05 | 0.00107006 | yes |
| ENSG00000215458 | ENSG00000215458 | AATBC | chr21:43805757-43812567 | OK | 1.74986 | 0.613396 | -1.51235 | -2.26556 | 5.00E-05 | 0.00107006 | yes |

| | | | | | | | | | | | |
|---|---|---|---|---|---|---|---|---|---|---|---|
| ENSG00000248213 | ENSG00000248213 | CICP16 | chr4:118635969-118638782 | OK | 1.38518 | 0.480703 | -1.52686 | -2.3025 | 5.00E-05 | 0.00107006 | yes |
| ENSG00000227028 | ENSG00000227028 | SLC8A1-AS1 | chr2:39786452-40611053 | OK | 10.17 | 3.50961 | -1.53493 | -0.749295 | 5.00E-05 | 0.00107006 | yes |
| ENSG00000180422 | ENSG00000180422 | LINC00304 | chr16:89159145-89164245 | OK | 2.64424 | 0.895567 | -1.56198 | -2.22881 | 5.00E-05 | 0.00107006 | yes |
| ENSG00000230606 | ENSG00000230606 | AC092683.1 | chr2:97416164-97434847 | OK | 15.0651 | 5.07512 | -1.5697 | -3.02318 | 5.00E-05 | 0.00107006 | yes |
| ENSG00000204525 | ENSG00000204525 | HLA-C | chr6:31268748-31357637 | OK | 13.7539 | 4.60436 | -1.57877 | -2.93917 | 5.00E-05 | 0.00107006 | yes |
| ENSG00000198569 | ENSG00000198569 | SLC34A3 | chr9:137230756-137236554 | OK | 0.879447 | 0.294005 | -1.58075 | -1.78774 | 5.00E-05 | 0.00107006 | yes |
| ENSG00000010278 | ENSG00000010278 | CD9 | chr12:6199714-6238271 | OK | 0.893273 | 0.289799 | -1.62405 | -0.889911 | 5.00E-05 | 0.00107006 | yes |
| ENSG00000233013 | ENSG00000233013 | FAM157B | chr9:138217067-138253217 | OK | 2.24947 | 0.725912 | -1.63172 | -1.24552 | 5.00E-05 | 0.00107006 | yes |
| ENSG00000049759 | ENSG00000049759 | NEDD4L | chr18:58044366-58401540 | OK | 191.496 | 60.4037 | -1.6646 | -1.31208 | 5.00E-05 | 0.00107006 | yes |
| ENSG00000159674 | ENSG00000159674 | SPON2 | chr4:1166931-1208962 | OK | 3.39165 | 1.04793 | -1.69444 | -1.59809 | 5.00E-05 | 0.00107006 | yes |
| ENSG00000198225 | ENSG00000198225 | FKBP1C | chr6:63211445-63213024 | OK | 1.76734 | 0.544997 | -1.69726 | -2.1471 | 5.00E-05 | 0.00107006 | yes |
| ENSG00000154529 | ENSG00000154529 | CNTNAP3B | chr9:41890313-42129510 | OK | 3.03891 | 0.931638 | -1.70571 | -2.07931 | 5.00E-05 | 0.00107006 | yes |
| ENSG00000282458 | ENSG00000282458 | WASH5P | chr19:60950-71626 | OK | 3.91996 | 1.18191 | -1.72972 | -1.34851 | 5.00E-05 | 0.00107006 | yes |
| ENSG00000163219 | ENSG00000163219 | ARHGAP25 | chr2:68679600-68826833 | OK | 1.16437 | 0.34983 | -1.73483 | -1.17467 | 5.00E-05 | 0.00107006 | yes |
| ENSG00000268350 | ENSG00000268350 | FAM156A | chrX:52926401-52995472 | OK | 3.61638 | 1.08281 | -1.73976 | -1.65338 | 5.00E-05 | 0.00107006 | yes |
| ENSG00000164031 | ENSG00000164031 | DNAJB14 | chr4:99896247-99946726 | OK | 30.0078 | 8.82264 | -1.76606 | -1.77117 | 5.00E-05 | 0.00107006 | yes |
| ENSG00000088827 | ENSG00000088827 | SIGLEC1 | chr20:3686969-3707128 | OK | 1.41649 | 0.410691 | -1.7862 | -3.20863 | 5.00E-05 | 0.00107006 | yes |
| ENSG00000180938 | ENSG00000180938 | ZNF572 | chr8:124973297-124979389 | OK | 0.739201 | 0.212486 | -1.7986 | -2.20788 | 5.00E-05 | 0.00107006 | yes |
| ENSG00000224769 | ENSG00000224769 | MUC20P1 | chr3:195614946-195620233 | OK | 1.87597 | 0.537149 | -1.80425 | -2.38928 | 5.00E-05 | 0.00107006 | yes |
| ENSG00000080224 | ENSG00000080224 | EPHA6 | chr3:96814580-97752460 | OK | 5.25375 | 1.47812 | -1.82959 | -2.79052 | 5.00E-05 | 0.00107006 | yes |
| ENSG00000259098 | ENSG00000259098 | AC025884.2 | chr15:22258137-22258848 | OK | 14.4225 | 4.05391 | -1.83093 | -3.62236 | 5.00E-05 | 0.00107006 | yes |
| ENSG00000186867 | ENSG00000186867 | QRFPR | chr4:121329311-121381059 | OK | 2.57302 | 0.717161 | -1.84309 | -2.5819 | 5.00E-05 | 0.00107006 | yes |
| ENSG00000140937 | ENSG00000140937 | CDH11 | chr16:64943752-65126112 | OK | 13.8899 | 3.82987 | -1.85867 | -2.69196 | 5.00E-05 | 0.00107006 | yes |
| ENSG00000181722 | ENSG00000181722 | ZBTB20 | chr3:114314500-115147271 | OK | 4.24206 | 1.1679 | -1.86085 | -1.29899 | 5.00E-05 | 0.00107006 | yes |
| ENSG00000215156 | ENSG00000215156 | AC138409.1 | chr5:34164697-34244796 | OK | 2.76927 | 0.761196 | -1.86317 | -3.10014 | 5.00E-05 | 0.00107006 | yes |
| ENSG00000274012 | ENSG00000274012 | RN7SL2 | chr14:49861175-49864379 | OK | 15342.4 | 4196.38 | -1.87031 | -7.94678 | 5.00E-05 | 0.00107006 | yes |
| ENSG00000164692 | ENSG00000164692 | COL1A2 | chr7:94394560-94431232 | OK | 2.74769 | 0.6976 | -1.97775 | -1.97548 | 5.00E-05 | 0.00107006 | yes |
| ENSG00000185760 | ENSG00000185760 | KCNQ5 | chr6:72621791-73198851 | OK | 0.675678 | 0.171396 | -1.979 | -0.768082 | 5.00E-05 | 0.00107006 | yes |
| ENSG00000260409 | ENSG00000260409 | AC012414.5 | chr15:20729746-20756183 | OK | 5.52346 | 1.38096 | -1.9999 | -3.61271 | 5.00E-05 | 0.00107006 | yes |
| ENSG00000149256 | ENSG00000149256 | TENM4 | chr11:78652830-79440948 | OK | 125.725 | 31.3669 | -2.00296 | -11.3947 | 5.00E-05 | 0.00107006 | yes |
| ENSG00000122012 | ENSG00000122012 | SV2C | chr5:76081945-76353939 | OK | 5.72331 | 1.42058 | -2.01037 | -5.58462 | 5.00E-05 | 0.00107006 | yes |
| ENSG00000232599 | ENSG00000232599 | AL008707.1 | chrX:125203804-125204338 | OK | 46.8613 | 11.5024 | -2.02646 | -4.28655 | 5.00E-05 | 0.00107006 | yes |
| ENSG00000159450 | ENSG00000159450 | TCHH | chr1:152106316-152115454 | OK | 0.815336 | 0.195474 | -2.06042 | -3.1736 | 5.00E-05 | 0.00107006 | yes |
| ENSG00000180229 | ENSG00000180229 | HERC2P3 | chr15:20379494-20506180 | OK | 21.7597 | 5.19222 | -2.06724 | -3.35007 | 5.00E-05 | 0.00107006 | yes |
| ENSG00000157227 | ENSG00000157227 | MMP14 | chr14:22836556-22849027 | OK | 1.56984 | 0.372561 | -2.07507 | -1.45847 | 5.00E-05 | 0.00107006 | yes |
| ENSG00000184956 | ENSG00000184956 | MUC6 | chr11:1012820-1036706 | OK | 1.57235 | 0.37304 | -2.07552 | -1.70757 | 5.00E-05 | 0.00107006 | yes |
| ENSG00000164778 | ENSG00000164778 | EN2 | chr7:155458128-155464831 | OK | 0.881542 | 0.204841 | -2.10553 | -2.4874 | 5.00E-05 | 0.00107006 | yes |
| ENSG00000185650 | ENSG00000185650 | ZFP36L1 | chr14:68787659-68796253 | OK | 1.87862 | 0.403008 | -2.22079 | -1.91505 | 5.00E-05 | 0.00107006 | yes |
| ENSG00000224597 | ENSG00000224597 | SVIL-AS1 | chr10:29409401-29736781 | OK | 46.4341 | 9.36472 | -2.30988 | -3.65645 | 5.00E-05 | 0.00107006 | yes |
| ENSG00000205609 | ENSG00000205609 | EIF3CL | chr16:28379578-28403879 | OK | 3.26655 | 0.649841 | -2.32961 | -4.14842 | 5.00E-05 | 0.00107006 | yes |
| ENSG00000108244 | ENSG00000108244 | KRT23 | chr17:40921429-40975926 | OK | 1.00126 | 0.181853 | -2.46097 | -1.19376 | 5.00E-05 | 0.00107006 | yes |
| ENSG00000131747 | ENSG00000131747 | TOP2A | chr17:40388515-40417950 | OK | 0.8157 | 0.145773 | -2.48432 | -0.805063 | 5.00E-05 | 0.00107006 | yes |
| ENSG00000227189 | ENSG00000227189 | AC092535.1 | chr4:1113638-1153726 | OK | 0.731284 | 0.129269 | -2.50005 | -2.19044 | 5.00E-05 | 0.00107006 | yes |
| ENSG00000213185 | ENSG00000213185 | FAM24B | chr10:122832148-122879641 | OK | 4.10782 | 0.717341 | -2.51764 | -2.3453 | 5.00E-05 | 0.00107006 | yes |
| ENSG00000133433 | ENSG00000133433 | GSTT2B | chr22:23957413-23961186 | OK | 12.6087 | 2.09385 | -2.59019 | -4.25695 | 5.00E-05 | 0.00107006 | yes |
| ENSG00000157873 | ENSG00000157873 | TNFRSF14 | chr1:2549919-2565382 | OK | 0.633008 | 0.102263 | -2.62994 | -0.609982 | 5.00E-05 | 0.00107006 | yes |
| ENSG00000116016 | ENSG00000116016 | EPAS1 | chr2:46293666-46386703 | OK | 0.706898 | 0.113115 | -2.64372 | -0.717173 | 5.00E-05 | 0.00107006 | yes |

| | | | | | | | | | | | |
|---|---|---|---|---|---|---|---|---|---|---|---|
| ENSG00000255085 | ENSG00000255085 | AF186192.2 | chr8:144700352-144708517 | OK | 8.52748 | 1.3556 | -2.65319 | -3.06663 | 5.00E-05 | 0.00107006 | yes |
| ENSG00000171848 | ENSG00000171848 | RRM2 | chr2:10120697-10211725 | OK | 0.82652 | 0.125464 | -2.71978 | -0.702892 | 5.00E-05 | 0.00107006 | yes |
| ENSG00000283199 | ENSG00000283199 | FP565324.1 | chr13:113953704-113973997 | OK | 0.780665 | 0.117189 | -2.73587 | -1.48422 | 5.00E-05 | 0.00107006 | yes |
| ENSG00000258732 | ENSG00000258732 | AC025884.1 | chr15:22278970-22282872 | OK | 51.7394 | 7.72435 | -2.74378 | -6.29713 | 5.00E-05 | 0.00107006 | yes |
| ENSG00000227124 | ENSG00000227124 | ZNF717 | chr3:75709642-75785583 | OK | 4.98743 | 0.731587 | -2.76919 | -2.60576 | 5.00E-05 | 0.00107006 | yes |
| ENSG00000113209 | ENSG00000113209 | PCDHB5 | chr5:141100241-141249365 | OK | 14.0212 | 1.88622 | -2.89404 | -2.91672 | 5.00E-05 | 0.00107006 | yes |
| ENSG00000163046 | ENSG00000163046 | ANKRD30BL | chr2:132147590-132257969 | OK | 2.16164 | 0.287772 | -2.90913 | -2.59982 | 5.00E-05 | 0.00107006 | yes |
| ENSG00000230333 | ENSG00000230333 | AC004160.1 | chr7:11180901-11832198 | OK | 1.36568 | 0.181165 | -2.91424 | -0.742125 | 5.00E-05 | 0.00107006 | yes |
| ENSG00000188153 | ENSG00000188153 | COL4A5 | chrX:108439843-108697545 | OK | 0.640927 | 0.0698056 | -3.19875 | -0.551213 | 5.00E-05 | 0.00107006 | yes |
| ENSG00000275395 | ENSG00000275395 | FCGBP | chr19:39863322-39906323 | OK | 0.672428 | 0.0699906 | -3.26415 | -2.75247 | 5.00E-05 | 0.00107006 | yes |
| ENSG00000117983 | ENSG00000117983 | MUC5B | chr11:1223065-1262172 | OK | 0.8443 | 0.0861538 | -3.29277 | -0.872444 | 5.00E-05 | 0.00107006 | yes |
| ENSG00000127920 | ENSG00000127920 | GNG11 | chr7:93921698-93928610 | OK | 0.873892 | 0.0874855 | -3.32034 | -2.96084 | 5.00E-05 | 0.00107006 | yes |
| ENSG00000105825 | ENSG00000105825 | TFPI2 | chr7:93591572-93911265 | OK | 5.16122 | 0.469266 | -3.45923 | -2.77495 | 5.00E-05 | 0.00107006 | yes |
| ENSG00000108821 | ENSG00000108821 | COL1A1 | chr17:50183288-50201632 | OK | 4.83775 | 0.415988 | -3.53972 | -2.31869 | 5.00E-05 | 0.00107006 | yes |
| ENSG00000125730 | ENSG00000125730 | C3 | chr19:6677703-6737603 | OK | 2.34916 | 0.181065 | -3.69756 | -0.793921 | 5.00E-05 | 0.00107006 | yes |
| ENSG00000104879 | ENSG00000104879 | CKM | chr19:45306413-45322977 | OK | 2.93067 | 0.211731 | -3.79093 | -3.69716 | 5.00E-05 | 0.00107006 | yes |
| ENSG00000143632 | ENSG00000143632 | ACTA1 | chr1:229431244-229434098 | OK | 5.29051 | 0.346191 | -3.93377 | -4.14098 | 5.00E-05 | 0.00107006 | yes |
| ENSG00000100181 | ENSG00000100181 | TPTEP1 | chr22:16601886-16704477 | OK | 2.19977 | 0.133319 | -4.0444 | -1.21162 | 5.00E-05 | 0.00107006 | yes |
| ENSG00000231473 | ENSG00000231473 | LINC00441 | chr13:48296512-48303661 | OK | 9.10271 | 0.492059 | -4.20939 | -3.85225 | 5.00E-05 | 0.00107006 | yes |
| ENSG00000205663 | ENSG00000205663 | FAM239B | chrX:3891437-3920746 | OK | 7.98673 | 0.371575 | -4.42588 | -3.51499 | 5.00E-05 | 0.00107006 | yes |
| ENSG00000234665 | ENSG00000234665 | AL512625.3 | chr9:62856998-62900104 | OK | 5.81071 | 0.184868 | -4.97414 | -2.30773 | 5.00E-05 | 0.00107006 | yes |
| ENSG00000253731 | ENSG00000253731 | PCDHGA6 | chr5:141330570-141512981 | OK | 10.0797 | 0.107368 | -6.55274 | -1.44927 | 5.00E-05 | 0.00107006 | yes |
| ENSG00000180071 | ENSG00000180071 | ANKRD18A | chr9:38433695-38624990 | OK | 0.231531 | 0.852383 | 1.8803 | 1.15899 | 0.0001 | 0.0019628 | yes |
| ENSG00000152093 | ENSG00000152093 | CFC1B | chr2:130515996-130528604 | OK | 0.348509 | 1.1727 | 1.75057 | 1.57774 | 0.0001 | 0.0019628 | yes |
| ENSG00000106565 | ENSG00000106565 | TMEM176B | chr7:150791284-150805120 | OK | 0.745472 | 2.15911 | 1.53421 | 1.37741 | 0.0001 | 0.0019628 | yes |
| ENSG00000132329 | ENSG00000132329 | RAMP1 | chr2:237858892-237912114 | OK | 0.958502 | 2.76829 | 1.53014 | 1.6717 | 0.0001 | 0.0019628 | yes |
| ENSG00000103175 | ENSG00000103175 | WFDC1 | chr16:84294645-84329851 | OK | 0.587679 | 1.54645 | 1.39586 | 1.58813 | 0.0001 | 0.0019628 | yes |
| ENSG00000151617 | ENSG00000151617 | EDNRA | chr4:147480916-147544954 | OK | 0.319295 | 0.764911 | 1.2604 | 1.08767 | 0.0001 | 0.0019628 | yes |
| ENSG00000008128 | ENSG00000008128 | CDK11A | chr1:1702729-1745992 | OK | 5.87856 | 13.6399 | 1.21429 | 1.40098 | 0.0001 | 0.0019628 | yes |
| ENSG00000213366 | ENSG00000213366 | GSTM2 | chr1:109656080-109709551 | OK | 5.87896 | 12.8979 | 1.1335 | 1.64282 | 0.0001 | 0.0019628 | yes |
| ENSG00000164039 | ENSG00000164039 | BDH2 | chr4:103019867-103099883 | OK | 4.73826 | 10.0773 | 1.08869 | 1.45008 | 0.0001 | 0.0019628 | yes |
| ENSG00000189067 | ENSG00000189067 | LITAF | chr16:11547721-11636381 | OK | 0.98516 | 1.98474 | 1.01052 | 0.800624 | 0.0001 | 0.0019628 | yes |
| ENSG00000182484 | ENSG00000182484 | WASH6P | chrX:155997580-156027877 | OK | 3.22026 | 1.59951 | -1.00955 | -1.33736 | 0.0001 | 0.0019628 | yes |
| ENSG00000215374 | ENSG00000215374 | FAM66B | chr8:7301610-7355354 | OK | 3.29175 | 1.62133 | -1.02168 | -1.726 | 0.0001 | 0.0019628 | yes |
| ENSG00000233198 | ENSG00000233198 | RNF224 | chr9:137227270-137229638 | OK | 2.08265 | 1.00378 | -1.05298 | -1.67795 | 0.0001 | 0.0019628 | yes |
| ENSG00000100154 | ENSG00000100154 | TTC28 | chr22:27851668-28679865 | OK | 28.7839 | 13.535 | -1.08857 | -1.36478 | 0.0001 | 0.0019628 | yes |
| ENSG00000233757 | ENSG00000233757 | AC092835.1 | chr2:95207534-95259774 | OK | 4.06242 | 1.75172 | -1.21356 | -1.546 | 0.0001 | 0.0019628 | yes |
| ENSG00000253771 | ENSG00000253771 | TPTE2P1 | chr13:24924676-24968487 | OK | 6.06768 | 2.48996 | -1.28503 | -1.55948 | 0.0001 | 0.0019628 | yes |
| ENSG00000278249 | ENSG00000278249 | SCARNA2 | chr1:109100192-109100619 | OK | 16.9306 | 6.85305 | -1.30482 | -1.76107 | 0.0001 | 0.0019628 | yes |
| ENSG00000279954 | ENSG00000279954 | BX324167.2 | chr22:45875931-45887748 | OK | 0.649309 | 0.25689 | -1.33776 | -1.69203 | 0.0001 | 0.0019628 | yes |
| ENSG00000147655 | ENSG00000147655 | RSPO2 | chr8:107899315-108083648 | OK | 3.35933 | 1.22135 | -1.45969 | -1.32438 | 0.0001 | 0.0019628 | yes |
| ENSG00000282936 | ENSG00000282936 | AC004706.4 | chr17:6578147-6651634 | OK | 5.78485 | 2.05177 | -1.49541 | -1.81373 | 0.0001 | 0.0019628 | yes |
| ENSG00000104894 | ENSG00000104894 | CD37 | chr19:49331616-49388081 | OK | 0.592831 | 0.206873 | -1.51887 | -0.523815 | 0.0001 | 0.0019628 | yes |
| ENSG00000255052 | ENSG00000255052 | FAM66D | chr8:12115781-12177550 | OK | 5.09386 | 1.43447 | -1.82825 | -1.98983 | 0.0001 | 0.0019628 | yes |
| ENSG00000180178 | ENSG00000180178 | FAR2P1 | chr2:129988157-130051131 | OK | 0.246179 | 32.2427 | 7.03312 | 2.90727 | 0.00015 | 0.00276892 | yes |
| ENSG00000160201 | ENSG00000160201 | U2AF1 | chr21:43092955-43107587 | OK | 3.20765 | 9.21542 | 1.52254 | 1.87944 | 0.00015 | 0.00276892 | yes |
| ENSG00000169894 | ENSG00000169894 | MUC3A | chr7:100949554-100968346 | OK | 1.02523 | 2.7035 | 1.39888 | 1.61585 | 0.00015 | 0.00276892 | yes |

| | | | | | | | | | | | |
|---|---|---|---|---|---|---|---|---|---|---|---|
| ENSG00000120057 | ENSG00000120057 | SFRP5 | chr10:97766750-97771952 | OK | 0.57036 | 1.33538 | 1.22731 | 1.63601 | 0.00015 | 0.00276892 | yes |
| ENSG00000142632 | ENSG00000142632 | ARHGEF19 | chr1:16197853-16212609 | OK | 1.60517 | 3.75045 | 1.22434 | 1.49484 | 0.00015 | 0.00276892 | yes |
| ENSG00000087842 | ENSG00000087842 | PIR | chrX:15384798-15556529 | OK | 1.82164 | 3.93422 | 1.11084 | 1.50606 | 0.00015 | 0.00276892 | yes |
| ENSG00000259316 | ENSG00000259316 | AC087632.1 | chr15:64165516-64387687 | OK | 0.884922 | 0.209843 | -2.07624 | -0.401648 | 0.00015 | 0.00276892 | yes |
| ENSG00000274471 | ENSG00000274471 | AC242376.2 | chr15:23309606-23313276 | OK | 1.61989 | 3.88242 | 1.26106 | 1.75929 | 0.0002 | 0.00351582 | yes |
| ENSG00000128536 | ENSG00000128536 | CDHR3 | chr7:105876795-106033773 | OK | 2.1076 | 4.8205 | 1.19359 | 1.46129 | 0.0002 | 0.00351582 | yes |
| ENSG00000237515 | ENSG00000237515 | SHISA9 | chr16:12901619-13240413 | OK | 2.80085 | 5.75712 | 1.03948 | 1.65212 | 0.0002 | 0.00351582 | yes |
| ENSG00000180574 | ENSG00000180574 | AC068775.1 | chr12:10505601-10523135 | OK | 1.05221 | 2.14402 | 1.02689 | 1.5596 | 0.0002 | 0.00351582 | yes |
| ENSG00000262155 | ENSG00000262155 | LINC02175 | chr16:25066936-25068943 | OK | 1.76437 | 0.854596 | -1.04584 | -1.59021 | 0.0002 | 0.00351582 | yes |
| ENSG00000165929 | ENSG00000165929 | TC2N | chr14:91580695-91867536 | OK | 2.31143 | 1.08731 | -1.08802 | -1.53367 | 0.0002 | 0.00351582 | yes |
| ENSG00000215154 | ENSG00000215154 | AC141586.1 | chr16:2603349-2643296 | OK | 5.52257 | 2.1751 | -1.34426 | -1.49944 | 0.0002 | 0.00351582 | yes |
| ENSG00000144119 | ENSG00000144119 | C1QL2 | chr2:119156242-119158889 | OK | 0.340117 | 0.881654 | 1.37418 | 1.53753 | 0.00025 | 0.00421621 | yes |
| ENSG00000166250 | ENSG00000166250 | CLMP | chr11:123069864-123228277 | OK | 0.932279 | 2.28728 | 1.2948 | 1.24623 | 0.00025 | 0.00421621 | yes |
| ENSG00000173727 | ENSG00000173727 | AP000769.1 | chr11:65455257-65466720 | OK | 2.74358 | 6.1276 | 1.15926 | 2.04227 | 0.00025 | 0.00421621 | yes |
| ENSG00000141510 | ENSG00000141510 | TP53 | chr17:7661778-7703502 | OK | 4.29197 | 2.04565 | -1.06908 | -1.05146 | 0.00025 | 0.00421621 | yes |
| ENSG00000106278 | ENSG00000106278 | PTPRZ1 | chr7:121873088-122062036 | OK | 2.84476 | 1.23963 | -1.19839 | -1.20285 | 0.00025 | 0.00421621 | yes |
| ENSG00000185245 | ENSG00000185245 | GP1BA | chr17:4932296-4935030 | OK | 1.05175 | 0.4395 | -1.25885 | -1.65184 | 0.00025 | 0.00421621 | yes |
| ENSG00000084453 | ENSG00000084453 | SLCO1A2 | chr12:21264599-21419594 | OK | 1.38795 | 0.529504 | -1.39024 | -0.894614 | 0.00025 | 0.00421621 | yes |
| ENSG00000259129 | ENSG00000259129 | LINC00648 | chr14:47764953-47795092 | OK | 1.17155 | 0.428438 | -1.45127 | -1.05061 | 0.00025 | 0.00421621 | yes |
| ENSG00000100842 | ENSG00000100842 | EFS | chr14:23356401-23365752 | OK | 0.573515 | 0.20402 | -1.49112 | -1.62026 | 0.00025 | 0.00421621 | yes |
| ENSG00000189143 | ENSG00000189143 | CLDN4 | chr7:73799541-73832693 | OK | 0.661237 | 0.206053 | -1.68215 | -1.15447 | 0.00025 | 0.00421621 | yes |
| ENSG00000251402 | ENSG00000251402 | FAM90A25P | chr8:12415079-12418090 | OK | 1.12061 | 0.32651 | -1.77908 | -1.80689 | 0.00025 | 0.00421621 | yes |
| ENSG00000125414 | ENSG00000125414 | MYH2 | chr17:10383131-10623886 | OK | 0.696698 | 0.0709771 | -3.29511 | -0.887764 | 0.00025 | 0.00421621 | yes |
| ENSG00000167653 | ENSG00000167653 | PSCA | chr8:142657459-142682724 | OK | 1.66757 | 0.112628 | -3.88811 | -0.869676 | 0.00025 | 0.00421621 | yes |
| ENSG00000182885 | ENSG00000182885 | ADGRG3 | chr16:57668186-57731805 | OK | 0.240879 | 0.728013 | 1.59566 | 1.08459 | 0.0003 | 0.00488215 | yes |
| ENSG00000162009 | ENSG00000162009 | SSTR5 | chr16:1078780-1080142 | OK | 0.656121 | 1.52318 | 1.21505 | 1.51491 | 0.0003 | 0.00488215 | yes |
| ENSG00000171847 | ENSG00000171847 | FAM90A1 | chr12:8221259-8227618 | OK | 1.05541 | 0.481589 | -1.13193 | -1.43026 | 0.0003 | 0.00488215 | yes |
| ENSG00000196296 | ENSG00000196296 | ATP2A1 | chr16:28878404-28939346 | OK | 2.36942 | 0.938757 | -1.33571 | -0.993351 | 0.0003 | 0.00488215 | yes |
| ENSG00000142089 | ENSG00000142089 | IFITM3 | chr11:318639-330122 | OK | 1.79984 | 0.147586 | -3.60824 | -1.27385 | 0.0003 | 0.00488215 | yes |
| ENSG00000009950 | ENSG00000009950 | MLXIPL | chr7:73593193-73624543 | OK | 0.80464 | 2.14224 | 1.4127 | 1.47881 | 0.00035 | 0.00550383 | yes |
| ENSG00000121690 | ENSG00000121690 | DEPDC7 | chr11:33015863-33033582 | OK | 0.451707 | 1.13904 | 1.33435 | 1.46585 | 0.00035 | 0.00550383 | yes |
| ENSG00000273877 | ENSG00000273877 | AC236972.3 | chrX:153225648-153230357 | OK | 0.892761 | 1.97116 | 1.1427 | 1.45282 | 0.00035 | 0.00550383 | yes |
| ENSG00000100526 | ENSG00000100526 | CDKN3 | chr14:54396848-54420218 | OK | 1.33508 | 2.901 | 1.11962 | 1.00486 | 0.00035 | 0.00550383 | yes |
| ENSG00000140443 | ENSG00000140443 | IGF1R | chr15:98648970-99135593 | OK | 25.1086 | 12.2093 | -1.0402 | -1.45513 | 0.00035 | 0.00550383 | yes |
| ENSG00000111679 | ENSG00000111679 | PTPN6 | chr12:6946467-6961316 | OK | 1.1691 | 0.548613 | -1.09154 | -0.583789 | 0.00035 | 0.00550383 | yes |
| ENSG00000224687 | ENSG00000224687 | RASAL2-AS1 | chr1:178091507-178093984 | OK | 1.21082 | 0.433175 | -1.48296 | -1.39209 | 0.00035 | 0.00550383 | yes |
| ENSG00000181449 | ENSG00000181449 | SOX2 | chr3:180983708-181836880 | OK | 3.24709 | 1.07734 | -1.59167 | -1.75152 | 0.00035 | 0.00550383 | yes |
| ENSG00000135346 | ENSG00000135346 | CGA | chr6:87085497-87095406 | OK | 3.19863 | 0.561696 | -2.50959 | -1.74803 | 0.00035 | 0.00550383 | yes |
| ENSG00000125618 | ENSG00000125618 | PAX8 | chr2:113211521-113278950 | OK | 0.026985 | 0.587357 | 4.44401 | 0.304149 | 0.0004 | 0.00614885 | yes |
| ENSG00000189238 | ENSG00000189238 | LINC00943 | chr12:126726269-126772411 | OK | 0.0759736 | 1.06778 | 3.81298 | 0.749081 | 0.0004 | 0.00614885 | yes |
| ENSG00000125731 | ENSG00000125731 | SH2D3A | chr19:6752159-6767588 | OK | 0.278374 | 0.924257 | 1.73127 | 1.07885 | 0.0004 | 0.00614885 | yes |
| ENSG00000072858 | ENSG00000072858 | SIDT1 | chr3:113532295-113629578 | OK | 0.305779 | 0.759952 | 1.31342 | 0.703141 | 0.0004 | 0.00614885 | yes |
| ENSG00000100918 | ENSG00000100918 | REC8 | chr14:24171852-24239242 | OK | 9.68709 | 22.3284 | 1.20475 | 1.16914 | 0.0004 | 0.00614885 | yes |
| ENSG00000213777 | ENSG00000213777 | AC011487.1 | chr19:53503391-53512687 | OK | 6.13113 | 2.21413 | -1.46941 | -1.62708 | 0.0004 | 0.00614885 | yes |
| ENSG00000232815 | ENSG00000232815 | DUX4L50 | chr9:63817747-63818462 | OK | 2.40587 | 0.701335 | -1.77838 | -1.66271 | 0.0004 | 0.00614885 | yes |
| ENSG00000112936 | ENSG00000112936 | C7 | chr5:40909251-40982939 | OK | 0.604941 | 0.168351 | -1.84532 | -0.566881 | 0.0004 | 0.00614885 | yes |
| ENSG00000249158 | ENSG00000249158 | PCDHA11 | chr5:140786135-141012347 | OK | 6.97327 | 1.82988 | -1.93009 | -1.23894 | 0.0004 | 0.00614885 | yes |

| | | | | | | | | | | | | |
|---|---|---|---|---|---|---|---|---|---|---|---|---|
| ENSG00000255974 | ENSG00000255974 | CYP2A6 | chr19:40771647-40900508 | OK | 0.882522 | 0.0797614 | | -3.46787 | -0.508001 | 0.0004 | 0.00614885 | yes |
| ENSG00000280670 | ENSG00000280670 | CCDC163 | chr1:45493865-45523047 | OK | 0.402607 | 2.18157 | | 2.43792 | 1.13025 | 0.00045 | 0.00675493 | yes |
| ENSG00000183486 | ENSG00000183486 | MX2 | chr21:41361942-41409390 | OK | 0.26198 | 0.843085 | | 1.68622 | 0.762942 | 0.00045 | 0.00675493 | yes |
| ENSG00000149571 | ENSG00000149571 | KIRREL3 | chr11:126355639-127006058 | OK | 1.18369 | 2.60652 | | 1.13883 | 1.18187 | 0.00045 | 0.00675493 | yes |
| ENSG00000114439 | ENSG00000114439 | BBX | chr3:107522935-107811324 | OK | 5.12269 | 2.53541 | | -1.01468 | -1.13459 | 0.00045 | 0.00675493 | yes |
| ENSG00000258186 | ENSG00000258186 | SLC7A5P2 | chr16:21402236-21520444 | OK | 21.4948 | 7.69057 | | -1.48283 | -1.77259 | 0.00045 | 0.00675493 | yes |
| ENSG00000279875 | ENSG00000279875 | AC074029.4 | chr12:48189348-48191207 | OK | 0.927268 | 0.330449 | | -1.48856 | -1.66372 | 0.00045 | 0.00675493 | yes |
| ENSG00000170231 | ENSG00000170231 | FABP6 | chr5:160187366-160238735 | OK | 1.21055 | 3.66331 | | 1.59749 | 1.3495 | 0.0005 | 0.00734648 | yes |
| ENSG00000174332 | ENSG00000174332 | GLIS1 | chr1:53506236-53738106 | OK | 0.299447 | 0.6793 | | 1.18175 | 1.46813 | 0.0005 | 0.00734648 | yes |
| ENSG00000114013 | ENSG00000114013 | CD86 | chr3:122055365-122121139 | OK | 0.87039 | 0.413148 | | -1.075 | -0.867111 | 0.0005 | 0.00734648 | yes |
| ENSG00000125148 | ENSG00000125148 | MT2A | chr16:56608198-56609497 | OK | 2.13825 | 0.848752 | | -1.33301 | -1.21465 | 0.0005 | 0.00734648 | yes |
| ENSG00000143768 | ENSG00000143768 | LEFTY2 | chr1:225936410-225941489 | OK | 0.633762 | 0.120857 | | -2.39065 | -1.31831 | 0.0005 | 0.00734648 | yes |
| ENSG00000127666 | ENSG00000127666 | TICAM1 | chr19:4815931-4831704 | OK | 0.17758 | 0.558642 | | 1.65346 | 1.47652 | 0.00055 | 0.0079124 | yes |
| ENSG00000197540 | ENSG00000197540 | GZMM | chr19:544033-549924 | OK | 1.73852 | 0.502421 | | -1.79089 | -1.7461 | 0.00055 | 0.0079124 | yes |
| ENSG00000253873 | ENSG00000253873 | PCDHGA11 | chr5:141330570-141512981 | OK | 2.66915 | 0.320967 | | -3.05588 | -0.896899 | 0.00055 | 0.0079124 | yes |
| ENSG00000268790 | ENSG00000268790 | AC008764.4 | chr19:16479056-16660442 | OK | 0.726925 | 0.0700467 | | -3.37542 | -0.165737 | 0.00055 | 0.0079124 | yes |
| ENSG00000093100 | ENSG00000093100 | AC016026.1 | chr22:17787648-18024559 | OK | 0.830404 | 7.34549 | | 3.14497 | 2.05576 | 0.0006 | 0.00848922 | yes |
| ENSG00000256463 | ENSG00000256463 | SALL3 | chr18:78980274-79002677 | OK | 0.318842 | 0.992086 | | 1.63762 | 0.834326 | 0.0006 | 0.00848922 | yes |
| ENSG00000177406 | ENSG00000177406 | AC021054.1 | chr12:564295-663779 | OK | 0.184286 | 0.567407 | | 1.62244 | 1.21965 | 0.0006 | 0.00848922 | yes |
| ENSG00000197714 | ENSG00000197714 | ZNF460 | chr19:57267219-57294469 | OK | 7.69024 | 3.8239 | | -1.00798 | -1.50558 | 0.0006 | 0.00848922 | yes |
| ENSG00000173889 | ENSG00000173889 | PHC3 | chr3:170086731-170181749 | OK | 10.7817 | 5.11458 | | -1.0759 | -1.29616 | 0.0006 | 0.00848922 | yes |
| ENSG00000230183 | ENSG00000230183 | CNOT6LP1 | chr15:56005714-56007176 | OK | 1.0932 | 0.45803 | | -1.25504 | -1.42847 | 0.0006 | 0.00848922 | yes |
| ENSG00000117013 | ENSG00000117013 | KCNQ4 | chr1:40784011-40840452 | OK | 0.354539 | 0.728523 | | 1.03903 | 1.39862 | 0.00065 | 0.00905328 | yes |
| ENSG00000198468 | ENSG00000198468 | FLVCR1-AS1 | chr1:212852107-212858088 | OK | 2.93277 | 1.30173 | | -1.17183 | -1.34687 | 0.00065 | 0.00905328 | yes |
| ENSG00000214273 | ENSG00000214273 | AGGF1P1 | chr4:190041341-190043414 | OK | 0.549356 | 0.142569 | | -1.94609 | -1.72157 | 0.00065 | 0.00905328 | yes |
| ENSG00000172058 | ENSG00000172058 | SERF1A | chr5:70900664-70918530 | OK | 0.465201 | 1.04238 | | 1.16395 | 0.904842 | 0.0007 | 0.00960252 | yes |
| ENSG00000135750 | ENSG00000135750 | KCNK1 | chr1:233614003-233672512 | OK | 0.657931 | 1.47288 | | 1.16263 | 1.17691 | 0.0007 | 0.00960252 | yes |
| ENSG00000228330 | ENSG00000228330 | AL355355.1 | chr10:13142184-13142977 | OK | 1.68891 | 0.477069 | | -1.82383 | -1.64566 | 0.0007 | 0.00960252 | yes |

| test_id | gene_id | gene | locus | status | AHN | SNCA | log2(fold_change) | test_stat | p_value | q_value | significant |
|---|---|---|---|---|---|---|---|---|---|---|---|
| ENSG00000140598 | ENSG00000140598 | EFL1 | chr15:82130229-82262763 | OK | 4.14493 | 156.874 | 5.24212 | 13.7887 | 5.00E-05 | 0.00107006 | yes |
| ENSG00000237424 | ENSG00000237424 | FOXD2-AS1 | chr1:47432132-47434641 | OK | 0.109593 | 2.60391 | 4.57045 | 4.44413 | 5.00E-05 | 0.00107006 | yes |
| ENSG00000186564 | ENSG00000186564 | FOXD2 | chr1:47436016-47440691 | OK | 0.107405 | 2.43992 | 4.5057 | 5.55158 | 5.00E-05 | 0.00107006 | yes |
| ENSG00000227617 | ENSG00000227617 | CERS6-AS1 | chr2:168455861-168913371 | OK | 0.35822 | 7.54796 | 4.39717 | 1.0593 | 5.00E-05 | 0.00107006 | yes |
| ENSG00000189238 | ENSG00000189238 | LINC00943 | chr12:126726269-126772411 | OK | 0.0759736 | 1.29703 | 4.09357 | 0.824963 | 0.0004 | 0.00614885 | yes |
| ENSG00000112246 | ENSG00000112246 | SIM1 | chr6:100385014-100464929 | OK | 0.121461 | 1.87056 | 3.94491 | 2.44858 | 5.00E-05 | 0.00107006 | yes |
| ENSG00000170044 | ENSG00000170044 | ZPLD1 | chr3:102099243-102479841 | OK | 0.154168 | 2.32756 | 3.91624 | 2.62867 | 5.00E-05 | 0.00107006 | yes |
| ENSG00000146469 | ENSG00000146469 | VIP | chr6:152750797-152759765 | OK | 0.816407 | 8.56891 | 3.39175 | 5.40327 | 5.00E-05 | 0.00107006 | yes |
| ENSG00000109991 | ENSG00000109991 | P2RX3 | chr11:57338373-57370600 | OK | 0.158026 | 1.64696 | 3.38158 | 1.92634 | 5.00E-05 | 0.00107006 | yes |
| ENSG00000120907 | ENSG00000120907 | ADRA1A | chr8:26748149-26867273 | OK | 0.0653301 | 0.624479 | 3.25683 | 0.803045 | 0.0002 | 0.00351582 | yes |
| ENSG00000189127 | ENSG00000189127 | ANKRD34B | chr5:80556754-80570488 | OK | 0.0887271 | 0.761661 | 3.1017 | 1.61716 | 5.00E-05 | 0.00107006 | yes |
| ENSG00000114200 | ENSG00000114200 | BCHE | chr3:165772903-165837472 | OK | 0.60155 | 4.13483 | 2.78107 | 2.69162 | 5.00E-05 | 0.00107006 | yes |
| ENSG00000181195 | ENSG00000181195 | PENK | chr8:56436673-56559823 | OK | 0.941177 | 6.46594 | 2.78032 | 2.92068 | 5.00E-05 | 0.00107006 | yes |
| ENSG00000141750 | ENSG00000141750 | STAC2 | chr17:39210535-39225872 | OK | 0.752849 | 4.7308 | 2.65165 | 5.33403 | 5.00E-05 | 0.00107006 | yes |
| ENSG00000128283 | ENSG00000128283 | CDC42EP1 | chr22:37560446-37569405 | OK | 0.246997 | 1.55052 | 2.65019 | 1.99165 | 5.00E-05 | 0.00107006 | yes |
| ENSG00000146122 | ENSG00000146122 | DAAM2 | chr6:39792297-39934551 | OK | 0.526982 | 2.9726 | 2.4959 | 2.50023 | 5.00E-05 | 0.00107006 | yes |
| ENSG00000140945 | ENSG00000140945 | CDH13 | chr16:82626802-83807834 | OK | 3.5617 | 19.7499 | 2.47121 | 3.4572 | 5.00E-05 | 0.00107006 | yes |
| ENSG00000054803 | ENSG00000054803 | CBLN4 | chr20:55997439-56005472 | OK | 0.170019 | 0.9035 | 2.40983 | 2.78915 | 5.00E-05 | 0.00107006 | yes |
| ENSG00000065675 | ENSG00000065675 | PRKCQ | chr10:6427142-6580301 | OK | 0.181627 | 0.951516 | 2.38925 | 2.73028 | 5.00E-05 | 0.00107006 | yes |
| ENSG00000147488 | ENSG00000147488 | ST18 | chr8:52110838-52460959 | OK | 0.186612 | 0.96939 | 2.37703 | 0.760978 | 5.00E-05 | 0.00107006 | yes |
| ENSG00000120669 | ENSG00000120669 | SOHLH2 | chr13:36168207-36297842 | OK | 0.315417 | 1.60292 | 2.34537 | 1.93259 | 0.00015 | 0.00276892 | yes |
| ENSG00000100427 | ENSG00000100427 | MLC1 | chr22:50059390-50085902 | OK | 0.241339 | 1.17711 | 2.28612 | 1.215 | 5.00E-05 | 0.00107006 | yes |
| ENSG00000152527 | ENSG00000152527 | PLEKHH2 | chr2:43637272-43767987 | OK | 1.09549 | 5.32497 | 2.2812 | 2.59076 | 5.00E-05 | 0.00107006 | yes |
| ENSG00000168952 | ENSG00000168952 | STXBP6 | chr14:24809655-25050297 | OK | 1.3362 | 5.98452 | 2.1631 | 4.40086 | 5.00E-05 | 0.00107006 | yes |
| ENSG00000178038 | ENSG00000178038 | ALS2CL | chr3:46668996-46693704 | OK | 0.35044 | 1.54858 | 2.1437 | 2.26676 | 5.00E-05 | 0.00107006 | yes |
| ENSG00000197747 | ENSG00000197747 | S100A10 | chr1:151982914-151994390 | OK | 2.14494 | 9.45593 | 2.14028 | 2.7548 | 5.00E-05 | 0.00107006 | yes |
| ENSG00000150457 | ENSG00000150457 | LATS2 | chr13:20973031-21061547 | OK | 1.40858 | 6.19324 | 2.13645 | 3.41897 | 5.00E-05 | 0.00107006 | yes |
| ENSG00000279926 | ENSG00000279926 | AL138831.3 | chr6:3982672-3984130 | OK | 0.286039 | 1.25356 | 2.13174 | 2.11735 | 5.00E-05 | 0.00107006 | yes |
| ENSG00000107317 | ENSG00000107317 | PTGDS | chr9:136975091-136986410 | OK | 2.05501 | 8.94445 | 2.12185 | 2.47405 | 5.00E-05 | 0.00107006 | yes |
| ENSG00000152689 | ENSG00000152689 | RASGRP3 | chr2:33436323-33564750 | OK | 1.27168 | 5.50755 | 2.11468 | 2.06039 | 5.00E-05 | 0.00107006 | yes |
| ENSG00000183054 | ENSG00000183054 | RGPD6 | chr2:110513811-110577185 | OK | 0.38669 | 1.66748 | 2.10842 | 1.28284 | 5.00E-05 | 0.00107006 | yes |
| ENSG00000152583 | ENSG00000152583 | SPARCL1 | chr4:87473334-87531061 | OK | 0.349115 | 1.49214 | 2.09561 | 1.12279 | 0.00025 | 0.00421621 | yes |
| ENSG00000233621 | ENSG00000233621 | LINC01137 | chr1:37454878-37474411 | OK | 0.210401 | 0.898476 | 2.09434 | 1.81556 | 0.00025 | 0.00421621 | yes |
| ENSG00000170893 | ENSG00000170893 | TRH | chr3:129974304-129977938 | OK | 3.83414 | 16.1942 | 2.0785 | 5.42775 | 5.00E-05 | 0.00107006 | yes |
| ENSG00000249859 | ENSG00000249859 | PVT1 | chr8:127794532-128101253 | OK | 1.84958 | 7.58011 | 2.03502 | 1.71391 | 5.00E-05 | 0.00107006 | yes |
| ENSG00000162998 | ENSG00000162998 | FRZB | chr2:182833274-182867162 | OK | 0.146006 | 0.589016 | 2.01228 | 2.10322 | 5.00E-05 | 0.00107006 | yes |
| ENSG00000152214 | ENSG00000152214 | RIT2 | chr18:42743226-43115691 | OK | 10.3868 | 41.8298 | 2.00978 | 5.63609 | 5.00E-05 | 0.00107006 | yes |
| ENSG00000137573 | ENSG00000137573 | SULF1 | chr8:69466623-69660915 | OK | 2.60418 | 10.2675 | 1.97919 | 1.76073 | 5.00E-05 | 0.00107006 | yes |
| ENSG00000174607 | ENSG00000174607 | UGT8 | chr4:114598454-114678224 | OK | 0.156597 | 0.613548 | 1.97012 | 1.61188 | 5.00E-05 | 0.00107006 | yes |
| ENSG00000278532 | ENSG00000278532 | AC026585.1 | chr18:69398725-69399249 | OK | 0.986235 | 3.85833 | 1.96797 | 1.71334 | 0.00035 | 0.00550383 | yes |
| ENSG00000112561 | ENSG00000112561 | TFEB | chr6:41683977-41736259 | OK | 0.408288 | 1.58297 | 1.95497 | 0.875738 | 5.00E-05 | 0.00107006 | yes |
| ENSG00000125968 | ENSG00000125968 | ID1 | chr20:31605282-31606515 | OK | 4.3627 | 16.9059 | 1.95423 | 4.19016 | 5.00E-05 | 0.00107006 | yes |
| ENSG00000135406 | ENSG00000135406 | PRPH | chr12:49292630-49331731 | OK | 0.357354 | 1.37818 | 1.94734 | 1.28727 | 5.00E-05 | 0.00107006 | yes |
| ENSG00000115507 | ENSG00000115507 | OTX1 | chr2:63050056-63057836 | OK | 0.379802 | 1.45407 | 1.93678 | 1.69977 | 5.00E-05 | 0.00107006 | yes |
| ENSG00000113805 | ENSG00000113805 | CNTN3 | chr3:74262567-74521140 | OK | 0.227378 | 0.860062 | 1.91935 | 1.25066 | 0.00035 | 0.00550383 | yes |

| | | | | | | | | | | | |
|---|---|---|---|---|---|---|---|---|---|---|---|
| ENSG00000198075 | ENSG00000198075 | SULT1C4 | chr2:108377910-108388057 | OK | 1.46054 | 5.48738 | 1.90962 | 3.0093 | 5.00E-05 | 0.00107006 | yes |
| ENSG00000168743 | ENSG00000168743 | NPNT | chr4:105552619-106022478 | OK | 13.1701 | 49.3046 | 1.90446 | 5.07628 | 5.00E-05 | 0.00107006 | yes |
| ENSG00000278959 | ENSG00000278959 | AC006378.2 | chr7:93954043-93956519 | OK | 0.287012 | 1.05405 | 1.87676 | 2.31024 | 5.00E-05 | 0.00107006 | yes |
| ENSG00000237943 | ENSG00000237943 | PRKCQ-AS1 | chr10:6580418-6616452 | OK | 0.42174 | 1.52619 | 1.85551 | 0.978261 | 5.00E-05 | 0.00107006 | yes |
| ENSG00000157445 | ENSG00000157445 | CACNA2D3 | chr3:54122546-55074557 | OK | 2.69052 | 9.66419 | 1.84476 | 3.20134 | 5.00E-05 | 0.00107006 | yes |
| ENSG00000178401 | ENSG00000178401 | DNAJC22 | chr12:49346916-49357546 | OK | 0.183334 | 0.635011 | 1.7923 | 0.979358 | 0.0002 | 0.00351582 | yes |
| ENSG00000146674 | ENSG00000146674 | IGFBP3 | chr7:45912244-45921874 | OK | 0.927386 | 3.20203 | 1.78775 | 2.36131 | 5.00E-05 | 0.00107006 | yes |
| ENSG00000258498 | ENSG00000258498 | DIO3OS | chr14:101552220-101560431 | OK | 0.610981 | 2.09804 | 1.77985 | 1.47719 | 5.00E-05 | 0.00107006 | yes |
| ENSG00000109255 | ENSG00000109255 | NMU | chr4:55595228-55636698 | OK | 1.58611 | 5.41589 | 1.7717 | 2.2578 | 5.00E-05 | 0.00107006 | yes |
| ENSG00000278266 | ENSG00000278266 | AC079949.2 | chr12:127147148-127150081 | OK | 0.997401 | 3.39196 | 1.76587 | 3.56127 | 5.00E-05 | 0.00107006 | yes |
| ENSG00000196843 | ENSG00000196843 | ARID5A | chr2:96536742-96552638 | OK | 0.36285 | 1.21996 | 1.74938 | 1.82015 | 5.00E-05 | 0.00107006 | yes |
| ENSG00000144119 | ENSG00000144119 | C1QL2 | chr2:119156242-119158889 | OK | 0.340117 | 1.14127 | 1.74653 | 2.01874 | 5.00E-05 | 0.00107006 | yes |
| ENSG00000182993 | ENSG00000182993 | C12orf60 | chr12:14803571-14906586 | OK | 0.916797 | 3.07176 | 1.74439 | 1.85291 | 5.00E-05 | 0.00107006 | yes |
| ENSG00000233639 | ENSG00000233639 | LINC01158 | chr2:104807571-104853183 | OK | 0.856514 | 2.85998 | 1.73946 | 1.59528 | 5.00E-05 | 0.00107006 | yes |
| ENSG00000100302 | ENSG00000100302 | RASD2 | chr22:35540867-35554001 | OK | 1.55308 | 5.1709 | 1.73528 | 4.26839 | 5.00E-05 | 0.00107006 | yes |
| ENSG00000150630 | ENSG00000150630 | VEGFC | chr4:176669620-176792727 | OK | 0.232754 | 0.774876 | 1.73516 | 1.54367 | 5.00E-05 | 0.00107006 | yes |
| ENSG00000144331 | ENSG00000144331 | ZNF385B | chr2:179441981-179861505 | OK | 0.422701 | 1.4038 | 1.73163 | 1.07399 | 5.00E-05 | 0.00107006 | yes |
| ENSG00000132329 | ENSG00000132329 | RAMP1 | chr2:237858892-237912114 | OK | 0.958502 | 3.168 | 1.72472 | 1.86594 | 5.00E-05 | 0.00107006 | yes |
| ENSG00000204381 | ENSG00000204381 | LAYN | chr11:111540279-111561745 | OK | 0.241822 | 0.79472 | 1.7165 | 0.86103 | 0.0002 | 0.00351582 | yes |
| ENSG00000258175 | ENSG00000258175 | LINC02300 | chr14:28592222-28613623 | OK | 1.2739 | 4.18008 | 1.71428 | 1.9228 | 0.0001 | 0.0019628 | yes |
| ENSG00000176769 | ENSG00000176769 | TCERG1L | chr10:131092390-131311721 | OK | 0.274039 | 0.898003 | 1.71234 | 2.11382 | 5.00E-05 | 0.00107006 | yes |
| ENSG00000189056 | ENSG00000189056 | RELN | chr7:103297421-103989516 | OK | 2.27522 | 7.39732 | 1.701 | 1.75835 | 5.00E-05 | 0.00107006 | yes |
| ENSG00000183036 | ENSG00000183036 | PCP4 | chr21:39867316-39929397 | OK | 12.8401 | 41.0923 | 1.67822 | 4.11776 | 5.00E-05 | 0.00107006 | yes |
| ENSG00000164778 | ENSG00000164778 | EN2 | chr7:155458128-155464831 | OK | 0.881542 | 2.80829 | 1.67159 | 3.18051 | 5.00E-05 | 0.00107006 | yes |
| ENSG00000138378 | ENSG00000138378 | STAT4 | chr2:191021525-191151596 | OK | 1.03762 | 3.26525 | 1.65391 | 1.89976 | 5.00E-05 | 0.00107006 | yes |
| ENSG00000221818 | ENSG00000221818 | EBF2 | chr8:25841729-26045397 | OK | 0.181697 | 0.569915 | 1.64921 | 1.29822 | 0.0003 | 0.00488215 | yes |
| ENSG00000260578 | ENSG00000260578 | AC110597.1 | chr18:67481790-67484966 | OK | 0.292219 | 0.904464 | 1.63001 | 2.20047 | 5.00E-05 | 0.00107006 | yes |
| ENSG00000117318 | ENSG00000117318 | ID3 | chr1:23557917-23559794 | OK | 2.23275 | 6.88541 | 1.62472 | 2.47303 | 5.00E-05 | 0.00107006 | yes |
| ENSG00000104888 | ENSG00000104888 | SLC17A7 | chr19:49429400-49442360 | OK | 0.493272 | 1.50966 | 1.61377 | 2.30792 | 5.00E-05 | 0.00107006 | yes |
| ENSG00000122863 | ENSG00000122863 | CHST3 | chr10:71964364-72013564 | OK | 1.05401 | 3.1872 | 1.5964 | 4.3124 | 5.00E-05 | 0.00107006 | yes |
| ENSG00000168135 | ENSG00000168135 | KCNJ4 | chr22:38424287-38455199 | OK | 2.74004 | 8.28525 | 1.59635 | 3.88846 | 5.00E-05 | 0.00107006 | yes |
| ENSG00000104375 | ENSG00000104375 | STK3 | chr8:98401402-98942827 | OK | 0.234567 | 0.708884 | 1.59555 | 0.552212 | 5.00E-05 | 0.00107006 | yes |
| ENSG00000175766 | ENSG00000175766 | EIF4E1B | chr5:176630681-176646641 | OK | 0.860528 | 2.53237 | 1.5572 | 1.2286 | 0.00045 | 0.00675493 | yes |
| ENSG00000152229 | ENSG00000152229 | PSTPIP2 | chr18:45983535-46072272 | OK | 0.412172 | 1.20659 | 1.54962 | 1.55638 | 5.00E-05 | 0.00107006 | yes |
| ENSG00000157404 | ENSG00000157404 | KIT | chr4:54657917-54740715 | OK | 1.11498 | 3.2431 | 1.54035 | 2.33141 | 5.00E-05 | 0.00107006 | yes |
| ENSG00000109846 | ENSG00000109846 | CRYAB | chr11:111908564-111926872 | OK | 6.89256 | 19.9443 | 1.53287 | 3.06205 | 5.00E-05 | 0.00107006 | yes |
| ENSG00000184160 | ENSG00000184160 | ADRA2C | chr4:3766347-3768526 | OK | 3.80479 | 10.9154 | 1.52047 | 4.05528 | 5.00E-05 | 0.00107006 | yes |
| ENSG00000254656 | ENSG00000254656 | RTL1 | chr14:100779409-101027415 | OK | 1.34571 | 3.84889 | 1.51607 | 3.57128 | 5.00E-05 | 0.00107006 | yes |
| ENSG00000134376 | ENSG00000134376 | CRB1 | chr1:197268203-197478455 | OK | 0.418312 | 1.19305 | 1.51201 | 1.41299 | 5.00E-05 | 0.00107006 | yes |
| ENSG00000275763 | ENSG00000275763 | C18orf65 | chr18:76495520-76498088 | OK | 0.232812 | 0.663315 | 1.51053 | 1.56712 | 0.0002 | 0.00351582 | yes |
| ENSG00000131914 | ENSG00000131914 | LIN28A | chr1:26410777-26429722 | OK | 0.573054 | 1.62214 | 1.50116 | 2.65834 | 5.00E-05 | 0.00107006 | yes |
| ENSG00000120318 | ENSG00000120318 | ARAP3 | chr5:141653400-141682221 | OK | 1.91448 | 5.4068 | 1.49783 | 3.07937 | 5.00E-05 | 0.00107006 | yes |
| ENSG00000186897 | ENSG00000186897 | C1QL4 | chr12:49332410-49337188 | OK | 3.88448 | 10.826 | 1.47871 | 3.92564 | 5.00E-05 | 0.00107006 | yes |
| ENSG00000197405 | ENSG00000197405 | C5AR1 | chr19:47290022-47322066 | OK | 0.342387 | 0.949563 | 1.47163 | 1.29288 | 5.00E-05 | 0.00107006 | yes |
| ENSG00000158089 | ENSG00000158089 | GALNT14 | chr2:30910466-31155202 | OK | 1.34903 | 3.73366 | 1.46867 | 1.61815 | 5.00E-05 | 0.00107006 | yes |
| ENSG00000143768 | ENSG00000143768 | LEFTY2 | chr1:225936410-225941489 | OK | 0.633762 | 1.7519 | 1.46691 | 1.82674 | 5.00E-05 | 0.00107006 | yes |

| | | | | | | | | | | | | |
|---|---|---|---|---|---|---|---|---|---|---|---|---|
| ENSG00000178075 | ENSG00000178075 | GRAMD1C | chr3:113828181-113947570 | OK | 1.27038 | 3.50987 | | 1.46616 | 1.24029 | 0.0001 | 0.0019628 | yes |
| ENSG00000258818 | ENSG00000258818 | RNASE4 | chr14:20684099-20707120 | OK | 1.33642 | 3.68804 | | 1.46448 | 1.8507 | 5.00E-05 | 0.00107006 | yes |
| ENSG00000277954 | ENSG00000277954 | AC092376.2 | chr16:78099412-79212667 | OK | 0.51281 | 1.40324 | | 1.45227 | 1.96597 | 5.00E-05 | 0.00107006 | yes |
| ENSG00000149571 | ENSG00000149571 | KIRREL3 | chr11:126355639-127006058 | OK | 1.18369 | 3.22793 | | 1.44731 | 1.41936 | 5.00E-05 | 0.00107006 | yes |
| ENSG00000178573 | ENSG00000178573 | MAF | chr16:79585842-79600714 | OK | 1.44039 | 3.92055 | | 1.4446 | 2.10681 | 5.00E-05 | 0.00107006 | yes |
| ENSG00000152137 | ENSG00000152137 | HSPB8 | chr12:119178641-119221131 | OK | 0.457178 | 1.2418 | | 1.4416 | 1.57581 | 5.00E-05 | 0.00107006 | yes |
| ENSG00000152661 | ENSG00000152661 | GJA1 | chr6:121435691-121449727 | OK | 0.211075 | 0.572911 | | 1.44056 | 1.59727 | 0.0003 | 0.00488215 | yes |
| ENSG00000142102 | ENSG00000142102 | PGGHG | chr11:289134-296107 | OK | 1.1411 | 3.09657 | | 1.44024 | 1.56233 | 5.00E-05 | 0.00107006 | yes |
| ENSG00000163273 | ENSG00000163273 | NPPC | chr2:231921819-231926403 | OK | 2.96095 | 8.0301 | | 1.43936 | 2.55682 | 5.00E-05 | 0.00107006 | yes |
| ENSG00000081052 | ENSG00000081052 | COL4A4 | chr2:227002710-227164113 | OK | 0.400856 | 1.07831 | | 1.42762 | 3.13325 | 5.00E-05 | 0.00107006 | yes |
| ENSG00000114346 | ENSG00000114346 | ECT2 | chr3:172750681-172821474 | OK | 0.579545 | 1.55675 | | 1.42555 | 1.20089 | 5.00E-05 | 0.00107006 | yes |
| ENSG00000283709 | ENSG00000283709 | FAM238C | chr10:26931205-26944418 | OK | 0.724941 | 1.93534 | | 1.41665 | 1.33141 | 0.00025 | 0.00421621 | yes |
| ENSG00000104327 | ENSG00000104327 | CALB1 | chr8:90058607-90095475 | OK | 27.7843 | 73.332 | | 1.40017 | 5.76048 | 5.00E-05 | 0.00107006 | yes |
| ENSG00000171124 | ENSG00000171124 | FUT3 | chr19:5842887-5858239 | OK | 1.50184 | 3.96301 | | 1.39986 | 1.63019 | 5.00E-05 | 0.00107006 | yes |
| ENSG00000187091 | ENSG00000187091 | PLCD1 | chr3:38007495-38029762 | OK | 4.87008 | 12.8253 | | 1.39697 | 2.67941 | 5.00E-05 | 0.00107006 | yes |
| ENSG00000172345 | ENSG00000172345 | STARD5 | chr15:81159574-81324183 | OK | 1.44889 | 3.81275 | | 1.39589 | 1.559 | 5.00E-05 | 0.00107006 | yes |
| ENSG00000088992 | ENSG00000088992 | TESC | chr12:117038922-117099479 | OK | 1.18556 | 3.11641 | | 1.39432 | 1.61958 | 5.00E-05 | 0.00107006 | yes |
| ENSG00000070159 | ENSG00000070159 | PTPN3 | chr9:109375465-109498313 | OK | 1.14185 | 2.99935 | | 1.39328 | 1.98087 | 5.00E-05 | 0.00107006 | yes |
| ENSG00000169692 | ENSG00000169692 | AGPAT2 | chr9:136673142-136687423 | OK | 0.681595 | 1.78316 | | 1.38745 | 1.68953 | 5.00E-05 | 0.00107006 | yes |
| ENSG00000113749 | ENSG00000113749 | HRH2 | chr5:175658029-175710756 | OK | 0.330403 | 0.863109 | | 1.38532 | 1.58176 | 0.0003 | 0.00488215 | yes |
| ENSG00000113100 | ENSG00000113100 | CDH9 | chr5:26880599-27121150 | OK | 3.03873 | 7.93728 | | 1.38518 | 1.96449 | 5.00E-05 | 0.00107006 | yes |
| ENSG00000130158 | ENSG00000130158 | DOCK6 | chr19:11199294-11262481 | OK | 4.19864 | 10.9631 | | 1.38466 | 3.11993 | 5.00E-05 | 0.00107006 | yes |
| ENSG00000116014 | ENSG00000116014 | KISS1R | chr19:917286-921015 | OK | 1.22653 | 3.19238 | | 1.38006 | 1.41874 | 0.00035 | 0.00550383 | yes |
| ENSG00000176907 | ENSG00000176907 | TCIM | chr8:40153454-40155308 | OK | 1.10437 | 2.84791 | | 1.36668 | 2.32751 | 5.00E-05 | 0.00107006 | yes |
| ENSG00000008441 | ENSG00000008441 | NFIX | chr19:12995607-13098796 | OK | 0.259318 | 0.668597 | | 1.36641 | 0.643136 | 0.00065 | 0.00905328 | yes |
| ENSG00000115556 | ENSG00000115556 | PLCD4 | chr2:218607764-218637184 | OK | 1.0927 | 2.81555 | | 1.36552 | 1.64601 | 5.00E-05 | 0.00107006 | yes |
| ENSG00000076706 | ENSG00000076706 | MCAM | chr11:119206275-119321521 | OK | 6.81953 | 17.5501 | | 1.36374 | 1.50632 | 5.00E-05 | 0.00107006 | yes |
| ENSG00000278916 | ENSG00000278916 | CEP83-AS1 | chr12:94460002-94462484 | OK | 0.575687 | 1.47592 | | 1.35825 | 2.07317 | 5.00E-05 | 0.00107006 | yes |
| ENSG00000122254 | ENSG00000122254 | HS3ST2 | chr16:22814176-22916338 | OK | 1.81524 | 4.63646 | | 1.35287 | 2.93333 | 5.00E-05 | 0.00107006 | yes |
| ENSG00000137507 | ENSG00000137507 | LRRC32 | chr11:76657055-76670747 | OK | 0.401286 | 1.02132 | | 1.34774 | 1.47045 | 5.00E-05 | 0.00107006 | yes |
| ENSG00000124508 | ENSG00000124508 | BTN2A2 | chr6:26383095-26394874 | OK | 1.056 | 2.68731 | | 1.34756 | 1.47836 | 5.00E-05 | 0.00107006 | yes |
| ENSG00000172554 | ENSG00000172554 | SNTG2 | chr2:950867-1367613 | OK | 0.708411 | 1.80269 | | 1.34749 | 0.863914 | 0.0005 | 0.00734648 | yes |
| ENSG00000162817 | ENSG00000162817 | C1orf115 | chr1:220689844-220699157 | OK | 0.553619 | 1.40704 | | 1.34569 | 2.27301 | 5.00E-05 | 0.00107006 | yes |
| ENSG00000120903 | ENSG00000120903 | CHRNA2 | chr8:27459760-27479883 | OK | 0.769314 | 1.95096 | | 1.34254 | 0.832665 | 0.00015 | 0.00276892 | yes |
| ENSG00000169894 | ENSG00000169894 | MUC3A | chr7:100949554-100968346 | OK | 1.02523 | 2.59907 | | 1.34205 | 1.50365 | 0.0003 | 0.00488215 | yes |
| ENSG00000214274 | ENSG00000214274 | ANG | chr14:20684099-20707120 | OK | 0.92362 | 2.33938 | | 1.34075 | 1.25838 | 0.00055 | 0.0079124 | yes |
| ENSG00000143507 | ENSG00000143507 | DUSP10 | chr1:221701423-221742176 | OK | 1.51689 | 3.83984 | | 1.33993 | 2.2725 | 5.00E-05 | 0.00107006 | yes |
| ENSG00000005981 | ENSG00000005981 | ASB4 | chr7:95478443-95540232 | OK | 1.12353 | 2.83252 | | 1.33405 | 2.0951 | 5.00E-05 | 0.00107006 | yes |
| ENSG00000239467 | ENSG00000239467 | AC007405.3 | chr2:170771112-170778148 | OK | 1.40464 | 3.52276 | | 1.32651 | 1.51325 | 0.0006 | 0.00848922 | yes |
| ENSG00000187676 | ENSG00000187676 | B3GLCT | chr13:31199935-31332276 | OK | 0.61455 | 1.53572 | | 1.32131 | 2.29222 | 5.00E-05 | 0.00107006 | yes |
| ENSG00000121690 | ENSG00000121690 | DEPDC7 | chr11:33015863-33033582 | OK | 0.451707 | 1.12153 | | 1.312 | 1.447 | 0.0003 | 0.00488215 | yes |
| ENSG00000105355 | ENSG00000105355 | PLIN3 | chr19:4838331-4867768 | OK | 2.74343 | 6.8034 | | 1.31027 | 2.80578 | 5.00E-05 | 0.00107006 | yes |
| ENSG00000181444 | ENSG00000181444 | ZNF467 | chr7:149764181-149773479 | OK | 1.46505 | 3.62286 | | 1.30618 | 2.19744 | 5.00E-05 | 0.00107006 | yes |
| ENSG00000103196 | ENSG00000103196 | CRISPLD2 | chr16:84819983-84920768 | OK | 0.489297 | 1.20674 | | 1.30234 | 1.29729 | 5.00E-05 | 0.00107006 | yes |
| ENSG00000133636 | ENSG00000133636 | NTS | chr12:85874294-85882992 | OK | 3.19161 | 7.85692 | | 1.29968 | 2.61699 | 5.00E-05 | 0.00107006 | yes |
| ENSG00000110628 | ENSG00000110628 | SLC22A18 | chr11:2887779-2925246 | OK | 2.96468 | 7.28498 | | 1.29705 | 1.89926 | 5.00E-05 | 0.00107006 | yes |

| | | | | | | | | | | | |
|---|---|---|---|---|---|---|---|---|---|---|---|
| ENSG00000118271 | ENSG00000118271 | TTR | chr18:31591725-31599021 | OK | 3.2583 | 7.93937 | 1.28491 | 2.06628 | 5.00E-05 | 0.00107006 | yes |
| ENSG00000075651 | ENSG00000075651 | PLD1 | chr3:171600404-171810950 | OK | 0.38845 | 0.946482 | 1.28485 | 0.638883 | 0.0004 | 0.00614885 | yes |
| ENSG00000231249 | ENSG00000231249 | ITPR1-AS1 | chr3:4490890-4493163 | OK | 0.833427 | 2.02086 | 1.27784 | 1.37296 | 0.0006 | 0.00848922 | yes |
| ENSG00000143178 | ENSG00000143178 | TBX19 | chr1:168281039-168314426 | OK | 0.267892 | 0.649298 | 1.27723 | 1.40183 | 0.00035 | 0.00550383 | yes |
| ENSG00000267731 | ENSG00000267731 | AC005332.5 | chr17:68189883-68192802 | OK | 0.722094 | 1.74864 | 1.27598 | 2.19831 | 5.00E-05 | 0.00107006 | yes |
| ENSG00000116183 | ENSG00000116183 | PAPPA2 | chr1:176463170-176845605 | OK | 1.44296 | 3.49244 | 1.27521 | 1.26949 | 0.0002 | 0.00351582 | yes |
| ENSG00000130711 | ENSG00000130711 | PRDM12 | chr9:130664593-130682981 | OK | 0.367116 | 0.888533 | 1.27519 | 1.64468 | 0.00015 | 0.00276892 | yes |
| ENSG00000186193 | ENSG00000186193 | SAPCD2 | chr9:137057662-137070588 | OK | 0.637661 | 1.54186 | 1.27381 | 2.25394 | 5.00E-05 | 0.00107006 | yes |
| ENSG00000100593 | ENSG00000100593 | ISM2 | chr14:77474393-77498850 | OK | 0.558759 | 1.35066 | 1.27336 | 1.4331 | 5.00E-05 | 0.00107006 | yes |
| ENSG00000279601 | ENSG00000279601 | AC005052.1 | chrX:119693567-119696059 | OK | 0.371201 | 0.895776 | 1.27094 | 1.67181 | 5.00E-05 | 0.00107006 | yes |
| ENSG00000171476 | ENSG00000171476 | HOPX | chr4:56647987-56681899 | OK | 1.10643 | 2.66744 | 1.26954 | 1.6394 | 5.00E-05 | 0.00107006 | yes |
| ENSG00000148357 | ENSG00000148357 | HMCN2 | chr9:130265881-130434123 | OK | 0.965214 | 2.32558 | 1.26867 | 1.67239 | 5.00E-05 | 0.00107006 | yes |
| ENSG00000125733 | ENSG00000125733 | TRIP10 | chr19:6737924-6751526 | OK | 1.96443 | 4.73267 | 1.26854 | 1.7911 | 5.00E-05 | 0.00107006 | yes |
| ENSG00000117152 | ENSG00000117152 | RGS4 | chr1:163068774-163076802 | OK | 27.9958 | 66.8623 | 1.25599 | 2.96472 | 5.00E-05 | 0.00107006 | yes |
| ENSG00000197406 | ENSG00000197406 | DIO3 | chr14:101561350-101563452 | OK | 1.62909 | 3.8801 | 1.25203 | 2.50576 | 5.00E-05 | 0.00107006 | yes |
| ENSG00000064666 | ENSG00000064666 | CNN2 | chr19:1026580-1039068 | OK | 5.6224 | 13.3827 | 1.2511 | 3.33758 | 5.00E-05 | 0.00107006 | yes |
| ENSG00000105173 | ENSG00000105173 | CCNE1 | chr19:29811897-29824308 | OK | 1.82084 | 4.32522 | 1.24817 | 2.15836 | 5.00E-05 | 0.00107006 | yes |
| ENSG00000162989 | ENSG00000162989 | KCNJ3 | chr2:154698298-154858352 | OK | 1.03431 | 2.45255 | 1.24562 | 1.40142 | 0.00045 | 0.00675493 | yes |
| ENSG00000115380 | ENSG00000115380 | EFEMP1 | chr2:55865966-55924139 | OK | 0.481846 | 1.13967 | 1.24197 | 0.787565 | 0.00015 | 0.00276892 | yes |
| ENSG00000130950 | ENSG00000130950 | NUTM2F | chr9:94318195-94328644 | OK | 0.272981 | 0.639264 | 1.22761 | 1.42748 | 0.0002 | 0.00351582 | yes |
| ENSG00000103966 | ENSG00000103966 | EHD4 | chr15:41828084-41972578 | OK | 1.07698 | 2.51514 | 1.22365 | 1.57613 | 5.00E-05 | 0.00107006 | yes |
| ENSG00000031691 | ENSG00000031691 | CENPQ | chr6:49463377-49493107 | OK | 0.744237 | 1.73071 | 1.21753 | 1.75267 | 5.00E-05 | 0.00107006 | yes |
| ENSG00000227908 | ENSG00000227908 | FLJ31104 | chr5:55995166-56003649 | OK | 0.413707 | 0.96123 | 1.21627 | 1.55578 | 0.0002 | 0.00351582 | yes |
| ENSG00000115738 | ENSG00000115738 | ID2 | chr2:8666635-8684453 | OK | 11.5628 | 26.8463 | 1.21523 | 2.68294 | 5.00E-05 | 0.00107006 | yes |
| ENSG00000250241 | ENSG00000250241 | AC105383.1 | chr4:133075310-133149116 | OK | 2.0855 | 4.83595 | 1.21341 | 2.4131 | 5.00E-05 | 0.00107006 | yes |
| ENSG00000146054 | ENSG00000146054 | TRIM7 | chr5:181191923-181205293 | OK | 5.60726 | 12.9494 | 1.20752 | 2.62199 | 5.00E-05 | 0.00107006 | yes |
| ENSG00000169891 | ENSG00000169891 | REPS2 | chrX:16946690-17153280 | OK | 0.949547 | 2.19096 | 1.20625 | 1.72987 | 0.0001 | 0.0019628 | yes |
| ENSG00000230269 | ENSG00000230269 | LINC02525 | chr6:3182743-3195756 | OK | 4.71861 | 10.8673 | 1.20356 | 2.99352 | 5.00E-05 | 0.00107006 | yes |
| ENSG00000120549 | ENSG00000120549 | KIAA1217 | chr10:23694745-24547848 | OK | 0.592728 | 1.36199 | 1.20028 | 1.1279 | 0.0007 | 0.00960252 | yes |
| ENSG00000189266 | ENSG00000189266 | PNRC2 | chr1:23959108-23963462 | OK | 20.999 | 48.2483 | 1.20016 | 5.14229 | 5.00E-05 | 0.00107006 | yes |
| ENSG00000283312 | ENSG00000283312 | AC017104.4 | chr2:231493707-231495299 | OK | 0.815605 | 1.87324 | 1.19959 | 1.70549 | 5.00E-05 | 0.00107006 | yes |
| ENSG00000171621 | ENSG00000171621 | SPSB1 | chr1:9292879-9369532 | OK | 1.57754 | 3.62318 | 1.19958 | 1.81762 | 5.00E-05 | 0.00107006 | yes |
| ENSG00000166002 | ENSG00000166002 | SMCO4 | chr11:93478471-93543508 | OK | 8.60712 | 19.7354 | 1.19718 | 1.70155 | 0.00025 | 0.00421621 | yes |
| ENSG00000168994 | ENSG00000168994 | PXDC1 | chr6:3722613-3753871 | OK | 1.4934 | 3.41754 | 1.19436 | 1.94252 | 5.00E-05 | 0.00107006 | yes |
| ENSG00000121905 | ENSG00000121905 | HPCA | chr1:32885993-32901438 | OK | 4.11839 | 9.41357 | 1.19266 | 1.7412 | 5.00E-05 | 0.00107006 | yes |
| ENSG00000173559 | ENSG00000173559 | NABP1 | chr2:191678067-191696659 | OK | 0.846543 | 1.92997 | 1.18892 | 1.31979 | 0.0005 | 0.00734648 | yes |
| ENSG00000171729 | ENSG00000171729 | TMEM51 | chr1:15152531-15220480 | OK | 0.814732 | 1.85736 | 1.18886 | 1.6915 | 5.00E-05 | 0.00107006 | yes |
| ENSG00000135218 | ENSG00000135218 | CD36 | chr7:80312573-80679277 | OK | 0.501191 | 1.13952 | 1.18499 | 0.66236 | 0.00045 | 0.00675493 | yes |
| ENSG00000179399 | ENSG00000179399 | GPC5 | chr13:91398606-92873682 | OK | 1.04365 | 2.37184 | 1.18437 | 1.39753 | 5.00E-05 | 0.00107006 | yes |
| ENSG00000155367 | ENSG00000155367 | PPM1J | chr1:112701105-112715477 | OK | 5.10448 | 11.5636 | 1.17976 | 2.33869 | 5.00E-05 | 0.00107006 | yes |
| ENSG00000163932 | ENSG00000163932 | PRKCD | chr3:53156008-53192717 | OK | 3.14086 | 7.09274 | 1.17518 | 2.93255 | 5.00E-05 | 0.00107006 | yes |
| ENSG00000101198 | ENSG00000101198 | NKAIN4 | chr20:63240783-63272694 | OK | 4.27332 | 9.63355 | 1.17271 | 2.57729 | 5.00E-05 | 0.00107006 | yes |
| ENSG00000188338 | ENSG00000188338 | SLC38A3 | chr3:50205245-50221486 | OK | 0.734022 | 1.65442 | 1.17243 | 1.06879 | 0.0002 | 0.00351582 | yes |
| ENSG00000186493 | ENSG00000186493 | C5orf38 | chr5:2752130-2755397 | OK | 21.6577 | 48.808 | 1.17224 | 2.11641 | 5.00E-05 | 0.00107006 | yes |
| ENSG00000072274 | ENSG00000072274 | TFRC | chr3:196027182-196082189 | OK | 17.2922 | 38.8833 | 1.16903 | 3.2045 | 5.00E-05 | 0.00107006 | yes |
| ENSG00000117643 | ENSG00000117643 | MAN1C1 | chr1:25617467-25786207 | OK | 1.76029 | 3.95279 | 1.16705 | 1.75152 | 5.00E-05 | 0.00107006 | yes |

| | | | | | | | | | | | | |
|---|---|---|---|---|---|---|---|---|---|---|---|---|
| ENSG00000177606 | ENSG00000177606 | JUN | chr1:58780787-58784327 | OK | 10.1456 | 22.738 | | 1.16425 | 4.64435 | 5.00E-05 | 0.00107006 | yes |
| ENSG00000175183 | ENSG00000175183 | CSRP2 | chr12:76858714-76880352 | OK | 31.2915 | 70.0362 | | 1.16233 | 4.06055 | 5.00E-05 | 0.00107006 | yes |
| ENSG00000156413 | ENSG00000156413 | FUT6 | chr19:5830609-5839731 | OK | 0.414426 | 0.927051 | | 1.16153 | 0.830808 | 0.00055 | 0.0079124 | yes |
| ENSG00000260941 | ENSG00000260941 | LINC00622 | chr1:119597701-119599271 | OK | 1.58008 | 3.50367 | | 1.14887 | 2.0346 | 5.00E-05 | 0.00107006 | yes |
| ENSG00000112893 | ENSG00000112893 | MAN2A1 | chr5:109689365-109869625 | OK | 2.40481 | 5.32007 | | 1.14552 | 2.23155 | 5.00E-05 | 0.00107006 | yes |
| ENSG00000170340 | ENSG00000170340 | B3GNT2 | chr2:62196112-62224731 | OK | 2.09137 | 4.61537 | | 1.14199 | 2.7348 | 5.00E-05 | 0.00107006 | yes |
| ENSG00000181524 | ENSG00000181524 | RPL24P4 | chr6:42956344-42956765 | OK | 29.4895 | 65.072 | | 1.14184 | 2.62863 | 5.00E-05 | 0.00107006 | yes |
| ENSG00000171189 | ENSG00000171189 | GRIK1 | chr21:29193479-29940033 | OK | 4.68943 | 10.3313 | | 1.13954 | 1.89485 | 5.00E-05 | 0.00107006 | yes |
| ENSG00000021645 | ENSG00000021645 | NRXN3 | chr14:78170372-79868290 | OK | 7.91636 | 17.3991 | | 1.1361 | 3.34277 | 5.00E-05 | 0.00107006 | yes |
| ENSG00000128573 | ENSG00000128573 | FOXP2 | chr7:114086326-114693772 | OK | 9.89369 | 21.7378 | | 1.13562 | 2.85795 | 5.00E-05 | 0.00107006 | yes |
| ENSG00000163032 | ENSG00000163032 | VSNL1 | chr2:17539125-17657018 | OK | 18.2899 | 40.1124 | | 1.133 | 3.57637 | 5.00E-05 | 0.00107006 | yes |
| ENSG00000073605 | ENSG00000073605 | GSDMB | chr17:39904594-39919854 | OK | 5.01164 | 10.9551 | | 1.12824 | 2.16975 | 5.00E-05 | 0.00107006 | yes |
| ENSG00000260526 | ENSG00000260526 | AC109347.1 | chr4:112229560-112231596 | OK | 1.16875 | 2.54116 | | 1.12052 | 1.97606 | 5.00E-05 | 0.00107006 | yes |
| ENSG00000131941 | ENSG00000131941 | RHPN2 | chr19:32978592-33064888 | OK | 1.33253 | 2.89378 | | 1.11879 | 1.38288 | 0.00015 | 0.00276892 | yes |
| ENSG00000132692 | ENSG00000132692 | BCAN | chr1:156641389-156661424 | OK | 0.75195 | 1.62209 | | 1.10915 | 1.35303 | 0.0001 | 0.0019628 | yes |
| ENSG00000241494 | ENSG00000241494 | AL355032.1 | chr14:101634453-101732522 | OK | 10.2978 | 22.1955 | | 1.10793 | 1.86604 | 5.00E-05 | 0.00107006 | yes |
| ENSG00000132196 | ENSG00000132196 | HSD17B7 | chr1:162790701-162812817 | OK | 15.5531 | 33.498 | | 1.10687 | 2.90047 | 5.00E-05 | 0.00107006 | yes |
| ENSG00000087253 | ENSG00000087253 | LPCAT2 | chr16:55508997-55586670 | OK | 2.31686 | 4.96755 | | 1.10036 | 1.4928 | 5.00E-05 | 0.00107006 | yes |
| ENSG00000117971 | ENSG00000117971 | CHRNB4 | chr15:78624118-78735495 | OK | 2.005 | 4.29277 | | 1.09831 | 2.28805 | 5.00E-05 | 0.00107006 | yes |
| ENSG00000240376 | ENSG00000240376 | AC010343.1 | chr5:32925638-33297910 | OK | 11.706 | 24.977 | | 1.09335 | 2.41443 | 5.00E-05 | 0.00107006 | yes |
| ENSG00000237515 | ENSG00000237515 | SHISA9 | chr16:12901619-13240413 | OK | 2.80085 | 5.95674 | | 1.08866 | 1.94231 | 5.00E-05 | 0.00107006 | yes |
| ENSG00000171119 | ENSG00000171119 | NRTN | chr19:5823801-5828324 | OK | 1.86494 | 3.96624 | | 1.08865 | 1.76094 | 5.00E-05 | 0.00107006 | yes |
| ENSG00000076003 | ENSG00000076003 | MCM6 | chr2:135839625-135876426 | OK | 2.58355 | 5.48495 | | 1.08612 | 2.93847 | 5.00E-05 | 0.00107006 | yes |
| ENSG00000236279 | ENSG00000236279 | CLEC2L | chr7:139523855-139544984 | OK | 3.02763 | 6.42067 | | 1.08454 | 2.08531 | 5.00E-05 | 0.00107006 | yes |
| ENSG00000151812 | ENSG00000151812 | SLC35F4 | chr14:57563921-57982194 | OK | 0.904534 | 1.91819 | | 1.0845 | 1.29737 | 5.00E-05 | 0.00107006 | yes |
| ENSG00000090539 | ENSG00000090539 | CHRD | chr3:184380072-184390736 | OK | 2.17088 | 4.59533 | | 1.08189 | 1.54678 | 5.00E-05 | 0.00107006 | yes |
| ENSG00000101190 | ENSG00000101190 | TCFL5 | chr20:62816243-62861763 | OK | 2.8444 | 6.00006 | | 1.07686 | 2.66091 | 5.00E-05 | 0.00107006 | yes |
| ENSG00000130830 | ENSG00000130830 | MPP1 | chrX:154778683-154821007 | OK | 3.79179 | 7.98024 | | 1.07355 | 2.10503 | 5.00E-05 | 0.00107006 | yes |
| ENSG00000284642 | ENSG00000284642 | AL139424.2 | chr1:10395415-10397432 | OK | 5.25577 | 11.0361 | | 1.07026 | 2.01738 | 5.00E-05 | 0.00107006 | yes |
| ENSG00000154640 | ENSG00000154640 | BTG3 | chr21:17593652-17633199 | OK | 6.25867 | 13.1214 | | 1.068 | 2.43448 | 5.00E-05 | 0.00107006 | yes |
| ENSG00000147571 | ENSG00000147571 | CRH | chr8:66176381-66178725 | OK | 13.5973 | 28.4989 | | 1.06759 | 3.61704 | 5.00E-05 | 0.00107006 | yes |
| ENSG00000158201 | ENSG00000158201 | ABHD3 | chr18:21650896-21704805 | OK | 2.9406 | 6.1607 | | 1.06698 | 2.1209 | 5.00E-05 | 0.00107006 | yes |
| ENSG00000111907 | ENSG00000111907 | TPD52L1 | chr6:125119048-125302078 | OK | 5.09043 | 10.6608 | | 1.06646 | 1.39171 | 0.0007 | 0.00960252 | yes |
| ENSG00000088280 | ENSG00000088280 | ASAP3 | chr1:23428562-23484568 | OK | 7.79573 | 16.2329 | | 1.05817 | 2.7563 | 5.00E-05 | 0.00107006 | yes |
| ENSG00000234745 | ENSG00000234745 | HLA-B | chr6:31268748-31357637 | OK | 8.64657 | 17.9473 | | 1.05357 | 2.09921 | 5.00E-05 | 0.00107006 | yes |
| ENSG00000134215 | ENSG00000134215 | VAV3 | chr1:107571159-107994607 | OK | 2.55407 | 5.28959 | | 1.05036 | 1.5893 | 5.00E-05 | 0.00107006 | yes |
| ENSG00000260075 | ENSG00000260075 | NSFP1 | chr17:46372854-46487141 | OK | 1.77069 | 3.66343 | | 1.04888 | 1.8034 | 5.00E-05 | 0.00107006 | yes |
| ENSG00000119514 | ENSG00000119514 | GALNT12 | chr9:98807698-98872415 | OK | 2.45079 | 5.05513 | | 1.0445 | 1.36182 | 0.0006 | 0.00848922 | yes |
| ENSG00000162616 | ENSG00000162616 | DNAJB4 | chr1:77979174-78138449 | OK | 8.6428 | 17.8212 | | 1.04402 | 2.48587 | 5.00E-05 | 0.00107006 | yes |
| ENSG00000145506 | ENSG00000145506 | NKD2 | chr5:1008828-1038943 | OK | 9.25233 | 19.076 | | 1.04387 | 2.36889 | 5.00E-05 | 0.00107006 | yes |
| ENSG00000044115 | ENSG00000044115 | CTNNA1 | chr5:138610966-138935034 | OK | 54.1461 | 111.547 | | 1.04273 | 2.23551 | 5.00E-05 | 0.00107006 | yes |
| ENSG00000158445 | ENSG00000158445 | KCNB1 | chr20:49219294-49484297 | OK | 8.47171 | 17.4191 | | 1.03995 | 2.95981 | 5.00E-05 | 0.00107006 | yes |
| ENSG00000153012 | ENSG00000153012 | LGI2 | chr4:24998846-25030879 | OK | 13.6603 | 28.0359 | | 1.03728 | 4.61402 | 5.00E-05 | 0.00107006 | yes |
| ENSG00000131669 | ENSG00000131669 | NINJ1 | chr9:93121488-93134288 | OK | 9.90033 | 20.2959 | | 1.03564 | 2.9809 | 5.00E-05 | 0.00107006 | yes |
| ENSG00000151929 | ENSG00000151929 | BAG3 | chr10:119651369-119677819 | OK | 1.25417 | 2.56357 | | 1.03142 | 1.94766 | 5.00E-05 | 0.00107006 | yes |
| ENSG00000205922 | ENSG00000205922 | ONECUT3 | chr19:1752372-1780988 | OK | 3.70692 | 7.57102 | | 1.03027 | 3.90686 | 5.00E-05 | 0.00107006 | yes |

| | | | | | | | | | | | |
|---|---|---|---|---|---|---|---|---|---|---|---|
| ENSG00000139354 | ENSG00000139354 | GAS2L3 | chr12:100573682-100628286 | OK | 2.20418 | 4.49596 | 1.02839 | 1.89862 | 5.00E-05 | 0.00107006 | yes |
| ENSG00000138653 | ENSG00000138653 | NDST4 | chr4:114827762-115113876 | OK | 0.384675 | 0.78381 | 1.02687 | 1.3195 | 0.00025 | 0.00421621 | yes |
| ENSG00000177551 | ENSG00000177551 | NHLH2 | chr1:115836376-115843917 | OK | 2.91362 | 5.93555 | 1.02657 | 2.33421 | 5.00E-05 | 0.00107006 | yes |
| ENSG00000146592 | ENSG00000146592 | CREB5 | chr7:28299320-28825894 | OK | 3.33369 | 6.77854 | 1.02385 | 1.31921 | 5.00E-05 | 0.00107006 | yes |
| ENSG00000128335 | ENSG00000128335 | APOL2 | chr22:36226202-36239954 | OK | 2.97059 | 6.03548 | 1.02272 | 1.83654 | 5.00E-05 | 0.00107006 | yes |
| ENSG00000273148 | ENSG00000273148 | AL035563.1 | chr20:18794528-18796067 | OK | 1.34475 | 2.7317 | 1.02246 | 1.67754 | 5.00E-05 | 0.00107006 | yes |
| ENSG00000130956 | ENSG00000130956 | HABP4 | chr9:96450200-96619830 | OK | 33.2833 | 67.5748 | 1.02169 | 4.20349 | 5.00E-05 | 0.00107006 | yes |
| ENSG00000120833 | ENSG00000120833 | SOCS2 | chr12:93542462-93583487 | OK | 2.65014 | 5.35793 | 1.01561 | 1.67676 | 5.00E-05 | 0.00107006 | yes |
| ENSG00000127922 | ENSG00000127922 | SEM1 | chr7:96481625-96709891 | OK | 21.9184 | 44.1569 | 1.0105 | 1.34845 | 0.00055 | 0.0079124 | yes |
| ENSG00000166033 | ENSG00000166033 | HTRA1 | chr10:122461524-122514908 | OK | 5.79528 | 11.671 | 1.00998 | 2.47565 | 5.00E-05 | 0.00107006 | yes |
| ENSG00000125354 | ENSG00000125354 | 6-Sep | chrX:119615723-119693370 | OK | 18.9739 | 38.196 | 1.0094 | 3.91547 | 5.00E-05 | 0.00107006 | yes |
| ENSG00000151892 | ENSG00000151892 | GFRA1 | chr10:116056924-116273467 | OK | 11.1671 | 22.4068 | 1.00467 | 1.3453 | 5.00E-05 | 0.00107006 | yes |
| ENSG00000156504 | ENSG00000156504 | FAM122B | chrX:134769565-134854610 | OK | 7.59317 | 15.2232 | 1.00349 | 2.41205 | 5.00E-05 | 0.00107006 | yes |
| ENSG00000164070 | ENSG00000164070 | HSPA4L | chr4:127781820-127844040 | OK | 12.1514 | 24.322 | 1.00114 | 3.06587 | 5.00E-05 | 0.00107006 | yes |
| ENSG00000054967 | ENSG00000054967 | RELT | chr11:73376263-73397474 | OK | 8.68002 | 17.3612 | 1.0001 | 3.47866 | 5.00E-05 | 0.00107006 | yes |
| ENSG00000242498 | ENSG00000242498 | ARPIN | chr15:89830598-89912956 | OK | 7.60342 | 3.80008 | -1.00062 | -1.26194 | 5.00E-05 | 0.00107006 | yes |
| ENSG00000153207 | ENSG00000153207 | AHCTF1 | chr1:246771836-246931978 | OK | 16.6648 | 8.31348 | -1.00328 | -1.45864 | 5.00E-05 | 0.00107006 | yes |
| ENSG00000135976 | ENSG00000135976 | ANKRD36 | chr2:97113495-97264521 | OK | 13.1583 | 6.56137 | -1.0039 | -2.03926 | 5.00E-05 | 0.00107006 | yes |
| ENSG00000282458 | ENSG00000282458 | WASH5P | chr19:60950-71626 | OK | 3.91996 | 1.95067 | -1.00687 | -1.1735 | 0.0006 | 0.00848922 | yes |
| ENSG00000048540 | ENSG00000048540 | LMO3 | chr12:16347141-16610594 | OK | 160.802 | 80.0056 | -1.00711 | -3.49464 | 5.00E-05 | 0.00107006 | yes |
| ENSG00000178965 | ENSG00000178965 | ERICH3 | chr1:74568110-74673738 | OK | 1.1954 | 0.594474 | -1.0078 | -1.56868 | 0.00015 | 0.00276892 | yes |
| ENSG00000159905 | ENSG00000159905 | ZNF221 | chr19:43951222-43967709 | OK | 3.83031 | 1.90153 | -1.0103 | -1.87848 | 5.00E-05 | 0.00107006 | yes |
| ENSG00000269918 | ENSG00000269918 | AF131215.6 | chr8:10896044-11201366 | OK | 6.76507 | 3.35706 | -1.0109 | -1.68803 | 5.00E-05 | 0.00107006 | yes |
| ENSG00000108106 | ENSG00000108106 | UBE2S | chr19:55385344-55407777 | OK | 83.6794 | 41.494 | -1.01197 | -1.48772 | 0.00025 | 0.00421621 | yes |
| ENSG00000161681 | ENSG00000161681 | SHANK1 | chr19:50661826-50719450 | OK | 58.5191 | 29.0167 | -1.01202 | -1.67882 | 0.0002 | 0.00351582 | yes |
| ENSG00000130338 | ENSG00000130338 | TULP4 | chr6:158232235-158511828 | OK | 52.8138 | 26.1191 | -1.01581 | -1.46833 | 0.0001 | 0.0019628 | yes |
| ENSG00000185339 | ENSG00000185339 | TCN2 | chr22:30576624-30627278 | OK | 11.8526 | 5.85668 | -1.01705 | -1.62086 | 5.00E-05 | 0.00107006 | yes |
| ENSG00000274422 | ENSG00000274422 | AC245060.5 | chr22:22283927-22287220 | OK | 1.96689 | 0.9703 | -1.01941 | -2.00872 | 5.00E-05 | 0.00107006 | yes |
| ENSG00000092098 | ENSG00000092098 | RNF31 | chr14:24143361-24167402 | OK | 34.5676 | 17.0504 | -1.01961 | -1.58573 | 5.00E-05 | 0.00107006 | yes |
| ENSG00000170396 | ENSG00000170396 | ZNF804A | chr2:184593576-184939492 | OK | 3.88755 | 1.91037 | -1.02501 | -2.75381 | 5.00E-05 | 0.00107006 | yes |
| ENSG00000115274 | ENSG00000115274 | INO80B | chr2:74455022-74460891 | OK | 39.5188 | 19.4116 | -1.02562 | -1.9556 | 5.00E-05 | 0.00107006 | yes |
| ENSG00000140365 | ENSG00000140365 | COMMD4 | chr15:75335890-75343224 | OK | 16.983 | 8.32375 | -1.02879 | -2.12265 | 5.00E-05 | 0.00107006 | yes |
| ENSG00000164330 | ENSG00000164330 | EBF1 | chr5:158695915-159099761 | OK | 21.2704 | 10.4129 | -1.03048 | -2.13689 | 5.00E-05 | 0.00107006 | yes |
| ENSG00000127399 | ENSG00000127399 | LRRC61 | chr7:150243915-150338150 | OK | 12.71 | 6.21654 | -1.03179 | -2.54153 | 5.00E-05 | 0.00107006 | yes |
| ENSG00000237517 | ENSG00000237517 | DGCR5 | chr22:18970513-19031242 | OK | 3.9769 | 1.9403 | -1.03536 | -2.15437 | 5.00E-05 | 0.00107006 | yes |
| ENSG00000185900 | ENSG00000185900 | POMK | chr8:43093505-43123434 | OK | 25.103 | 12.2403 | -1.03622 | -3.40123 | 5.00E-05 | 0.00107006 | yes |
| ENSG00000281508 | ENSG00000281508 | AL078639.1 | chrX:140782404-140784871 | OK | 6.53189 | 3.17623 | -1.04018 | -1.68176 | 0.00015 | 0.00276892 | yes |
| ENSG00000137841 | ENSG00000137841 | PLCB2 | chr15:40278175-40307935 | OK | 3.52941 | 1.70876 | -1.04648 | -1.30225 | 0.0003 | 0.00488215 | yes |
| ENSG00000182674 | ENSG00000182674 | KCNB2 | chr8:72537390-72938349 | OK | 3.61009 | 1.7463 | -1.04773 | -2.56219 | 5.00E-05 | 0.00107006 | yes |
| ENSG00000174243 | ENSG00000174243 | DDX23 | chr12:48829763-48852842 | OK | 41.5529 | 20.0953 | -1.04809 | -2.37318 | 5.00E-05 | 0.00107006 | yes |
| ENSG00000112182 | ENSG00000112182 | BACH2 | chr6:89926528-90296908 | OK | 24.9172 | 12.0432 | -1.04893 | -2.00421 | 5.00E-05 | 0.00107006 | yes |
| ENSG00000128965 | ENSG00000128965 | CHAC1 | chr15:40952961-40956519 | OK | 39.5536 | 19.0285 | -1.05565 | -3.79047 | 5.00E-05 | 0.00107006 | yes |
| ENSG00000180626 | ENSG00000180626 | ZNF594 | chr17:5179535-5191883 | OK | 5.37034 | 2.57819 | -1.05865 | -1.70844 | 5.00E-05 | 0.00107006 | yes |
| ENSG00000140937 | ENSG00000140937 | CDH11 | chr16:64943752-65126112 | OK | 13.8899 | 6.66342 | -1.0597 | -1.71617 | 5.00E-05 | 0.00107006 | yes |
| ENSG00000257354 | ENSG00000257354 | AC048341.2 | chr12:62601750-62622213 | OK | 0.622557 | 0.298431 | -1.06081 | -1.78821 | 5.00E-05 | 0.00107006 | yes |
| ENSG00000111834 | ENSG00000111834 | RSPH4A | chr6:116616478-116632985 | OK | 1.18773 | 0.568993 | -1.06173 | -1.54579 | 0.00025 | 0.00421621 | yes |

| | | | | | | | | | | | | |
|---|---|---|---|---|---|---|---|---|---|---|---|---|
| ENSG00000204175 | ENSG00000204175 | GPRIN2 | chr10:46549043-46555530 | OK | 5.04886 | 2.41712 | | -1.06267 | -2.20725 | 5.00E-05 | 0.00107006 | yes |
| ENSG00000281490 | ENSG00000281490 | CICP14 | chr7:128655961-128658791 | OK | 12.1363 | 5.80871 | | -1.06304 | -3.46733 | 5.00E-05 | 0.00107006 | yes |
| ENSG00000273079 | ENSG00000273079 | GRIN2B | chr12:13437941-13981957 | OK | 14.409 | 6.87722 | | -1.06708 | -1.79867 | 5.00E-05 | 0.00107006 | yes |
| ENSG00000162068 | ENSG00000162068 | NTN3 | chr16:2471498-2474145 | OK | 1.61993 | 0.771356 | | -1.07046 | -1.62393 | 0.0001 | 0.0019628 | yes |
| ENSG00000269313 | ENSG00000269313 | MAGIX | chrX:49162563-49168483 | OK | 5.4103 | 2.57568 | | -1.07076 | -1.37466 | 0.0005 | 0.00734648 | yes |
| ENSG00000132434 | ENSG00000132434 | LANCL2 | chr7:55365447-55433742 | OK | 20.1535 | 9.55409 | | -1.07684 | -3.88066 | 5.00E-05 | 0.00107006 | yes |
| ENSG00000260855 | ENSG00000260855 | AL591848.4 | chr1:246771836-246931978 | OK | 4.26762 | 2.02013 | | -1.07898 | -1.61061 | 0.00025 | 0.00421621 | yes |
| ENSG00000029534 | ENSG00000029534 | ANK1 | chr8:41653219-41896762 | OK | 30.0287 | 14.1112 | | -1.0895 | -4.0425 | 5.00E-05 | 0.00107006 | yes |
| ENSG00000168959 | ENSG00000168959 | GRM5 | chr11:88504575-89065945 | OK | 0.998584 | 0.468982 | | -1.09035 | -1.29458 | 0.00015 | 0.00276892 | yes |
| ENSG00000279255 | ENSG00000279255 | Z97653.2 | chr16:707649-709067 | OK | 1.4164 | 0.662698 | | -1.09581 | -1.36329 | 0.0004 | 0.00614885 | yes |
| ENSG00000265179 | ENSG00000265179 | AP000894.2 | chr18:894434-912172 | OK | 10.8739 | 5.0812 | | -1.09762 | -1.77 | 5.00E-05 | 0.00107006 | yes |
| ENSG00000241852 | ENSG00000241852 | C8orf58 | chr8:22578278-22604150 | OK | 8.0318 | 3.75023 | | -1.09875 | -2.41248 | 5.00E-05 | 0.00107006 | yes |
| ENSG00000141519 | ENSG00000141519 | CCDC40 | chr17:80036631-80100613 | OK | 21.7256 | 10.1259 | | -1.10135 | -2.17437 | 5.00E-05 | 0.00107006 | yes |
| ENSG00000159263 | ENSG00000159263 | SIM2 | chr21:36698772-36749917 | OK | 2.32602 | 1.08392 | | -1.1016 | -2.09895 | 5.00E-05 | 0.00107006 | yes |
| ENSG00000128482 | ENSG00000128482 | RNF112 | chr17:19411124-19417276 | OK | 12.4495 | 5.78819 | | -1.1049 | -1.33621 | 0.0006 | 0.00848922 | yes |
| ENSG00000085224 | ENSG00000085224 | ATRX | chrX:77504877-77786269 | OK | 43.3817 | 20.1407 | | -1.10697 | -1.69009 | 0.00015 | 0.00276892 | yes |
| ENSG00000162931 | ENSG00000162931 | TRIM17 | chr1:228407380-228416861 | OK | 1.44085 | 0.666857 | | -1.11147 | -1.2811 | 0.0001 | 0.0019628 | yes |
| ENSG00000076928 | ENSG00000076928 | ARHGEF1 | chr19:41883160-41930150 | OK | 29.4336 | 13.5349 | | -1.12078 | -2.56064 | 5.00E-05 | 0.00107006 | yes |
| ENSG00000260464 | ENSG00000260464 | AL049796.1 | chr1:93847173-93848939 | OK | 4.57617 | 2.10245 | | -1.12207 | -2.2868 | 5.00E-05 | 0.00107006 | yes |
| ENSG00000124160 | ENSG00000124160 | NCOA5 | chr20:46060984-46089952 | OK | 19.4798 | 8.94093 | | -1.12348 | -3.04457 | 5.00E-05 | 0.00107006 | yes |
| ENSG00000221843 | ENSG00000221843 | C2orf16 | chr2:27537385-27582721 | OK | 1.81771 | 0.834242 | | -1.12359 | -3.3733 | 5.00E-05 | 0.00107006 | yes |
| ENSG00000279608 | ENSG00000279608 | AL353795.3 | chr9:35038624-35045991 | OK | 0.937915 | 0.430456 | | -1.12359 | -1.68999 | 5.00E-05 | 0.00107006 | yes |
| ENSG00000196275 | ENSG00000196275 | GTF2IRD2 | chr7:74796143-74851551 | OK | 2.80435 | 1.28422 | | -1.12677 | -1.47394 | 0.0004 | 0.00614885 | yes |
| ENSG00000143429 | ENSG00000143429 | AC116050.1 | chr2:91589493-91659972 | OK | 3.44112 | 1.57077 | | -1.13141 | -2.00446 | 0.00055 | 0.0079124 | yes |
| ENSG00000105699 | ENSG00000105699 | LSR | chr19:35248329-35267964 | OK | 1.82401 | 0.830979 | | -1.13423 | -1.06527 | 0.0001 | 0.0019628 | yes |
| ENSG00000144619 | ENSG00000144619 | CNTN4 | chr3:2098812-3126613 | OK | 51.7202 | 23.4683 | | -1.14002 | -2.66975 | 5.00E-05 | 0.00107006 | yes |
| ENSG00000112319 | ENSG00000112319 | EYA4 | chr6:133240597-133895553 | OK | 1.08971 | 0.494455 | | -1.14004 | -1.22406 | 5.00E-05 | 0.00107006 | yes |
| ENSG00000159450 | ENSG00000159450 | TCHH | chr1:152106316-152115454 | OK | 0.815336 | 0.369472 | | -1.14193 | -2.12907 | 5.00E-05 | 0.00107006 | yes |
| ENSG00000139718 | ENSG00000139718 | SETD1B | chr12:121804179-121832584 | OK | 13.6043 | 6.13137 | | -1.14978 | -4.78978 | 5.00E-05 | 0.00107006 | yes |
| ENSG00000122691 | ENSG00000122691 | TWIST1 | chr7:19020990-19117672 | OK | 11.1911 | 5.03511 | | -1.15226 | -1.84526 | 5.00E-05 | 0.00107006 | yes |
| ENSG00000197483 | ENSG00000197483 | ZNF628 | chr19:55475982-55484487 | OK | 6.88083 | 3.0954 | | -1.15246 | -1.44989 | 0.0002 | 0.00351582 | yes |
| ENSG00000171044 | ENSG00000171044 | XKR6 | chr8:10896044-11201366 | OK | 10.3713 | 4.65487 | | -1.15578 | -1.64133 | 5.00E-05 | 0.00107006 | yes |
| ENSG00000026508 | ENSG00000026508 | CD44 | chr11:35138869-35232402 | OK | 0.657309 | 0.294232 | | -1.15962 | -0.359229 | 5.00E-05 | 0.00107006 | yes |
| ENSG00000164093 | ENSG00000164093 | PITX2 | chr4:110617422-110642123 | OK | 33.7479 | 15.0955 | | -1.16067 | -3.97278 | 5.00E-05 | 0.00107006 | yes |
| ENSG00000157240 | ENSG00000157240 | FZD1 | chr7:91264363-91271326 | OK | 11.8699 | 5.30712 | | -1.1613 | -4.76135 | 5.00E-05 | 0.00107006 | yes |
| ENSG00000165246 | ENSG00000165246 | NLGN4Y | chrY:14522637-14845650 | OK | 21.0411 | 9.3896 | | -1.16407 | -2.04353 | 5.00E-05 | 0.00107006 | yes |
| ENSG00000171847 | ENSG00000171847 | FAM90A1 | chr12:8221259-8227618 | OK | 1.05541 | 0.470738 | | -1.1648 | -1.47019 | 0.0001 | 0.0019628 | yes |
| ENSG00000146938 | ENSG00000146938 | NLGN4X | chrX:5840636-6228863 | OK | 20.9066 | 9.29465 | | -1.16948 | -3.33409 | 5.00E-05 | 0.00107006 | yes |
| ENSG00000127586 | ENSG00000127586 | CHTF18 | chr16:784973-800737 | OK | 9.35822 | 4.16042 | | -1.1695 | -1.40278 | 0.00015 | 0.00276892 | yes |
| ENSG00000259436 | ENSG00000259436 | AC010247.2 | chr19:42086109-42196585 | OK | 29.2139 | 12.9697 | | -1.17151 | -1.53506 | 0.00015 | 0.00276892 | yes |
| ENSG00000135926 | ENSG00000135926 | TMBIM1 | chr2:218270391-218368099 | OK | 0.578085 | 0.256421 | | -1.17277 | -0.344228 | 0.00035 | 0.00550383 | yes |
| ENSG00000182199 | ENSG00000182199 | SHMT2 | chr12:57229326-57240715 | OK | 56.5491 | 24.8711 | | -1.18503 | -3.49169 | 5.00E-05 | 0.00107006 | yes |
| ENSG00000100154 | ENSG00000100154 | TTC28 | chr22:27851668-28679865 | OK | 28.7839 | 12.6499 | | -1.18613 | -1.58398 | 5.00E-05 | 0.00107006 | yes |
| ENSG00000256043 | ENSG00000256043 | CTSO | chr4:155924117-155953917 | OK | 0.616143 | 0.270004 | | -1.19028 | -1.48 | 0.00055 | 0.0079124 | yes |
| ENSG00000255310 | ENSG00000255310 | AF131215.5 | chr8:10896044-11201366 | OK | 9.92089 | 4.34316 | | -1.19172 | -2.37835 | 5.00E-05 | 0.00107006 | yes |
| ENSG00000150275 | ENSG00000150275 | PCDH15 | chr10:53802770-55627942 | OK | 1.91934 | 0.838104 | | -1.19541 | -1.25759 | 5.00E-05 | 0.00107006 | yes |

| | | | | | | | | | | | | |
|---|---|---|---|---|---|---|---|---|---|---|---|---|
| ENSG00000179168 | ENSG00000179168 | GGN | chr19:38384264-38388082 | OK | 7.11793 | 3.09764 | | -1.20029 | -1.54506 | 5.00E-05 | 0.00107006 | yes |
| ENSG00000184986 | ENSG00000184986 | TMEM121 | chr14:105526602-105530202 | OK | 17.5269 | 7.60536 | | -1.20448 | -2.33184 | 5.00E-05 | 0.00107006 | yes |
| ENSG00000181873 | ENSG00000181873 | IBA57 | chr1:228165814-228182257 | OK | 4.25045 | 1.84122 | | -1.20696 | -1.74641 | 5.00E-05 | 0.00107006 | yes |
| ENSG00000204347 | ENSG00000204347 | BTBD17 | chr17:74356415-74361946 | OK | 5.25318 | 2.26872 | | -1.21131 | -2.56982 | 5.00E-05 | 0.00107006 | yes |
| ENSG00000187323 | ENSG00000187323 | DCC | chr18:52340171-53568283 | OK | 27.1099 | 11.6703 | | -1.21598 | -2.44742 | 5.00E-05 | 0.00107006 | yes |
| ENSG00000181585 | ENSG00000181585 | TMIE | chr3:46701332-46710886 | OK | 2.30976 | 0.990621 | | -1.22134 | -1.97813 | 5.00E-05 | 0.00107006 | yes |
| ENSG00000019582 | ENSG00000019582 | CD74 | chr5:150401636-150412929 | OK | 4.49015 | 1.90838 | | -1.23442 | -1.76995 | 5.00E-05 | 0.00107006 | yes |
| ENSG00000248213 | ENSG00000248213 | CICP16 | chr4:118635969-118638782 | OK | 1.38518 | 0.584126 | | -1.24573 | -1.97173 | 5.00E-05 | 0.00107006 | yes |
| ENSG00000173698 | ENSG00000173698 | ADGRG2 | chrX:18989308-19122637 | OK | 4.6952 | 1.97398 | | -1.25008 | -3.40737 | 5.00E-05 | 0.00107006 | yes |
| ENSG00000176125 | ENSG00000176125 | UFSP1 | chr7:100888722-100889718 | OK | 5.38573 | 2.26043 | | -1.25254 | -2.08509 | 5.00E-05 | 0.00107006 | yes |
| ENSG00000269821 | ENSG00000269821 | KCNQ1OT1 | chr11:2444683-2861568 | OK | 1.04513 | 0.436588 | | -1.25934 | -4.96988 | 5.00E-05 | 0.00107006 | yes |
| ENSG00000082482 | ENSG00000082482 | KCNK2 | chr1:215005774-215237093 | OK | 1.13011 | 0.471612 | | -1.2608 | -0.990256 | 0.0004 | 0.00614885 | yes |
| ENSG00000173599 | ENSG00000173599 | PC | chr11:66848232-66958376 | OK | 26.6038 | 11.1016 | | -1.26086 | -1.97282 | 5.00E-05 | 0.00107006 | yes |
| ENSG00000198569 | ENSG00000198569 | SLC34A3 | chr9:137230756-137236554 | OK | 0.879447 | 0.366816 | | -1.26154 | -1.50571 | 0.0002 | 0.00351582 | yes |
| ENSG00000080224 | ENSG00000080224 | EPHA6 | chr3:96814580-97752460 | OK | 5.25375 | 2.19059 | | -1.26203 | -2.17102 | 5.00E-05 | 0.00107006 | yes |
| ENSG00000099256 | ENSG00000099256 | PRTFDC1 | chr10:24848606-24952604 | OK | 4.15604 | 1.72357 | | -1.26981 | -2.06081 | 5.00E-05 | 0.00107006 | yes |
| ENSG00000139132 | ENSG00000139132 | FGD4 | chr12:32399528-32646050 | OK | 18.8836 | 7.80192 | | -1.27523 | -1.68851 | 5.00E-05 | 0.00107006 | yes |
| ENSG00000165655 | ENSG00000165655 | ZNF503 | chr10:75269818-75411842 | OK | 29.9218 | 12.1613 | | -1.29891 | -5.12001 | 5.00E-05 | 0.00107006 | yes |
| ENSG00000235244 | ENSG00000235244 | DANT2 | chrX:115917270-115969089 | OK | 1.47212 | 0.598201 | | -1.29919 | -1.87288 | 5.00E-05 | 0.00107006 | yes |
| ENSG00000250486 | ENSG00000250486 | FAM218A | chr4:164954445-164977668 | OK | 5.60879 | 2.27439 | | -1.30221 | -2.9047 | 5.00E-05 | 0.00107006 | yes |
| ENSG00000225725 | ENSG00000225725 | FAM66E | chr8:7955013-8008755 | OK | 1.29979 | 0.525529 | | -1.30644 | -1.55331 | 0.0005 | 0.00734648 | yes |
| ENSG00000086730 | ENSG00000086730 | LAT2 | chr7:74199651-74229834 | OK | 1.04309 | 0.418357 | | -1.31805 | -1.11365 | 5.00E-05 | 0.00107006 | yes |
| ENSG00000112139 | ENSG00000112139 | MDGA1 | chr6:37630678-37699306 | OK | 19.1393 | 7.64384 | | -1.32417 | -3.54155 | 5.00E-05 | 0.00107006 | yes |
| ENSG00000166987 | ENSG00000166987 | MBD6 | chr12:57520709-57547331 | OK | 25.5798 | 10.2125 | | -1.32466 | -1.25548 | 0.00015 | 0.00276892 | yes |
| ENSG00000239268 | ENSG00000239268 | AC092691.1 | chr3:117672153-117997592 | OK | 105.179 | 41.78 | | -1.33196 | -4.18599 | 5.00E-05 | 0.00107006 | yes |
| ENSG00000070729 | ENSG00000070729 | CNGB1 | chr16:57882339-57971116 | OK | 17.0811 | 6.75007 | | -1.33943 | -2.47418 | 5.00E-05 | 0.00107006 | yes |
| ENSG00000135625 | ENSG00000135625 | EGR4 | chr2:73290928-73293705 | OK | 0.719995 | 0.283799 | | -1.34312 | -1.59028 | 0.00035 | 0.00550383 | yes |
| ENSG00000178295 | ENSG00000178295 | GEN1 | chr2:17663811-17800242 | OK | 2.54139 | 0.989642 | | -1.36064 | -1.02827 | 5.00E-05 | 0.00107006 | yes |
| ENSG00000170122 | ENSG00000170122 | FOXD4 | chr9:116230-118204 | OK | 2.00952 | 0.772076 | | -1.38003 | -2.05298 | 5.00E-05 | 0.00107006 | yes |
| ENSG00000279853 | ENSG00000279853 | AC004453.2 | chr7:44367139-44369292 | OK | 1.14703 | 0.440371 | | -1.38112 | -1.80967 | 5.00E-05 | 0.00107006 | yes |
| ENSG00000099994 | ENSG00000099994 | SUSD2 | chr22:24181258-24189110 | OK | 0.798251 | 0.305673 | | -1.38485 | -1.81828 | 0.00015 | 0.00276892 | yes |
| ENSG00000188738 | ENSG00000188738 | FSIP2 | chr2:185719873-185833290 | OK | 3.90559 | 1.47746 | | -1.40242 | -1.59904 | 5.00E-05 | 0.00107006 | yes |
| ENSG00000258986 | ENSG00000258986 | TMEM179 | chr14:104474677-104605647 | OK | 92.4244 | 34.9599 | | -1.40257 | -4.72869 | 5.00E-05 | 0.00107006 | yes |
| ENSG00000213599 | ENSG00000213599 | SLX1A-SULT1A3 | chr16:30192933-30204310 | OK | 5.06239 | 1.90375 | | -1.41097 | -2.03154 | 5.00E-05 | 0.00107006 | yes |
| ENSG00000172771 | ENSG00000172771 | EFCAB12 | chr3:129401320-129428651 | OK | 0.687101 | 0.256185 | | -1.42333 | -1.34774 | 0.0005 | 0.00734648 | yes |
| ENSG00000144227 | ENSG00000144227 | NXPH2 | chr2:138670771-138780348 | OK | 5.19641 | 1.93313 | | -1.42658 | -2.33614 | 5.00E-05 | 0.00107006 | yes |
| ENSG00000055332 | ENSG00000055332 | EIF2AK2 | chr2:37084450-37157065 | OK | 5.0962 | 1.87856 | | -1.4398 | -1.73621 | 5.00E-05 | 0.00107006 | yes |
| ENSG00000181215 | ENSG00000181215 | C4orf50 | chr4:5897372-6018507 | OK | 1.95024 | 0.716008 | | -1.4456 | -3.78411 | 5.00E-05 | 0.00107006 | yes |
| ENSG00000205670 | ENSG00000205670 | SMIM11A | chr21:34375479-34407866 | OK | 3.75446 | 1.37732 | | -1.44674 | -1.88748 | 5.00E-05 | 0.00107006 | yes |
| ENSG00000164031 | ENSG00000164031 | DNAJB14 | chr4:99896247-99946726 | OK | 30.0078 | 11.0072 | | -1.44689 | -1.26687 | 0.0005 | 0.00734648 | yes |
| ENSG00000227028 | ENSG00000227028 | SLC8A1-AS1 | chr2:39786452-40611053 | OK | 10.17 | 3.72711 | | -1.44819 | -0.72301 | 0.00015 | 0.00276892 | yes |
| ENSG00000147432 | ENSG00000147432 | CHRNB3 | chr8:42697375-42737407 | OK | 8.34097 | 3.05259 | | -1.45018 | -2.66994 | 5.00E-05 | 0.00107006 | yes |
| ENSG00000114854 | ENSG00000114854 | TNNC1 | chr3:52451101-52454070 | OK | 2.09453 | 0.765077 | | -1.45295 | -1.29552 | 0.00065 | 0.00905328 | yes |
| ENSG00000277142 | ENSG00000277142 | LINC00235 | chr16:525154-527407 | OK | 1.73186 | 0.632567 | | -1.45303 | -2.20291 | 5.00E-05 | 0.00107006 | yes |
| ENSG00000080603 | ENSG00000080603 | SRCAP | chr16:30697706-30776307 | OK | 22.293 | 8.13827 | | -1.4538 | -2.2208 | 5.00E-05 | 0.00107006 | yes |
| ENSG00000181722 | ENSG00000181722 | ZBTB20 | chr3:114314500-115147271 | OK | 4.24206 | 1.5451 | | -1.45707 | -1.0827 | 5.00E-05 | 0.00107006 | yes |

| | | | | | | | | | | | |
|---|---|---|---|---|---|---|---|---|---|---|---|
| ENSG00000119772 | ENSG00000119772 | DNMT3A | chr2:25227854-25342590 | OK | 38.8005 | 13.9742 | -1.4733 | -3.44025 | 5.00E-05 | 0.00107006 | yes |
| ENSG00000224109 | ENSG00000224109 | CENPVL3 | chrX:51618054-51618918 | OK | 7.72267 | 2.73375 | -1.49822 | -2.54525 | 5.00E-05 | 0.00107006 | yes |
| ENSG00000159674 | ENSG00000159674 | SPON2 | chr4:1166931-1208962 | OK | 3.39165 | 1.19031 | -1.51066 | -1.47246 | 5.00E-05 | 0.00107006 | yes |
| ENSG00000215154 | ENSG00000215154 | AC141586.1 | chr16:2603349-2643296 | OK | 5.52257 | 1.91986 | -1.52434 | -1.7082 | 5.00E-05 | 0.00107006 | yes |
| ENSG00000280134 | ENSG00000280134 | AL731532.2 | chr10:85555483-85557436 | OK | 1.00523 | 0.347206 | -1.53366 | -1.81076 | 0.0001 | 0.0019628 | yes |
| ENSG00000153071 | ENSG00000153071 | DAB2 | chr5:39284261-39462300 | OK | 2.61877 | 0.900445 | -1.54018 | -1.63052 | 5.00E-05 | 0.00107006 | yes |
| ENSG00000011590 | ENSG00000011590 | ZBTB32 | chr19:35704526-35717038 | OK | 0.715282 | 0.244543 | -1.54843 | -0.871467 | 0.00055 | 0.0079124 | yes |
| ENSG00000142700 | ENSG00000142700 | DMRTA2 | chr1:50417549-50423500 | OK | 59.9644 | 20.3129 | -1.56171 | -6.96832 | 5.00E-05 | 0.00107006 | yes |
| ENSG00000080947 | ENSG00000080947 | CROCCP3 | chr1:16460947-16499257 | OK | 9.3464 | 3.06328 | -1.60933 | -2.25926 | 5.00E-05 | 0.00107006 | yes |
| ENSG00000278249 | ENSG00000278249 | SCARNA2 | chr1:109100192-109100619 | OK | 16.9306 | 5.51809 | -1.61739 | -2.15855 | 5.00E-05 | 0.00107006 | yes |
| ENSG00000110887 | ENSG00000110887 | DAO | chr12:108858931-108901043 | OK | 0.946414 | 0.307882 | -1.62009 | -1.32105 | 5.00E-05 | 0.00107006 | yes |
| ENSG00000253366 | ENSG00000253366 | AC139272.1 | chr5:70751183-70795914 | OK | 2.93477 | 0.944437 | -1.63572 | -1.60439 | 0.0004 | 0.00614885 | yes |
| ENSG00000109063 | ENSG00000109063 | MYH3 | chr17:10628525-10657309 | OK | 0.653783 | 0.209231 | -1.64372 | -0.868355 | 0.0002 | 0.00351582 | yes |
| ENSG00000158125 | ENSG00000158125 | XDH | chr2:31334320-31414715 | OK | 1.37834 | 0.439906 | -1.64766 | -1.18126 | 0.00055 | 0.0079124 | yes |
| ENSG00000172817 | ENSG00000172817 | CYP7B1 | chr8:64587762-64798761 | OK | 0.911231 | 0.290795 | -1.64782 | -1.61284 | 5.00E-05 | 0.00107006 | yes |
| ENSG00000148143 | ENSG00000148143 | ZNF462 | chr9:106863096-107102988 | OK | 18.3992 | 5.85052 | -1.65301 | -2.60651 | 5.00E-05 | 0.00107006 | yes |
| ENSG00000175745 | ENSG00000175745 | NR2F1 | chr5:93409358-93594615 | OK | 86.6647 | 27.3827 | -1.66218 | -5.61881 | 5.00E-05 | 0.00107006 | yes |
| ENSG00000260727 | ENSG00000260727 | SLC7A5P1 | chr16:29613103-29613640 | OK | 3.41156 | 1.07337 | -1.66828 | -1.54044 | 0.0003 | 0.00488215 | yes |
| ENSG00000186867 | ENSG00000186867 | QRFPR | chr4:121329311-121381059 | OK | 2.57302 | 0.809388 | -1.66856 | -2.42419 | 5.00E-05 | 0.00107006 | yes |
| ENSG00000282936 | ENSG00000282936 | AC004706.4 | chr17:6578147-6651634 | OK | 5.78485 | 1.81549 | -1.67193 | -2.30993 | 5.00E-05 | 0.00107006 | yes |
| ENSG00000232599 | ENSG00000232599 | AL008707.1 | chrX:125203804-125204338 | OK | 46.8613 | 14.6334 | -1.67913 | -3.83813 | 5.00E-05 | 0.00107006 | yes |
| ENSG00000234945 | ENSG00000234945 | GTF3C2-AS1 | chr2:27325848-27370486 | OK | 0.996998 | 0.310217 | -1.68431 | -0.405778 | 0.00025 | 0.00421621 | yes |
| ENSG00000146858 | ENSG00000146858 | ZC3HAV1L | chr7:139025705-139036029 | OK | 2.05597 | 0.631482 | -1.70301 | -2.40606 | 5.00E-05 | 0.00107006 | yes |
| ENSG00000284700 | ENSG00000284700 | AL049637.2 | chr1:50423608-50425316 | OK | 5.37327 | 1.64055 | -1.71162 | -2.47092 | 5.00E-05 | 0.00107006 | yes |
| ENSG00000117983 | ENSG00000117983 | MUC5B | chr11:1223065-1262172 | OK | 0.8443 | 0.255494 | -1.72446 | -0.846707 | 0.0003 | 0.00488215 | yes |
| ENSG00000188707 | ENSG00000188707 | ZBED6CL | chr7:150243915-150338150 | OK | 1.93231 | 0.577353 | -1.7428 | -1.78845 | 0.00025 | 0.00421621 | yes |
| ENSG00000277027 | ENSG00000277027 | RMRP | chr9:35657750-35658018 | OK | 53.5785 | 15.8389 | -1.75818 | -1.78992 | 0.0002 | 0.00351582 | yes |
| ENSG00000175920 | ENSG00000175920 | DOK7 | chr4:3463310-3494483 | OK | 0.71593 | 0.211046 | -1.76226 | -0.984594 | 5.00E-05 | 0.00107006 | yes |
| ENSG00000215156 | ENSG00000215156 | AC138409.1 | chr5:34164697-34244796 | OK | 2.76927 | 0.813697 | -1.76694 | -2.77338 | 5.00E-05 | 0.00107006 | yes |
| ENSG00000180422 | ENSG00000180422 | LINC00304 | chr16:89159145-89164245 | OK | 2.64424 | 0.765869 | -1.78769 | -2.27808 | 5.00E-05 | 0.00107006 | yes |
| ENSG00000284610 | ENSG00000284610 | AC107918.4 | chr8:12006187-12015193 | OK | 3.53184 | 1.01522 | -1.79863 | -3.26081 | 5.00E-05 | 0.00107006 | yes |
| ENSG00000275395 | ENSG00000275395 | FCGBP | chr19:39863322-39906323 | OK | 0.672428 | 0.189874 | -1.82434 | -2.13418 | 5.00E-05 | 0.00107006 | yes |
| ENSG00000114013 | ENSG00000114013 | CD86 | chr3:122055365-122121139 | OK | 0.87039 | 0.240285 | -1.85691 | -1.19231 | 5.00E-05 | 0.00107006 | yes |
| ENSG00000178394 | ENSG00000178394 | HTR1A | chr5:63957892-63981043 | OK | 1.7432 | 0.47932 | -1.86268 | -1.46756 | 0.00015 | 0.00276892 | yes |
| ENSG00000064270 | ENSG00000064270 | ATP2C2 | chr16:84368526-84467361 | OK | 1.49709 | 0.408416 | -1.87405 | -0.994481 | 5.00E-05 | 0.00107006 | yes |
| ENSG00000284395 | ENSG00000284395 | AL032819.3 | chr16:1431034-1433397 | OK | 1.71291 | 0.463347 | -1.88629 | -1.7624 | 0.00025 | 0.00421621 | yes |
| ENSG00000102575 | ENSG00000102575 | ACP5 | chr19:11559373-11619135 | OK | 0.866373 | 0.22956 | -1.91612 | -0.538358 | 0.0007 | 0.00960252 | yes |
| ENSG00000180543 | ENSG00000180543 | TSPYL5 | chr8:97273473-97277964 | OK | 6.4484 | 1.64761 | -1.96856 | -5.43398 | 5.00E-05 | 0.00107006 | yes |
| ENSG00000084453 | ENSG00000084453 | SLCO1A2 | chr12:21264599-21419594 | OK | 1.38795 | 0.35197 | -1.97943 | -1.04728 | 5.00E-05 | 0.00107006 | yes |
| ENSG00000267322 | ENSG00000267322 | SNHG22 | chr18:49782166-50195093 | OK | 19.6669 | 4.91618 | -2.00016 | -1.73408 | 0.00035 | 0.00550383 | yes |
| ENSG00000182578 | ENSG00000182578 | CSF1R | chr5:150053290-150113372 | OK | 0.830789 | 0.206719 | -2.00681 | -0.714776 | 0.00055 | 0.0079124 | yes |
| ENSG00000122952 | ENSG00000122952 | ZWINT | chr10:56357227-56361275 | OK | 1.03593 | 0.251158 | -2.04426 | -1.39258 | 5.00E-05 | 0.00107006 | yes |
| ENSG00000258186 | ENSG00000258186 | SLC7A5P2 | chr16:21402236-21520444 | OK | 21.4948 | 5.15065 | -2.06116 | -2.24449 | 5.00E-05 | 0.00107006 | yes |
| ENSG00000118785 | ENSG00000118785 | SPP1 | chr4:87975649-87983426 | OK | 1.95203 | 0.463611 | -2.07399 | -1.52238 | 5.00E-05 | 0.00107006 | yes |
| ENSG00000215630 | ENSG00000215630 | GUSBP9 | chr5:71197645-71208130 | OK | 3.76302 | 0.870221 | -2.11244 | -1.8877 | 0.00065 | 0.00905328 | yes |
| ENSG00000253537 | ENSG00000253537 | PCDHGA7 | chr5:141330570-141512981 | OK | 9.8274 | 2.26728 | -2.11584 | -1.69605 | 0.0005 | 0.00734648 | yes |

| | | | | | | | | | | | |
|---|---|---|---|---|---|---|---|---|---|---|---|
| ENSG00000245468 | ENSG00000245468 | LINC02447 | chr4:7094570-7103385 | OK | 0.63281 | 0.145157 | -2.12416 | -0.957658 | 0.0005 | 0.00734648 | yes |
| ENSG00000042493 | ENSG00000042493 | CAPG | chr2:85394747-85418432 | OK | 0.859008 | 0.181142 | -2.24555 | -0.670782 | 5.00E-05 | 0.00107006 | yes |
| ENSG00000278771 | ENSG00000278771 | RN7SL3 | chr14:49853615-49853914 | OK | 40.0112 | 8.42426 | -2.24778 | -2.53327 | 5.00E-05 | 0.00107006 | yes |
| ENSG00000120738 | ENSG00000120738 | EGR1 | chr5:138465489-138469315 | OK | 4.36579 | 0.912544 | -2.25828 | -4.54728 | 5.00E-05 | 0.00107006 | yes |
| ENSG00000198125 | ENSG00000198125 | MB | chr22:35606763-35637951 | OK | 2.43205 | 0.505265 | -2.26706 | -1.34082 | 5.00E-05 | 0.00107006 | yes |
| ENSG00000185760 | ENSG00000185760 | KCNQ5 | chr6:72621791-73198851 | OK | 0.675678 | 0.139945 | -2.27148 | -0.722551 | 5.00E-05 | 0.00107006 | yes |
| ENSG00000175084 | ENSG00000175084 | DES | chr2:219418376-219426739 | OK | 1.23882 | 0.253088 | -2.29125 | -1.23504 | 0.0002 | 0.00351582 | yes |
| ENSG00000231473 | ENSG00000231473 | LINC00441 | chr13:48296512-48303661 | OK | 9.10271 | 1.7187 | -2.40498 | -5.98076 | 0.00015 | 0.00276892 | yes |
| ENSG00000184956 | ENSG00000184956 | MUC6 | chr11:1012820-1036706 | OK | 1.57235 | 0.271529 | -2.53374 | -1.9625 | 5.00E-05 | 0.00107006 | yes |
| ENSG00000230333 | ENSG00000230333 | AC004160.1 | chr7:11180901-11832198 | OK | 1.36568 | 0.227533 | -2.58547 | -0.807848 | 5.00E-05 | 0.00107006 | yes |
| ENSG00000276168 | ENSG00000276168 | RN7SL1 | chr14:49570983-49614672 | OK | 4380.19 | 715.012 | -2.61495 | -2.5034 | 5.00E-05 | 0.00107006 | yes |
| ENSG00000163046 | ENSG00000163046 | ANKRD30BL | chr2:132147590-132257969 | OK | 2.16164 | 0.295958 | -2.86866 | -2.73285 | 5.00E-05 | 0.00107006 | yes |
| ENSG00000268119 | ENSG00000268119 | AC010615.2 | chr19:21397118-21569237 | OK | 2.0258 | 0.270529 | -2.90464 | -1.33884 | 5.00E-05 | 0.00107006 | yes |
| ENSG00000125730 | ENSG00000125730 | C3 | chr19:6677703-6737603 | OK | 2.34916 | 0.302311 | -2.95804 | -0.952681 | 5.00E-05 | 0.00107006 | yes |
| ENSG00000259129 | ENSG00000259129 | LINC00648 | chr14:47764953-47795092 | OK | 1.17155 | 0.148327 | -2.98157 | -0.944733 | 5.00E-05 | 0.00107006 | yes |
| ENSG00000223804 | ENSG00000223804 | AC244669.1 | chr1:120197084-120341871 | OK | 30.0222 | 3.71105 | -3.01613 | -14.5532 | 0.0005 | 0.00734648 | yes |
| ENSG00000100181 | ENSG00000100181 | TPTEP1 | chr22:16601886-16704477 | OK | 2.19977 | 0.250619 | -3.13378 | -1.60997 | 5.00E-05 | 0.00107006 | yes |
| ENSG00000205609 | ENSG00000205609 | EIF3CL | chr16:28379578-28403879 | OK | 3.26655 | 0.360151 | -3.18109 | -4.71405 | 5.00E-05 | 0.00107006 | yes |
| ENSG00000110680 | ENSG00000110680 | CALCA | chr11:14904996-15082342 | OK | 1.05636 | 0.10052 | -3.39355 | -1.03955 | 0.0001 | 0.0019628 | yes |
| ENSG00000274012 | ENSG00000274012 | RN7SL2 | chr14:49861175-49864379 | OK | 15342.4 | 1435.04 | -3.41836 | -16.3456 | 5.00E-05 | 0.00107006 | yes |
| ENSG00000125414 | ENSG00000125414 | MYH2 | chr17:10383131-10623886 | OK | 0.696698 | 0.0593993 | -3.55202 | -1.02143 | 5.00E-05 | 0.00107006 | yes |
| ENSG00000143632 | ENSG00000143632 | ACTA1 | chr1:229431244-229434098 | OK | 5.29051 | 0.436575 | -3.5991 | -4.04537 | 5.00E-05 | 0.00107006 | yes |
| ENSG00000149256 | ENSG00000149256 | TENM4 | chr11:78652830-79440948 | OK | 125.725 | 10.3637 | -3.60067 | -8.21827 | 5.00E-05 | 0.00107006 | yes |
| ENSG00000230021 | ENSG00000230021 | AL669831.3 | chr1:586070-859446 | OK | 8.34561 | 0.252916 | -5.04429 | -0.979091 | 5.00E-05 | 0.00107006 | yes |
| ENSG00000104879 | ENSG00000104879 | CKM | chr19:45306413-45322977 | OK | 2.93067 | 0.08861 | -5.04762 | -3.54927 | 5.00E-05 | 0.00107006 | yes |
| ENSG00000166428 | ENSG00000166428 | PLD4 | chr14:104924815-104978357 | OK | 0.610942 | 0.0183835 | -5.05455 | -0.370878 | 5.00E-05 | 0.00107006 | yes |
| ENSG00000106153 | ENSG00000106153 | CHCHD2 | chr7:56101568-56106576 | OK | 68.8106 | 1.61703 | -5.41121 | -8.97752 | 5.00E-05 | 0.00107006 | yes |
| ENSG00000136352 | ENSG00000136352 | NKX2-1 | chr14:36320752-37173811 | OK | 0.702974 | 0.0152427 | -5.52728 | -0.545366 | 5.00E-05 | 0.00107006 | yes |

| Names | total | elements | Names | total | elements | Names | total | elements | Names | total | elements | Names | total | elements | Names | total | elements | Names | total | elements |
|---|---|---|---|---|---|---|---|---|---|---|---|---|---|---|---|---|---|---|---|---|
| AHN_GBA AHN_LRRK2 AHN_SNCA | 50 | AC092691.1 | AHN_GBA [illegible] | 81 | CHPT1 | AHN_GBA AHN_SNCA | | TRIM68 | AHN_LRRK2 AHN_SNCA | | AL669831.[illegible] | AHN_GBA | | B3GNT2 | AHN_LRRK2 | | DHRS2 | AHN_SNCA | | TMSB4Y |
| | | CNTN3 | | | FP236383.3 | | 30 | EIF2AK2 | | 84 | RIT2 | | 136 | ANKRD45 | | 235 | TSSC2 | | 277 | MLC1 |
| | | AL008707.1 | | | TPTE2P1 | | | ASB4 | | | CD44 | | | SLC18A1 | | | CLMP | | | AC010247.2 |
| | | CNN2 | | | CYBA | | | FOXP2 | | | ADRA2C | | | PTN | | | ANXA2 | | | XDH |
| | | SLC8A1-AS1 | | | COL4A5 | | | AC107918.4 | | | FOXD2 | | | RN7SK | | | BX324167.2 | | | APOL2 |
| | | HTR1A | | | ADA2 | | | CALB1 | | | SRCAP | | | COL3A1 | | | RWDD2B | | | ATRX |
| | | LINC00441 | | | UNC5B | | | EFL1 | | | DEPDC7 | | | LINC01310 | | | LYPD6B | | | NCOA5 |
| | | AC004706.4 | | | FAM66D | | | NKAIN4 | | | LRRC32 | | | CXCR4 | | | ZAP70 | | | ANKRD34B |
| | | TCHH | | | LRRC37A4P | | | MB | | | LAYN | | | PNMA6B | | | GLIS1 | | | DIO3 |
| | | MUC6 | | | GSTM1 | | | ZNF385B | | | LEFTY2 | | | IRAK1 | | | CMTM8 | | | ZPLD1 |
| | | STXBP6 | | | KIAA1683 | | | AL049637.2 | | | CDH13 | | | CXCL14 | | | DIRAS3 | | | EGR4 |
| | | C5orf38 | | | COL1A1 | | | CRYAB | | | TCIM | | | C1QTNF4 | | | SALL1 | | | RMRP |
| | | C3 | | | MMP14 | | | ADGRG2 | | | CRH | | | AC092117.1 | | | ZNF572 | | | FAM238C |
| | | PLCD1 | | | SNHG5 | | | CD74 | | | TENM4 | | | TMC4 | | | SIGIRR | | | MDGA1 |
| | | P2RX3 | | | CPXM1 | | | ISM2 | | | DANT2 | | | GDF15 | | | CD37 | | | KCNK2 |
| | | CERS6-AS1 | | | AL031282.1 | | | CALCA | | | MBD6 | | | TNC | | | GP1BA | | | HSPB8 |
| | | PGGHG | | | GSTT2B | | | EGR1 | | | C1QL2 | | | NAMPTP1 | | | AC138969.2 | | | LINC00235 |
| | | LINC00304 | | | LINC01971 | | | HLA-B | | | COL4A4 | | | CRABP2 | | | RPS2P55 | | | CNGB1 |
| | | EPHA6 | | | FAM182B | | | SHANK1 | | | MUC3A | | | AP006621.5 | | | SLC12A7 | | | ID3 |
| | | FGD4 | | | HERC2P3 | | | CAPG | | | TTC28 | | | URB1-AS1 | | | SIDT1 | | | CENPQ |
| | | PENK | | | GNAQ | | | GFRA1 | | | AC141586.1 | | | CCDC125 | | | SYNDIG1 | | | INO80B |
| | | VIP | | | RRM2 | | | EBF1 | | | CLEC2L | | | SYNE3 | | | TC2N | | | NLGN4X |
| | | WASH5P | | | TUBBP5 | | | LRRC61 | | | SOHLH2 | | | HERC2P2 | | | GZMM | | | PLCB2 |
| | | FCGBP | | | C11orf80 | | | NSFP1 | | | RASD2 | | | ZFP3 | | | TICAM1 | | | LATS2 |
| | | PRKCD | | | SERF1A | | | PVT1 | | | ID1 | | | MYOM2 | | | FUT1 | | | TMBIM1 |
| | | MYH2 | | | FAM156A | | | CSF1R | | | STAT4 | | | LHFPL2 | | | SFRP5 | | | TMEM51 |
| | | SPON2 | | | LINC00116 | | | TRIP10 | | | SHISA9 | | | TMSB15B | | | UCN | | | COMMD4 |
| | | TCERG1L | | | AC026403.1 | | | RGPD6 | | | FAM90A1 | | | AC068446.1 | | | AC004656.1 | | | DAB2 |
| | | STAC2 | | | FAM239B | | | ONECUT3 | | | TRH | | | LINC01979 | | | LDHD | | | UBE2S |
| | | MUC5B | | | WNT3 | | | CACNA2D3 | | | KCNQ5 | | | GRM8 | | | IFITM3 | | | ATP2C2 |
| | | CHRD | | | FAM24B | | | | | | ZBTB20 | | | TIMP3 | | | FABP7 | | | C8orf58 |
| | | NHLH2 | | | AC007614.4 | | | | | | CBLN4 | | | KC6 | | | LRP6 | | | PNRC2 |
| | | ZNF462 | | | AC092683.1 | | | | | | FAM218A | | | RASGRP1 | | | SH2D3A | | | PPM1J |
| | | ANKRD30BL | | | TXNRD2 | | | | | | LINC00943 | | | AC006453.2 | | | PIR | | | SUSD2 |
| | | TMIE | | | FABP3 | | | | | | EN2 | | | AC107027.3 | | | KCNQ4 | | | NR2F1 |
| | | AL355032.1 | | | FAM66A | | | | | | CHRNB4 | | | SLC16A7 | | | BX890604.1 | | | AL591848.4 |
| | | ZC3HAV1L | | | COL6A2 | | | | | | FSIP2 | | | GUCY1A2 | | | BCL2L11 | | | TTR |
| | | DNMT3A | | | MEG3 | | | | | | NPPC | | | LINC00623 | | | DGKG | | | GJA1 |
| | | ACTA1 | | | RPS2P46 | | | | | | PRPH | | | MEG9 | | | ZNF717 | | | PTGDS |
| | | IGFBP3 | | | IGSF9B | | | | | | S100A10 | | | TMEM255A | | | GRASP | | | RNF31 |
| | | LINC00648 | | | FKBP1C | | | | | | ARID5A | | | SLC18A2 | | | SMIM3 | | | UFSP1 |
| | | AC004160.1 | | | PDPR | | | | | | ARHGEF1 | | | SERPING1 | | | PTPRZ1 | | | BCAN |
| | | NPNT | | | PCDHGB6 | | | | | | AC079949.2 | | | AC092368.3 | | | AC092835.1 | | | GEN1 |
| | | LINC00622 | | | GRP | | | | | | KCNB2 | | | QPRT | | | CLDN4 | | | CTNNA1 |
| | | CKM | | | FAM66B | | | | | | PITX2 | | | CACNG3 | | | POU3F1 | | | RTL1 |
| | | CD86 | | | ARHGAP25 | | | | | | SIM1 | | | AC139749.1 | | | ENTPD3 | | | AC005052.1 |
| | | RN7SL2 | | | NES | | | | | | SLC34A3 | | | BCAM | | | AC215219.1 | | | C5AR1 |
| | | PSTPIP2 | | | RNF224 | | | | | | RAMP1 | | | PLEKHG4B | | | ARHGAP30 | | | SLC22A18 |
| | | C4orf50 | | | AC007614.5 | | | | | | TPTEP1 | | | EN1 | | | CDK11A | | | DNAJC22 |
| | | AC138409.1 | | | SIGLEC1 | | | | | | ALS2CL | | | CU633906.1 | | | COL9A2 | | | PXDC1 |
| | | | | | S100B | | | | | | PRKCQ | | | NBPF8 | | | ANXA3 | | | ACP5 |
| | | | | | CCDC144NL-AS1 | | | | | | KIRREL3 | | | DNAH9 | | | AGGF1P1 | | | RELT |
| | | | | | SPARC | | | | | | QRFPR | | | RNASEL | | | UAP1L1 | | | AL032819.3 |
| | | | | | PCDHB5 | | | | | | SIM2 | | | LINC00649 | | | TMEM52 | | | GPC5 |
| | | | | | HLA-C | | | | | | PCP4 | | | SNHG17 | | | PLXDC2 | | | GRIK1 |
| | | | | | RPS2P5 | | | | | | GRIN2B | | | IL6ST | | | CTDSPL2 | | | KCNJ4 |
| | | | | | U2AF1 | | | | | | CICP16 | | | ETS1 | | | ADGRG3 | | | NFIX |
| | | | | | NPIPA5 | | | | | | SCARNA2 | | | GRIN2C | | | POTEF | | | CDH9 |
| | | | | | FP565324.1 | | | | | | PLIN3 | | | SLC10A4 | | | SLC2A13 | | | SEM1 |
| | | | | | AATBC | | | | | | POMK | | | MEG8 | | | AC236972.3 | | | TRIM7 |
| | | | | | RBPMS2 | | | | | | EYA4 | | | NBPF26 | | | FYCO1 | | | AF131215.5 |
| | | | | | TFPI2 | | | | | | REPS2 | | | TRIM65 | | | ZFP36L1 | | | LGI2 |
| | | | | | AC068446.2 | | | | | | SLC7A5P2 | | | PKIB | | | TSPO | | | STARD5 |
| | | | | | BX664615.2 | | | | | | BAG3 | | | CR769775.4 | | | ANKRD36C | | | BTN2A2 |
| | | | | | AC087632.1 | | | | | | TULP4 | | | TUSC1 | | | CABP1 | | | FAM66E |

| | | | | |
|---|---|---|---|---|
| DHRS4-AS1 | SLCO1A2 | CBSL | PURA | CCNE1 |
| KLF9 | FOXD4 | FAM50B | LINC00969 | AL731532.2 |
| MGMT | MCAM | AC135983.2 | SEMA3E | CNTN4 |
| COL1A2 | FZD1 | TTTY14 | PIRT | LINC02525 |
| AC012414.5 | AC245060.5 | MLLT10 | SPOCK2 | AL138831.3 |
| TMF1 | DCC | C2CD2 | PHC3 | TRIM17 |
| TXNIP | EIF3CL | AL513318.2 | FNBP1P1 | AL139424.2 |
| AC015961.1 | CDH11 | APCDD1 | AP000769.1 | AL353795.3 |
| AF186192.2 | DNAJB14 | MKRN3 | A4GALT | BTBD17 |
| RPL9P9 | SLC17A7 | AL662844.4 | ULBP2 | PLD4 |
| CDK10 | TESC | HCG11 | CA10 | GTF3C2-AS1 |
| AL512625.3 | FOXD2-AS1 | LEKR1 | AC233280.1 | PRTFDC1 |
| ZP3 | CRISPLD2 | FAR2P2 | HSPA12B | AC026585.1 |
| EFS | NUTM2F | SVIL | AC025884.2 | FUT3 |
| SEL1L | C2orf16 | NGFR | ZNF460 | AGPAT2 |
| | SULT1C4 | FKBP10 | CRHBP | GTF2IRD2 |
| | PRKCQ-AS1 | MYO1F | ASS1 | SLC38A3 |
| | DIO3OS | DRAIC | AC025884.1 | BCHE |
| | C1QL4 | ZNF649 | TOP2A | PCDHGA7 |
| | | ISLR2 | GPR50 | LINC01137 |
| | | TPM2 | EDNRA | GRM5 |
| | | AL138756.1 | IGF1R | CD36 |
| | | EPHA7 | PKD1P5 | AC010343.1 |
| | | AC145124.1 | CNOT6LP1 | NKD2 |
| | | GPC3 | APOO | HPCA |
| | | CU639417.1 | EGFL6 | ANKRD36 |
| | | SMN2 | REC8 | EHD4 |
| | | RPL29P11 | BDH2 | DAO |
| | | LRRIQ3 | ADK | NDST4 |
| | | KCNC3 | POTEE | AC105383.1 |
| | | CBS | FAM239A | RGS4 |
| | | MRC2 | PI4KAP1 | RSPH4A |
| | | TMEM135 | CGA | CHRNB3 |
| | | PWRN1 | AC011487.1 | RN7SL1 |
| | | CDH12 | AC092535.1 | ABHD3 |
| | | SMIM32 | PDZRN4 | VSNL1 |
| | | U2AF1L5 | CARTPT | EBF2 |
| | | TAC1 | PAPPA | CROCCP3 |
| | | FKBP5 | AC097639.1 | FUT6 |
| | | GPR157 | PAX8 | SPARCL1 |
| | | COL5A1 | HRAT92 | ZNF467 |
| | | ZNF681 | AC068775.1 | ANK1 |
| | | CTSZ | HLA-DPB2 | KCNJ3 |
| | | VAMP7 | TNFRSF14 | HABP4 |
| | | NR2F2-AS1 | POMZP3 | PLD1 |
| | | POU3F3 | PTPN6 | AP000894.2 |
| | | PMP22 | NPIPB13 | PC |
| | | OR2L13 | LITAF | SMIM11A |
| | | MFAP3L | GNG11 | GSDMB |
| | | LPAR1 | SERTM1 | CENPVL3 |
| | | FHAD1 | RSPO2 | SAPCD2 |
| | | TES | PCDHA11 | RASGRP3 |
| | | SCART1 | FAM189A2 | TBX19 |
| | | PLS3 | EGFL7 | AC048341.2 |
| | | AC092070.2 | FABP6 | TNNC1 |
| | | PUS7L | ZNF805 | HTRA1 |
| | | DHRS3 | SLC32A1 | GALNT14 |
| | | SLC11A1 | ATP2A1 | HSPA4L |
| | | TNR | PAX8-AS1 | ID2 |
| | | CHODL | CYP2A6 | ADRA1A |
| | | MIR4458HG | GLB1L3 | OTX1 |
| | | ADAMTS17 | RASSF4 | AC005332.5 |
| | | UPK3BL1 | AL133216.2 | KISS1R |
| | | IGFBP5 | EYA1 | SETD1B |
| | | CTGF | IL13RA2 | NABP1 |
| | | H1F0 | NENF | CHRNA2 |

| | | |
|---|---|---|
| NR2F2 | TRPV2 | CICP14 |
| AC139495.3 | TGIF1 | TMEM121 |
| TMEM100 | TP53 | AF131215.6 |
| AC068282.1 | SYTL4 | PRDM12 |
| DPY19L2P3 | PCDHGA11 | ASAP3 |
| | CCDC163 | RPL24P4 |
| | FAM162A | LANCL2 |
| | AC104083.1 | AC004453.2 |
| | CDH23 | GGN |
| | INF2 | LMO3 |
| | EFCAB2 | SOCS2 |
| | NEDD4L | XKR6 |
| | WWC1 | Z97653.2 |
| | RPL7P1 | KCNQ1OT1 |
| | CFC1B | HRH2 |
| | KLK10 | ZNF594 |
| | AC074029.4 | AL035563.1 |
| | AC008764.4 | KIAA1217 |
| | FHL2 | SNTG2 |
| | TCF7L2 | ZNF628 |
| | FAM157B | SLX1A-SULT1A3 |
| | FAM221A | SULF1 |
| | AK5 | GAS2L3 |
| | HLA-A | SLC7A5P1 |
| | GTF2H2C | TMEM179 |
| | CNTNAP3B | CEP83-AS1 |
| | RSPO4 | MYH3 |
| | TNFRSF1A | SNHG22 |
| | CU638689.4 | VEGFC |
| | FAM19A1 | LSR |
| | FAR2P1 | HS3ST2 |
| | AGAP7P | SPSB1 |
| | KALRN | DDX23 |
| | KRT10 | LPCAT2 |
| | SVIL-AS1 | DES |
| | PLA2G16 | RNF112 |
| | FRG1BP | ZNF221 |
| | TCF19 | MAGIX |
| | MLXIPL | ITPR1-AS1 |
| | LINC02175 | DNAJB4 |
| | PCAT7 | KIT |
| | AC021054.1 | KCNB1 |
| | FAM90A25P | CSRP2 |
| | KCNK1 | MAN1C1 |
| | SV2C | NRXN3 |
| | MUC20P1 | PTPN3 |
| | CILP2 | GUSBP9 |
| | KC877982.1 | MAF |
| | DUX4L50 | C1orf115 |
| | EPAS1 | MCM6 |
| | CD9 | FRZB |
| | CDHR3 | MPP1 |
| | PLPP2 | RHPN2 |
| | CPNE9 | SPP1 |
| | ANKRD18A | AC007405.3 |
| | WFDC1 | DGCR5 |
| | RASAL2-AS1 | CYP7B1 |
| | RNF175 | NMU |
| | TMSB4X | ZBED6CL |
| | ARHGEF19 | TFEB |
| | PSCA | STK3 |
| | SALL3 | TCFL5 |
| | MAST4 | AL078639.1 |
| | HAS1 | GALNT12 |
| | GATAD2B | AC010615.2 |
| | WASH6P | ARPIN |

| | |
|---|---|
| AC016026.1 | EIF4E1B |
| GSTM2 | SMCO4 |
| KAT6A | CHTF18 |
| GRIA3 | C12orf60 |
| MT2A | NLGN4Y |
| SERPINF2 | PAPPA2 |
| FLVCR1-AS1 | PCDH15 |
| PCDHGA6 | LAT2 |
| C7 | CTSO |
| AC242376.2 | GPRIN2 |
| CDKN3 | MAN2A1 |
| AL355355.1 | HMCN2 |
| GYG2 | NINJ1 |
| NOMO3 | IBA57 |
| CEBPZOS | PLCD4 |
| SERPINA5 | FAM122B |
| SSTR5 | TSPYL5 |
| SIK1 | AC116050.1 |
| PHF11 | HSD17B7 |
| LINC01238 | ANG |
| FILIP1 | ERICH3 |
| AC140912.1 | ZNF804A |
| APOC1 | TPD52L1 |
| AC159540.2 | LINC01158 |
| MED30 | PLEKHH2 |
| TMEM176B | ZWINT |
| SOX2 | RN7SL3 |
| COMT | CCDC40 |
| CCDC8 | LIN28A |
| RPH3A | AHCTF1 |
| VIPR2 | CDC42EP1 |
| MKKS | RELN |
| AGO3 | TFRC |
| NYAP2 | EFCAB12 |
| KRT23 | DOK7 |
| MX2 | NTS |
| DAB1-AS1 | JUN |
| BBX | AC110597.1 |
| | ARAP3 |
| | BACH2 |
| | TCN2 |
| | UGT8 |
| | CHCHD2 |
| | AC109347.1 |
| | B3GLCT |
| | 6-Sep |
| | NKX2-1 |
| | ZBTB32 |
| | TWIST1 |
| | LINC02300 |
| | NTN3 |
| | CRB1 |
| | VAV3 |
| | AC017104.4 |
| | AC244669.1 |
| | DUSP10 |
| | FLJ31104 |
| | SLC35F4 |
| | ZNF503 |
| | RNASE4 |
| | NRTN |
| | AC006378.2 |
| | SHMT2 |
| | AC139272.1 |
| | BTG3 |
| | CHST3 |

AL049796.1
LINC02447
NXPH2
CHAC1
DMRTA2
AC092376.2
C18orf65
ST18
DAAM2
CREB5
EFEMP1
HOPX
ECT2
DOCK6
GRAMD1C
